# Molecular and functional characterization of different BrainSphere models for use in neurotoxicity testing on microelectrode arrays

**DOI:** 10.1101/2023.03.13.532388

**Authors:** Julia Hartmann, Noah Henschel, Kristina Bartmann, Arif Dönmez, Gabriele Brockerhoff, Katharina Koch, Ellen Fritsche

## Abstract

The currently accepted methods for neurotoxicity (NT) testing rely on animal studies. However, high costs and low testing throughput hinder their application for large numbers of chemicals. To overcome these limitations, *in vitro* methods are currently developed which are based on human induced pluripotent stem cells (hiPSC) that allow higher testing throughput at lower costs. We applied six different protocols to generate 3D BrainSphere models for acute NT evaluation. These include three different media for 2D neural induction and two media for subsequent 3D differentiation resulting in self-organized, organotypic neuron/astrocyte microtissues. All induction protocols yielded nearly 100 % nestin-positive hiPSC-derived neural progenitor cells (hiNPCs) yet with different gene expression profiles concerning regional patterning. Moreover, gene expression and immunocytochemistry analyses revealed that the choice of media determines neural differentiation patterns. On the functional level, BrainSpheres exhibited different levels of electrical activity on microelectrode arrays (MEA). Spike sorting allowed BrainSphere functional characterization with the mixed cultures consisting of GABAergic, glutamatergic, dopaminergic, serotonergic, and cholinergic neurons. A test method for acute NT testing, the human multi-neurotransmitter receptor (hMNR) assay, was proposed applying such MEA-based spike sorting. These models are not only promising tools in toxicology but also for drug development and disease modeling.

## 1. Introduction

The currently accepted methods for neurotoxicity (NT) testing rely on animal studies. These are defined in the OECD test guidelines (TG) 418, 419, and 424, [1–3]. The drawbacks of these TG studies are their resource intensities regarding money and time, their high variability, their lack of mechanistic understanding, and their uncertainties due to species differences [4,5]. Such species specificities between humans and rodents decrease confidence in TG neurotoxicity testing as the human brain holds unique features [6,7]. The human uniqueness of the brain is also reflected in the high attrition rates in central nervous system-related drug development when moving from preclinical research to clinical drug applications in humans [8,9]. One of the main arguments for a paradigm change in human health risk assessment, however, is the low testing throughput that has been leading to a large number of untested chemicals in our exposome [10].

There is international consensus that current testing needs cannot be satisfied by animal guideline studies. Hence, regulatory agencies, industry, and academia are currently promoting a change to mechanism-based new approach method (NAM)-based next-generation risk assessment (NGRA) [11–15]. NGRA aims at replacing apical endpoints measured in animals by broad biological coverage NAMs that establish a point-of-departure (POD) based on compounds’ bioactivities, and comparison of those PODs with exposure measures or their predictions e.g. by physiological-based kinetic modeling [16]. Uncertainties can be quantified by probability assessment [17]. NAMs should address both general cellular targets and targets specific to the function of the investigated organ system. Therefore, each method should be carefully examined for the presence or absence of biological processes, to thereby characterize a suitable application domain (fit-for-purpose models) that is best defined by applying model substances or performing case studies [18,19]. Since *in vitro* systems cannot represent an entire organism, they should be contextualized, e.g. in an integrated approach for testing and assessment (IATA) [11].

Since the nervous system is a very complex organ, it can be disrupted via a plethora of MoA involving amongst others neurotransmitter receptors and ion transporters [20]. In general, these MoA affect neuronal function and communication by inhibiting neurotransmitter synthesis or degradation, increasing or preventing neurotransmitter release, blocking neurotransmitter receptors, or interfering with the multiple ion channels [21–25]. Effects on neuronal function can be assessed by recording extracellular local field potentials of cultured neurons on microelectrode arrays (MEAs) thus providing functional readouts for neurotoxicity assessment [26]. Thereby data on spike-, burst- and network synchronicity-related parameters is generated [27]. MEAs were already successfully used for acute neurotoxicity studies with rodent cell cultures and genetically engineered human induced pluripotent stem cell (hiPSC)-derived neurons and human astrocytes [28–35]. However, brain regions, neuronal subtypes, and individual neuronal units were not evaluated in these studies.

Reprogramming of somatic cells into hiPSCs [36,37] opened new ways to generate self-organized human *in vitro* neural networks (NN) by ectodermal and further mixed culture (neuronal and glia) differentiation [38]. By now a plethora of 2D and 3D neural induction and differentiation protocols are available that can be applied fit-for-purpose [39]. One promising way to neurally induce hiPSC in 2D and 3D cultures is dual SMAD inhibition, which induces neuroectodermal differentiation by altering bone morphogenic protein (BMP) and transforming growth factor beta (TGF□)1 signaling pathways [40–42]. Subsequent neural differentiation of hiPSC-derived neural progenitor cells (hiNPC) into neurons and glial cells *in vitro* is performed with different neurotrophic factors and deprivation of growth factors such as EGF [43–50]. Medium composition in the stem cell differentiation process is critical [51]. For example, we recently showed that addition of creatine monohydrate, interferon-γ, neurotrophin-3, dibutyryl-cAMP, and ascorbic acid enhances neuronal electrical activity [52]. 2D protocols for neural induction and differentiation are highly efficient and reproducible due to the even distribution of the provided media supplements, however, they lack formation of a complex three-dimensional morphological architecture and cell-cell-contacts [41,53,54]. Brain organoid cultures overcome this drawback by containing morphological organization, but they require long differentiation periods beginning from 1 up to 9 months and show high variability between specimens [55,56]. Spheroid cultures, such BrainSpheres, allow more complex cytoarchitecture than monolayer cultures and form more cell-cell contacts than 2D cultures. Moreover, they need shorter differentiation times than organoids, are less variable and hence are suitable for medium-throughput testing in neurotoxicity studies, drug development, or disease modeling [53,57–65].

In this study, we compared the molecular and functional consequences of applying six different cell culture medium protocols for BrainSphere generation by gene and protein analyses as well as MEA recordings. In addition, we set up a MEA-based NAM for acute neurotoxicity testing by using spike sorting.

## 2. Materials and Methods

### 2.1. Cultivation of hiPSCs

To guarantee high-quality pluripotent cells, we banked our hiPSCs in a two-tiered process containing a fully characterized Master Cell Bank (MCB) and a partially characterized Working Cell Bank (WCB) as described in Tigges et al., 2021 [66]. The characterization of the cells includes assays regarding cell morphology and identity, pluripotency, karyotype stability, antigen and gene expression, viability, mycoplasma contamination, and post-thaw recovery. In this work, the hiPSC line IMR90 (clone 4, WiCell, USA) was used and cultivated under humidified conditions at 37 °C and 5 % CO_2_. The cells were grown on laminin (5 µg/mL, #LN521, Biolamina, Sweden)-coated 6-well plates in iPS-Brew XF medium (#130-104-368, Miltenyi Biotech, Germany) supplemented with 1 % Penicillin/Streptomycin (#P06-07100, PAN Biotech, Germany). The medium was replaced on 6 out of 7 days per week and on the 7th day the cells received the double amount of medium (“double feed”). The hiPSC colonies were passaged with 0.5 mM EDTA (#15575020, Thermo Fisher Scientific, USA) performing ‘colony-splittin’.

### 2.2. Neural Induction of hiPSCs into human induced neural progenitor cells (hiNPCs)

For each neural induction protocol, hiPSC-colonies were dissolved using the Gentle Cell Dissociation Reagent (#100-0485, Stemcell Technologies, Canada) for 10 min at 37 °C and 5 % CO_2_. After centrifugation (300 g, 10 min), the cell pellet was resuspended in the respective neural induction medium and the cells were seeded with a cell density of 2*10^6^ cells per well and cultivated under humidified conditions at 37 °C and 5 % CO_2_. During the neural induction, the cells were fed on 6 out of 7 days per week, and on the 7th day the cells were fed with twice the amount of medium.

#### 2.2.1. 2D-NIM protocol

The 2D-NIM protocol was adapted from our previously published 3D neural induction protocol (NIM; [52,67]) and modified as followed to achieve a 2D culture. The neural induction medium consists of 2 parts DMEM (high glucose, #31966021, Invitrogen, USA) and 1 part Ham’s F12 Nutrient Mix (#31765027, Invitrogen, USA) supplemented with 1 % Penicillin/Streptomycin (#P06-07100, PAN-Biotech, Germany), 2 % B-27™ supplement (50x, serum-free, #17504044, Invitrogen, USA), 1 % N-2 supplement (100x, #17502048, Thermo Fisher Scientific, USA), 20 ng/mL recombinant human EGF (#PHG0313, Gibco, USA), 20 % Knockout Serum Replacement (#10828028, Thermo Fisher Scientific, USA), 10 µM SB-431542 (#S4317, Sigma-Aldrich, USA), and 0.5 µM LDN-193189 hydrochloride (#SML0559, Sigma-Aldrich, USA). After the above-described singularization, the hiPSCs were transferred to a polyethyleneimine (PEI, 0.1 %; #181978, Sigma-Aldrich, USA)-laminin (15 µg/mL; #LN521, Biolamina, Sweden)-coated 6-well plate and cultivated in 2D-NIM medium supplemented with 10 µM Y-27632 (only for the first 24 h after passaging; #HB2297, Hello Bio, Great Britain). On days 12 and 17, hiNPCs were passaged by enzymatic dissociation with Accutase (#07920, Stemcell Technologies, Canada) for 10 min at 37 °C and 5 % CO_2_ and transferred to a new PEI-laminin-coated 6-well plate. From day 12, the cells were cultivated in neural progenitor medium based on 2D-NIM medium without the dual SMAD inhibitors SB-431542 and LDN-193189 and supplemented with 20 ng/mL recombinant human basic FGF (#233-FB, R&D Systems, Germany) and 10 µM Y-27632 (only for the first 24 h after passaging). On day 21, hiNPCs were singularized with Accutase and frozen in neural progenitor medium containing 10 % dimethyl sulfoxide (DMSO, #A994.1, Carl-Roth, Germany) and 10 µM Y-27632.

#### 2.2.2. GNEIB protocol

The GNEIB neural induction protocol published by Hyvärinen et al., 2019 and Shi et al., 2012 was applied with minor changes [42,44]: The hiPSC-colonies were singularized as described above and resuspended in GNEIB neural induction medium based on 1 part DMEM/F-12 (#31331028, Gibco, USA) and 1 part Neurobasal™ medium (#21103049, Gibco, USA) supplemented with 1 % Penicillin/Streptomycin (#P06-07100, PAN-Biotech, Germany), 1 % B-27™ supplement (50x, serum-free, #17504044, Thermo Fisher Scientific, USA), 0.5 % N-2 supplement (100x, #17502048, Thermo Fisher Scientific, USA), 0.5 mM GlutaMAX™ supplement (#35050061, Thermo Fisher Scientific, USA), 0.5 % MEM non-essential amino acids (100x, #11140050, Thermo Fisher Scientific, USA), 2.5 µg/mL insulin (#I9278, Sigma-Aldrich, USA), 50 µM beta-mercaptoethanol (#31350010, Thermo Fisher Scientific, USA), 10 µM SB-431542 (#S4317, Sigma-Aldrich, USA), and 100 nM LDN-193189 hydrochloride (#SML0559, Sigma-Aldrich, USA). The cells were transferred onto a poly-L-ornithine (PLO, 100 µg/mL; #P4957, Sigma-Aldrich, USA)-laminin (15 µg/mL; #LN521, Biolamina, Sweden)-coated 6-well plate and 10 µM Y-27632 (#HB2297, Hello Bio, Great Britain) were added for the first 24 h. On days 12 and 17, hiNPCs were passaged with Accutase (#07920, Stemcell Technologies, Canada) and transferred to a new PLO-laminin-coated plate. From day 12, the hiNPCS were cultivated in neural progenitor medium based on GNEIB medium without SB-431542 and LDN-193189, supplemented with 20 ng/mL recombinant human basic FGF (#233-FB, R&D Systems, Germany) and 10 µM Y-27632 (only for the first 24 h after passaging; #HB2297). On day 21, hiNPCs were singularized with Accutase for 10 min at 37 °C and 5 % CO_2_ before they were frozen in neural progenitor medium supplemented with 10 % dimethyl sulfoxide (DMSO, #A994.1, Carl-Roth, Germany) and 10 µM Y-27632.

#### 2.2.3. Stemdiff protocol

The Stemdiff protocol was performed using the STEMdiff™ SMADi Neural Induction Kit (#08581, Stemcell Technologies, Canada) according to the manufacturer’s protocol with minor modifications. Briefly, the hiPSC colonies were singularized as described above and resuspended in STEMdiff™ Neural Induction Medium supplemented with STEMdiff™ SMADi Neural Induction Supplement and 1 % Penicillin/Streptomycin (#P06-07100, PAN-Biotech, Germany). The cells were transferred to a PLO (15 µg/mL; #P4957, Sigma-Aldrich, USA)-laminin (10 µg/mL; #L2020, Sigma-Aldrich, USA)-coated 6-well plate with 10 µM Y-27632 (only for the first 24 h after passaging; #HB2297, Hello Bio, Great Britain). The cells were passaged on day 6 using Accutase (#07920, Stemcell Technologies, Canada) and transferred to a new PLO-laminin-coated 6-well plate with a cell density of 1.5*10^6^ cells per well. On day 12, the hiNPCs were singularized with Accutase and frozen in STEMdiff™ Neural Progenitor Freezing Medium (#05838, Stemcell Technologies, Canada) containing 10 µM Y-27632 (#HB2297, Hello Bio, Great Britain).

### 2.3. Thawing of hiNPCs

The vials of hiNPCs were quickly thawed in the palms of the hand and each vial containing 4 *10^6^ cells was directly diluted in 10 mL of the respective neural progenitor medium with 10 µM Y-27632 (#HB2297, Hello Bio, Great Britain). After centrifugation (300 g, 5 min), the cell pellet was resuspended in the respective neural progenitor medium with 10 µM Y-27632 (#HB2297, Hello Bio, Great Britain). The cells of one frozen cryovial were divided into three wells of a coated 6-well plate. The medium was replaced daily without the addition of Y-27632 (2D-NIM and GNEIB protocol) or with 10 µM Y-27632 (Stemdiff protocol).

### 2.4. Formation of BrainSpheres

On day 4 after thawing, the hiNPCs were singularized with Accutase for 10 min at 37 °C and 5 % CO_2_ (#07920, Stemcell Technologies, Canada) and centrifuged (300 g, 10 min). After the cell pellet was resuspended in the respective neural progenitor medium with 10 µM Y-27632 (#HB2297, Hello Bio, Great Britain), 2 *10^6^ cells were transferred into one well of a new 6-well plate (#83.3920, Sarstedt, Germany) in 4 mL medium. The sphere formation took place in a shaking incubator (#LT-X, Kuhner Shaker GmbH, Swiss) at 140 rpm, 12.5 mm diameter, 37 °C, 5 % CO_2_, and 85 % humidity for 7 days.

### 2.5. Neural Differentiation

For neural differentiation, the BrainSpheres were chopped to 250 µm (McIlwain tissue chopper, Mickle Laboratory Engineering Co. LTD.) and transferred to neural differentiation medium CINDA+ or Electro. CINDA+ consists of 2 parts DMEM (high glucose, #31966021, Invitrogen, USA) and 1 part Ham’s F12 Nutrient Mix (#31765027, Invitrogen, USA) supplemented with 1 % Penicillin/Streptomycin (#P06-07100, PAN-Biotech, Germany), 2 % B-27™ Plus supplement (#A3582801, Thermo Fisher Scientific, USA), 1 % N-2 supplement (#17502048, Thermo Fisher Scientific, USA), 650 µg/mL creatine monohydrate (#C3630, Sigma-Aldrich, USA), 5 ng/mL human recombinant interferon-y (IFN-y, #300-02, Peprotech, Germany), 20 ng/mL human recombinant neurotrophin-3 (#450-03, Peprotech, Germany), 20 µM L-ascorbic acid (A5960, Sigma-Aldrich, USA), and 3 mM N6,2′-O-Dibutyryladenosine 3′,5′-cyclic monophosphate sodium salt (cAMP, #D0260, Sigma-Aldrich, USA). Electro medium consists of 1:1 DMEM/F-12 (#31331028, Gibco, USA) and Neurobasal medium electro (#A14098-01, Gibco, USA) supplemented with 1 % Penicillin/Streptomycin (#P06-07100, PAN-Biotech, Germany), 1 % B-27 supplement electro (#A14097-01, Gibco, USA), 0.5 % N-2 supplement (100x, #17502048, Thermo Fisher Scientific, USA), 0.5 mM GlutaMAX™ supplement (#35050061, Thermo Fisher Scientific, USA), 0.5 % MEM non-essential amino acids (100x, #11140050, Thermo Fisher Scientific, USA), 2.5 µg/mL insulin (#I9278, Sigma-Aldrich, USA), 50 µM beta-mercaptoethanol (#31350010, Thermo Fisher Scientific, USA), 20 ng/mL recombinant human brain derived neurotrophic factor (BDNF, #450-02, Peprotech, Germany), 10 ng/mL recombinant human glial cell line-derived neurotrophic factor (GDNF, #212-GD-010, R&D Systems, Germany), 500 µM cAMP (#D0260, Sigma-Aldrich, USA) and 200 µM L-ascorbic acid (#A5960, Sigma-Aldrich, USA). The BrainSpheres were differentiated in a shaking incubator (#LT-XC, Kuhner Shaker, Germany) at 37 °C, 5 % CO_2_, 85 % humidity, 140 rpm (12.5 mm diameter) for 1, 2, or 3 weeks, and half of the medium was replaced twice a week.

### 2.6. Neural differentiation on microelectrode arrays (MEA)

To access the neuronal electrical activity, the BrainSpheres were plated on 96-well multielectrode arrays (MEA, #M768-tMEA-96B, Axion Biosystems, USA) after 3 weeks, 2 weeks, 1 week, or without 3D differentiation under constant shaking in CINDA+ or Electro differentiation medium. The MEA was coated with specific matrices for each differently generated and differentiated BrainSphere (see **Supplementary Table S1**). After coating with the respective matrix, one BrainSphere was placed in the middle of each well of the MEA, except for BrainSpheres generated with the GNEIB protocol and differentiated in Electro differentiation medium. Here, 3 BrainSpheres were placed per well as they were smaller in size. The BrainSpheres were fed twice per week by replacing half of the differentiation medium. The neuronal electrical activity was recorded twice per week, and BrainSpheres were acutely exposed to L-glutamate (50 µM), DL-2-Amino-5-phosphonovaleric acid (AP5, 50 µM), and NBQX disodium salt (NBQX, 50 µM), bicuculline (3 and 10 µM), picrotoxin (5, 10 and 20 µM), haloperidol (1 and 10 µM), carbaryl (5, 10, 20, 50, and 100 µM), and buspirone hydrochloride (5, 10, 20, 50, and 100 µM) after 2-6 weeks on the MEA. For the substance testing (**Figure 8**), the BrainSpheres were first exposed to the neurotransmitters glutamate (50 µM) or GABA (10 µM) before the antagonists AP5 (50 µM) and NBQX (50 µM) or bicuculline (10 µM) were added. The substances were removed with a complete exchange of the medium and the neuronal networks were allowed to recover for 2 to 3 hours. After another baseline recording, the test compounds trimethyltin chloride (TMT) and emamectin were consecutively added until the final concentration was reached. Detailed information such as CAS registry numbers (CASRN), suppliers, and solvents are listed in **Supplementary Table S2**. The data for subtype characterization were derived from 3 different MEA plates with 8 wells per condition and 8 electrodes per well, resulting in 192 electrodes per condition. The data of the substance testing experiment with TMT, emamectin, and quinpirole were derived from one MEA plate with 8 wells per condition and subtype specification, resulting in 64 electrodes. Statistical analysis was performed with GraphPad Prism (version 8.4.3). First, Brown-Forsythe test were used to detect significant differences between the SDs. If the SDs were not significantly different, one-way ANOVA followed by the post-hoc Dunnett test were applied. If the SDs were significantly different, Brown-Forsythe and Welch ANOVA followed by the post-hoc Games-Howell test were performed.

#### 2.6.1. Recording and data analysis of MEA neuronal electrical activity

Extracellular recording of the neuronal electrical activity was performed twice a week for 15 min (baseline and each tested compound concentration) at 37 °C and 5 % CO_2_ after the cells were allowed to equilibrate for 15 min in the Axion Maestro Pro system (Axion Biosystems, USA). Data recording was operated by the Axion Integrated Studios (AxIS) navigator software (version 3.1.2, Axion Biosystems, USA) with a sampling frequency of 12.5 kHz and a digital band-pass filter of 200-3000 Hz. Subsequent spike detection was performed using the method “adaptive threshold crossing” with a threshold of 6 root mean square (rms) noise on each electrode and a pre- and post-spike duration of 0.84 ms and 2.16 ms, respectively, and an electrode was defined as active with at least 2 spikes per min. Quantification of general electrical activity and neuronal network activity was performed with the Neural Metric Tool software (version 3.1.7, Axion Biosystems, USA). For burst detection, the method “Inter-spike interval (ISI) threshold” was used with a minimum of 5 contributing spikes and a maximum ISI of 100 ms. Network bursts were identified using the algorithm “envelope” with a threshold factor of 1.25, a minimal inter-burst interval (IBI) of 100 ms, at least 35 % participating electrodes, and 75 % burst inclusion. Parameters for neuronal activity (percentage of active electrodes and weighted mean firing rate (wMFR)) as well as for network maturation and synchronicity (burst frequency and network burst frequency) were analyzed.

#### 2.6.2. Spike sorting

For spike sorting, the AxIS generated .spk files of the baseline measurement and each corresponding treatment concentration were concatenated and converted into .nex files with a MATLAB (R2021b, R2022b, MathWorks, USA) script. The generated .nex files were sorted with the neural spike sorting software Offline Sorter (OFS, version 4.4, Plexon, USA) using the automatic clustering T-Distribution E-M method with 10 degrees of freedom (D.O.F) and 20 initial number of units. The sorted units were exported as per-unit and per-waveform data giving information about the number of spikes per unit per baseline and substance concentration. For analysis, only units with at least 2 spikes/min during the baseline measurement were analyzed and they were considered as responding units when the fold change to the baseline was at least ±0.25 (increase or decrease).

### 2.7. Cytotoxicity assessment

Cytotoxicity was assessed by measuring the lactate dehydrogenase (LDH) release from cells with damaged membranes using the CytoTox-ONE Homogeneous Membrane Integrity assay (#7891, Promega, USA). For this, parallel to the substance testing on the MEA, one Brainphere was placed in the middle of each well of a coated 96-well plate (for coating see **Supplementary Table S1**) and differentiated in CINDA+ for 4 weeks. After acute treatment with the respective substance (15 min at 37 °C), 50 µl medium from each well was transferred to a new 96-well plate and 50 µl CytoTox-ONE reagent was added. As lysis control, neurospheres were treated with 10 % Triton-X 100. Medium without spheres was used to correct for background fluorescence. Fluorescence was detected with the Tecan infinite M200 Pro reader (ex: 540 nm; em: 590 nm).

### 2.8. Cultivation of SynFire Cells

SynFire cells (SynFire Co-Culture Kit (MEA), #1010-7.5, NeuCyte, USA) were cultivated according to the manufacturer’s protocol and as described in detail in Bartmann et al. (preprint) [68]. Briefly, the cells were thawed, resuspended in seeding medium, and seeded on PEI (0.1 %, #181978, Sigma, USA)-laminin (20 µg/mL, #23017015, Thermo Fisher Scientific, USA)-coated 48-well MEA plates (#M768-KAP-48, Axion, USA) in a ratio of 140,000 glutamatergic neurons, 60,000 GABAergic neurons, and 70,000 astrocytes per well. On days 3 and 5, half of the medium was replaced with short-term medium. From day 7 onwards, the medium was gradually replaced by long-term medium and the cells were fed twice per week.

### 2.9. Flow cytometry

Flow cytometry analyses were performed to confirm the success of the neural inductions. Therefore, hiNPCs were singularized with Accutase (#07920, Stemcell Technologies, Canada) for 10 min at 37 °C and 5 % CO_2_ before they were stained with Fixable Viability Stain 510 (1:100, #564406, BD Horizon, USA) for 15 min at room temperature (RT) in the dark. Afterwards, they were washed twice by centrifuging (500 g, 5min, RT) and resuspending in Stain Buffer (#554656, BD Pharmingen, USA). Cells were fixed in Fixation Buffer (#554655, BD Cytofix, USA) for 20 min at RT in the dark, washed two times in DPBS (w/o Mg^2+^ and Cl^2+^, #12559069, Gibco) and then permeabilized in Perm Buffer III (#558050, BD Phosflow, USA). After two additional washing steps in Stain buffer, the hiNPCS were stained with PerCP-Cy5.5 mouse anti-Oct3/4 (1:20, #51-9006267, BD Pharmingen, USA), PE mouse anti-human Pax-6 (1:20, #561552, BD Pharmingen, USA), Alexa Fluor 647 mouse anti-Nestin (1:20, #560341, BD Pharmingen, USA), and Alexa Fluor 488 mouse anti-Ki-67 (1:20, #558616, BD Pharmingen, USA) for 30 min at RT in the dark. To exclude unspecific staining, the isotype controls PerCP-Cy5.5 mouse IgG1, κ (1:10, #51-9006272, BD, USA), PE mouse IgG2α, κ (1:20, #558595, BD Phosflow, USA), Alexa Fluor 647 mouse IgG1, κ (1:20, #557732, BD Pharmingen, USA), and Alexa Fluor 488 mouse IgG1, κ (1:5, #557782, BD Phosflow, USA) were used. Data acquisition of 20,000 cells per sample was performed with the BD FACSCanto II (BD, Bioscience, USA). Dead cells, cell debris, and doublets were discarded during the gating and analyzing process with FlowJo (version 10.8.1). Flow cytometry analyses were performed with hiNPCs derived from 3-4 independent neural inductions of each protocol (N=3-4) and statistical analyses were conducted with GraphPad Prism (version 8.4.3) using a two-way-ANOVA followed by Tukey test for correction of multiple comparisons.

### 2.10. Quantitative polymerase chain reaction (qPCR)

For gene expression analyses, samples were collected at different time points indicated in **Figure 1**. Messenger RNA (mRNA) was isolated using the Rneasy Mini Kit (#74104, Qiagen, Germany) and transcribed into cDNA using the QuantiTect Reverse Transcription Kit (#205311, Qiagen, Germany). qPCR was performed in the Rotor-Gene Q Cycler (#9001560, Qiagen, Germany) using the QuantiFast SYBR Green PCR Kit (#204056, Qiagen, Germany). For quantification, standard curves of all examined genes were generated for calculating copy numbers (CN) as described in Walter et al., 2019 [69]. All steps were performed according to manufacturer’s instructions and CN of the gene of interest were normalized to β-actin expression. Primer sequences are listed in **Supplementary Table S3**. The data were derived from 3-4 independent experiments, each performed with a different batch of hiNPCs. Statistical analyses were performed using GraphPad Prism (version 8.4.3). For media comparisons, an unpaired t-test with Welch’s correction was used. For comparison to the first time point (2D or 3D), one-way ANOVA and the post-hoc test Dunnett were used. The calculated p-values for each comparison can be found in **Appendix A**.

**Figure 1.**
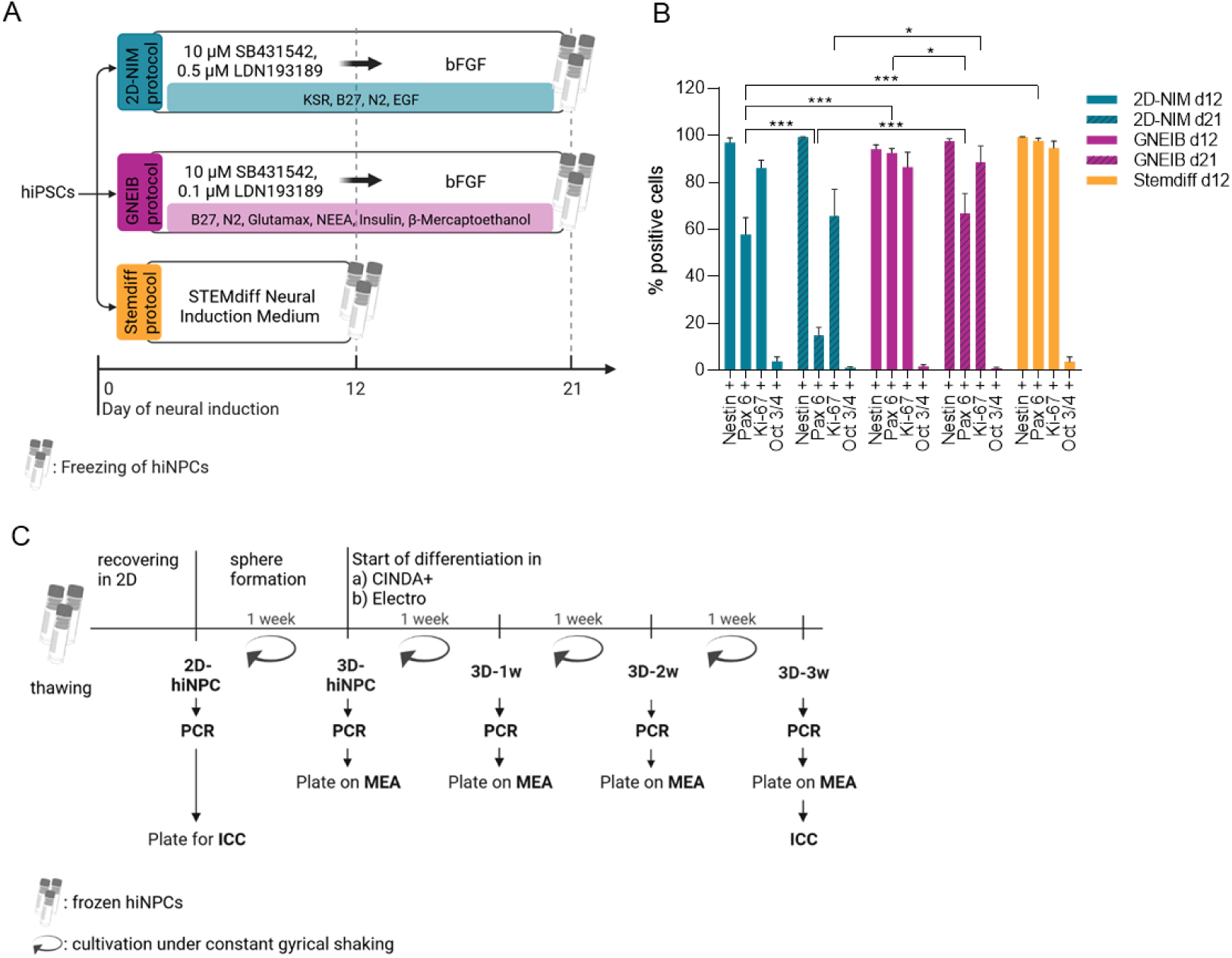
Schematic overview of experimental timeline and characterization of the human induced neural progenitor cells (hiNPCs). **A)** Overview of the three different neural induction protocols used. The hiPSC line IMR90 was neurally induced with the 2D-NIM, the GNEIB, and the Stemdiff protocol at least three times. The resulting hiNPCs were frozen on day 12 (Stemdiff protocol) or day 21 (2D-NIM and GNEIB protocol. **B)** Flow cytometry analysis of hiNPCs on day 12 and day 21 of the neural induction. All induction protocols generated hiNPCs expressing the neural progenitor markers nestin, Pax6, and the proliferation marker Ki-67, whereas the stemness marker Oct 3/4 was not expressed anymore. Data are represented as mean ± SEM of n=3-4 independent experiments (*p ≤ 0.05, **p ≤ 0.01, ***p ≤ 0.001). **C)** Experimental setup of BrainSphere formation and differentiation. After thawing, the hiNPCs were cultivated in 2D to recover and proliferate. On day 4, they were transferred to a shaking incubator, formed BrainSpheres (3D), and were differentiated in CINDA+ or Electro medium for 1 week (3D-1w), 2 weeks (3D-2w), or 3 weeks (3D-3w). Characterization of gene expression (PCR), protein expression (ICC), and electrical activity (MEA) was performed at the indicated time points. ICC, immunocytochemistry; MEA, microelectrode array. Created with biorender.com.

### 2.11. Immunocytochemistry (ICC) of adherent hiNPCs

On day 4 after thawing, hiNPCs were singularized with Accutase (10 min at 37 °C and 5 % CO_2_), 20,000 cells were transferred into each chamber of a coated 8-chamber slide (#354118, BD Falcon, USA) for another 3 days of cultivation before fixing with 4 % paraformaldehyde (PFA, #P6148, Sigma-Aldrich, USA) for 30 min at 37 °C. Unspecific binding sites were blocked by DPBS (w/o Mg^2+^ and Cl^2+^, #12559069, Gibco) containing 2 % bovine serum albumin (BSA, #11920.04, Serva, Germany), 10 % goat serum (GS, #G9023, Sigma-Aldrich, USA), and 0.1 % Triton X-100 (#T8787, Sigma-Aldrich, USA) for 60 min at 37 °C. After the hiNPCs were washed twice with DPBS, they were incubated with antibodies against nestin (anti-nestin AF 647, 1:20, #560341, BD Pharmingen™, Germany) and Ki-67 (anti-Ki-67 AF488, 1:20, #558616, BD Pharmingen™, Germany) in DPBS containing 10 % GS, 0.1 % Triton X-100 and 1 % Hoechst 34580 (#H21486, Thermo Fisher Scientific, USA) for 60 min at 37 °C. After removal of the staining solution, cells were washed three times with DPBS before embedding with Aqua-Poly/Mount (#18606-20, Polysciences Inc., USA) and a cover glass. Images were acquired with the High Content Analysis (HCA) platform Cellinsight CX7 (Thermo Fisher Scientific, USA). Quantification of nestin- and Ki-67-positive cells were performed using the HCS Studio Cellomics Scan software (version 6.6.1) and the protocol TargetActivation.V4. Ki-67-positive cells were identified by setting an intensity threshold within the perimeter of the Hoechst-stained nuclei as Ki-67 is a nuclear protein. Nestin-positive cells were identified by enlarging the perimeter of the detected nuclei to include cytoplasmatic nestin signals near the nucleus, without detecting the signal of neighboring cells.

### 2.12. ICC of BrainSpheres

After three weeks of differentiation under constant shaking, BrainSpheres were fixed with 4 % PFA (#P6148, Sigma-Aldrich, USA) for 60 min at room temperature (RT). Afterwards, they were washed twice with DPBS (w/o Mg^2+^ and Cl^2+^, #12559069, Gibco) and unspecific binding sites were blocked with DPBS containing 10 % GS (#G9023, Sigma-Aldrich, USA) for 30 min at 4°C. The blocking solution was removed and the first staining solution containing 2 % GS, 0.1 % Triton X-100 (#T8787, Sigma-Aldrich, USA), and the primary antibodies against TUBB3 (1:250, anti-TUBB3, Mouse IgG2b, #T8600, Sigma-Aldrich, USA) and S100B (1:500, anti-S100B, rabbit IgG, #ab52642, Abcam, Great Britain) or MAP2 (1:1000 or anti-MAP2, mouse IgG1, #13-1500, Thermo Fisher Scientific, USA) and TH (1:250, anti-TH, rabbit IgG, #ab112, Abcam, Great Britain) were added for 24 h at 4 °C. After the BrainSpheres had been washed twice with DPBS, the second staining solution containing 1 % Hoechst 34580 (#H21486, Thermo Fisher Scientific, USA), 2 % GS and the secondary antibodies (anti-mouse IgG, 1:500, AF488, #A10680, Thermo Fisher Scientific, USA; anti-rabbit IgG, 1:1000, AF546, #A11010, Thermo Fisher Scientific, USA) was added and incubated for 2 h at 4 °C. After washing twice with DPBS, the BrainSpheres were transferred to a microscopy slide, Aqua-Poly/Mount (#18606-20, Polysciences Inc., USA) was added and they were covered with a cover glass. Image acquisition was performed with the confocal laser scanning system 710 (LSM 710, Zeiss) at the Center for Advanced Imaging (CAi) of the Heinrich-Heine-University in Düsseldorf.

## 3. Results

### 3.1. All three neural induction protocols successfully induce hiPSCs into the neural lineage

The hiPSC line IMR-90 was quality controlled and banked according to Tigges et al. 2021 [66]. From a master cell bank a working cell bank was prepared to ensure equal starting material for all experiments. Three different neural induction protocols were compared for their capacity to generate human induced neural progenitor cells (hiNPCs). Two of them were modified from previously published protocols (2D-NIM, [52]; GNEIB, [42,44]) and a third one was commercially available (Stemdiff). The hiPS cell line IMR90 was neurally induced as described in **Figure 1A** and characteristic molecular marker expression was analyzed using flow cytometry on day 12 (all three protocols) and day 21 (2D-NIM and GNEIB) of the protocol (**Figure 1B**). The stemness marker octamer-binding transcription factor 3/4 (Oct 3/4) [36] was downregulated in all protocols to under 4 % positive cells. After 12 days of neural induction, the neural progenitor cell marker nestin [70] was expressed in 97 % (on day 21: 99 %), 94 % (on day 21: 98 %), and 99 % of hiNPCs generated with the 2D-NIM, GNEIB, and Stemdiff protocols, respectively. Expression of the neural progenitor marker PAX6 [71] was unequally expressed in the three protocols, i.e. in 93 % and 98 % of the cells generated with the GNEIB (day 12) and the Stemdiff protocol, respectively, but only in 58 % of cells generated with the 2D-NIM (day 12) protocol. Additionally, the number of cells expressing PAX6 decreased on day 21 for 2D-NIM (15 %) and GNEIB (67 %). The proliferation marker Ki-67 was expressed in 86 % (2D-NIM), 87 % (GNEIB), and 95 % (Stemdiff) of the cells on day 12 and decreased to 66 % (2D-NIM) and 67 % (GNEIB) cells on day 21 of neural induction. After neural induction and FACS evaluation, the hiNPCs were frozen in liquid nitrogen to start each protocol with the same passage number and reduce inter-experimental variability. To ensure that the freezing process did not change molecular marker expression, we confirmed nestin and Ki-67 expression after thawing via immunocytochemistry (**Supplementary Figure S1**). For forming BrainSpheres in 3D, the adherent hiNPCs were cultivated under shaking conditions in 6-well plates according to Honegger et al. 1979 and Pamies et al., 2017 [53,72] using the two differentiation media, i.e. CINDA+ and Electro. These media were chosen for optimization of BrainSphere electrical activity yet do not support oligodendrocyte differentiation. Spheres were characterized with regards to gene and protein expression, as well as neuronal network activity on MEAs in the proliferating state and after 1, 2 and 3 weeks of 3D differentiation (**Figure 1C**).

### 3.2. BrainSpheres differ in neural marker gene expression depending on the applied protocol

For characterizing BrainSpheres’ brain region specificity, developmental stage, neural subtypes, neurotransmitter processing and astrocyte maturation gene expression analyses were performed by qPCR at different time points as indicated in **Figure 1**. As forebrain, midbrain, and hindbrain markers, the genes *FOXG1* (forkhead box G1), *LMX1A* (LIM homeobox transcription factor 1 alpha), *EN1* (engrailed homeobox 1), and *HOXA2* (homeobox A2) were analyzed (**Figure 2, Appendix A**) [39,73]. The forebrain marker *FOXG1* was significantly higher expressed in BrainSpheres generated with the GNEIB protocol and almost not expressed in BrainSpheres generated with the 2D-NIM protocol.

**Figure 2.**
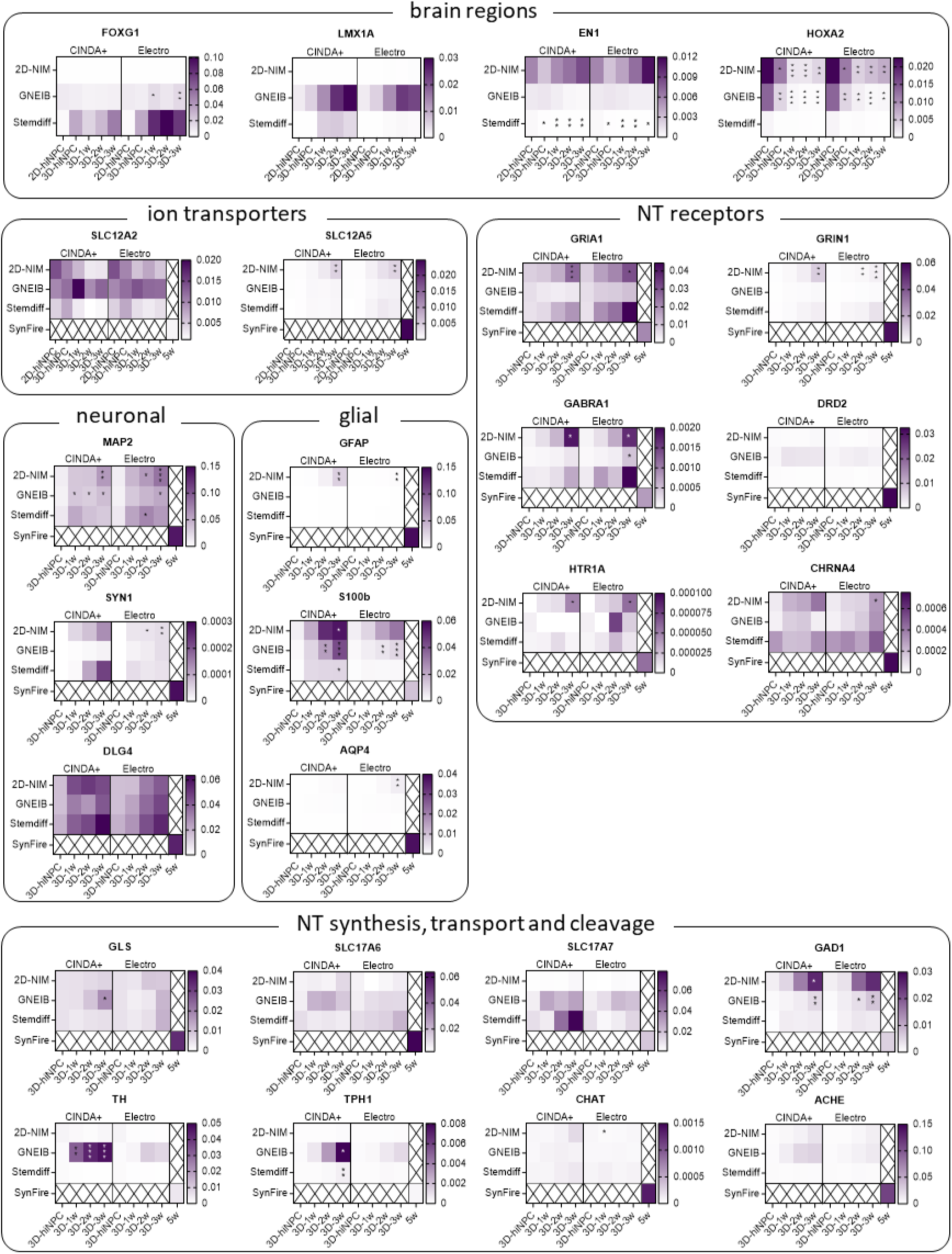
Expression profiles of genes distinguishing different brain regions (FOXG1, LMX1A, EN1, HOXA2), ion transporters defining the stage of the postnatal GABA shift (SLC12A2, SLC12A5), marker for neuronal (MAP2, SYN1, DLG4) and glial (GFAP, S100B, AQP4) cells, neurotransmitter (NT) receptors (GRIA1, GRINA1, GABRA1, DRD2, HTR1A, CHRNA4), NT synthesis enzymes (GLS, GAD1, TH, TPH1, CHAT), NT transporters (SLC17A6, SLC17A7), or NT cleavage enzymes (ACHE). Shown are the copy numbers (CN) of the genes normalized to CN of β-actin. Data are represented as median of n=3-4 independent experiments with 3 technical replicates each (*p ≤ 0.05, **p ≤ 0.01, ***p ≤ 0.001). The p-values resulting from statistical analyses of the comparisons between undifferentiated hiNPCs (2D-hiNPC or 3D-hiNPC) and differentiated BrainSpheres (3D-1w to 3D-3w) and between the 6 different protocols are listed in **Appendix A**.

Although *FOXG1* expression in Stemdiff-derived BrainSpheres was the highest, its induction wasn’t statistically significant due to high standard deviations. The midbrain marker *LMX1A* was highest expressed in BrainSpheres generated with the GNEIB protocol, yet this was not statistically significant. The midbrain marker *EN1* was higher expressed in BrainSpheres derived from the 2D-NIM and GNEIB protocols compared to BrainSpheres derived from the Stemdiff protocol, whose *EN1* expression decreased significantly during differentiation. The hindbrain marker *HOXA2* was highest expressed in the early hiNPC stages of the 2D-NIM and GNEIB protocol, whereas the expression decreased significantly during three weeks of differentiation.

The genes *SLC12A2* (solute carrier family 12 member 2) and *SLC12A5* (solute carrier family 12 member 5) are encoding for the two ion transporters, i.e. Na^+^-K^+^-2Cl^-^ cotransporter-1, (NKCC1) and K^+^-Cl^−^ cotransporter (KCC2), respectively, that are key in the postnatal shift from excitatory to inhibitory GABAergic neurons caused by a reduced expression of *SLC12A2* and an increased expression of *SLC12A5* [74]. While *SLC12A2* expression did not decrease significantly, *SLC12A5* significantly increased in BrainSpheres derived from the 2D-NIM protocol, yet with overall very low copy number expression. For comparison, mRNA expression from a fairly mature, 35-days differentiated neuron-glia mixed culture 2D network, i.e. the SynFire kit [68], was analyzed. *SLC12A2* copy number expression was much lower and *SLC12A5* copy number expression was much higher in the differentiated SynFire cells than in the BrainSpheres indicating prematurity of up to 3 weeks differentiated BrainSpheres independent of the applied protocol.

Expression of *MAP2* (microtubule associated protein 2), which encodes for a dendritic protein [75], significantly increased during BrainSphere differentiation generated with the 2D-NIM and GNEIB protocols. The pre-synaptic marker *SYN1* (synapsin 1) is highest expressed in BrainSpheres generated with the 2D-NIM and the Stemdiff protocols and differentiated in CINDA+, however, this increase is not statistically significant as in 2D-NIM-BrainSpheres differentiated in Electro medium. The postsynaptic *DLG4* (discs large MAGUK scaffold protein 4) expression was abundant in all BrainSpheres even without differentiation and reached similar or even higher expression levels (Stemdiff Electro) than in 35-day differentiated SynFire cells during differentiation.

For analyzing the potential for BrainSphere glial differentiation, the genes *GFAP* (glial fibrillary acidic protein), *S100B* (S100 calcium binding protein B) and *AQP4* (aquaporin 4), which represent different stages of astrocyte maturation, were chosen [76]. *GFAP* was only expressed in BrainSpheres generated with the 2D-NIM protocol and differentiated in CINDA+ for at least 2 weeks. *S100B* was already expressed after one week of differentiation with significantly increased expression in BrainSpheres generated with the 2D-NIM (CINDA+) and the GNEIB protocol (both differentiation media), whereas both conditions differentiated in CINDA+ reached higher expression levels than the 35-day differentiated SynFire cells. *AQP4*, which denotes astrocytes with higher maturity, was hardly expressed in all BrainSpheres, yet highly expressed in SynFire cells.

Expression of genes encoding for the glutamate receptors *GRIA1* (glutamate ionotropic receptor AMPA type subunit 1) and *GRIN1 (*glutamate ionotropic receptor NMDA type subunit 1) were only significantly increased during differentiation in BrainSpheres generated with the 2D-NIM protocol in both differentiation media, however, the *GRIA1* expression in Stemdiff BrainSpheres exceeded the expression of 35-day differentiated SynFire cells. The expression of *GABRA1* (gamma-aminobutyric acid type A receptor subunit alpha 1) increased during differentiation in BrainSpheres generated with the 2D-NIM and the Stemdiff protocols. The gene *DRD2* encoding for the dopamine receptor D2 was not significantly, yet higher expressed in BrainSpheres generated with the GNEIB protocol. The genes *HTR1A* (5-hydroxytryptamine receptor 1A) and *CHRNA4* (cholinergic receptor nicotinic alpha 4 subunit), encoding for serotonin and choline receptors, respectively, were scarcely expressed in all conditions. All receptors were expressed in the SynFire cells.

In addition to neurotransmitter receptors, synthesis and transport of neurotransmitters are also crucial for neural functioning. The enzyme glutaminase (*GLS*) is necessary for glutamate synthesis [77] and the encoding gene was already expressed in the undifferentiated BrainSpheres and increased significantly only in GNEIB-BrainSpheres during differentiation in CINDA+, whereas the gene for vesicular glutamate transporter 1 (*SLC17A7*) was highest expressed in Stemdiff BrainSpheres after 2 and 3 weeks of differentiation in CINDA+, however, this was not statistically significant. The expression of vesicular glutamate transporter 2 (*SLC17A6*) did not strongly increase during differentiation. Glutamate decarboxylase 1 (*GAD1*), which catalyzes a critical step in GABA synthesis [78], was highest expressed in BrainSpheres generated with the 2D-NIM protocol and the expression significantly increased during maturation. The highest expressions of tyrosine hydroxylase (*TH*) and tryptophan hydroxylase 1 (*TPH1*) were found in GNEIB-BrainSpheres differentiated in CINDA+. The genes *CHAT* (choline acetyltransferase) and *ACHE* (acetylcholinesterase) were hardly expressed in all BrainSpheres but highly in the SynFire cells. Additionally, *SLC17A7*, *GAD1*, *TH*, and *TPH1* were higher expressed in some BrainSpheres than in the 35-day differentiated SynFire cells.

### 3.3. Neural induction and differentiation protocols determine the potential of BrainSpheres to differentiate into astrocytes and dopaminergic neurons

Immunofluorescence stainings of BrainSpheres differentiated for 3 weeks revealed their differentiation potentials into astrocytes (S100B) and dopaminergic neurons (TH). BrainSpheres neurally induced with the 2D-NIM protocol differentiated into an abundance of S100B-positive cells that formed a dense layer underneath the TUBB3-positive neurons, regardless of the differentiation medium used. In contrast, BrainSpheres derived from the other two neural induction protocols generated only few or no cells of the astrocytic lineage (**Figure 3A**). Furthermore, BrainSpheres generated with the GNEIB protocol and differentiated in CINDA+ generated the most TH-positive dopaminergic neurons followed by BrainSpheres generated with the 2D-NIM protocol (CINDA+ and Electro) (**Figure 3B**). The other BrainSphere conditions showed no or only few TH-positive neurons. Furthermore, the BrainSpheres had different sphere and neuron morphologies. Especially GNEIB-BrainSpheres differentiated in Electro medium showed a more inhomogeneous distribution of neurons.

**Figure 3.**
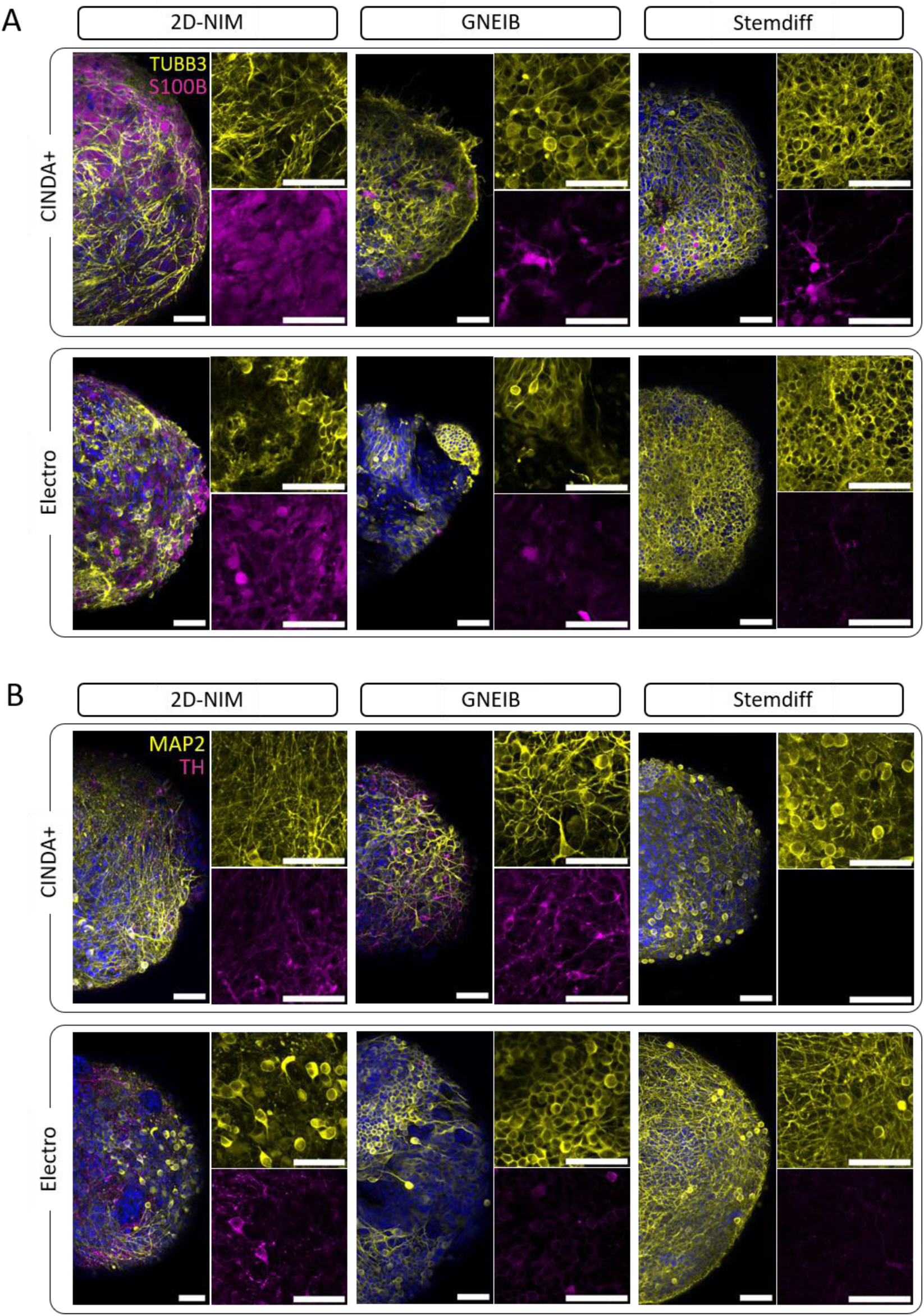
Immunocytochemical characterization of 3-week-differentiated BrainSpheres. **A)** Shown are representative images of BrainSpheres stained for TUBB3 (yellow, neurons) and S100B (magenta, astrocytic lineage). **B)** Representative images of 3-week-differentiated BrainSpheres stained for MAP2 (yellow, dendrites) and TH (magenta, dopaminergic neurons). **A, B)** Nuclei were counterstained with Hoechst 34580 (blue). Scale bar = 50 µM.

### 3.4. Neural induction and differentiation media determine neuronal activity and neural network function of BrainSpheres on MEAs

MEAs are powerful tools to evaluate the electrical activity of neuronal networks providing information on e.g. the number of active electrodes, the rate of action potentials per electrode (weighted mean firing rate; wMFR), the clustering of spikes to bursts as a sign of neuronal maturation (burst frequency) and the neuronal network activity mirroring synchronous communication between different neurons within the neuronal network (network burst frequency). To evaluate and compare the functionality of the neural networks (NN), BrainSpheres were plated on MEAs either without or after 3D differentiation on a gyrical shaker for 1, 2, or 3 weeks, and the electrical activity was measured twice a week for additional 7 weeks. In general, the most active electrodes and the highest wMFR were observed after 3 weeks 3D differentiation before plating BrainSpheres on the MEAs (**Figure 4**, **Supplementary Figures S2, S3, S4**). Therefore, all following experiments were performed with this condition. The number of active electrodes is a valuable parameter as we observed that an inactive electrode does not necessarily indicate that neurons are not growing across the electrode, but a non-electrically active cell may cover it (**Supplementary Figure S5**). The NN generated with the 2D-NIM induction protocol and 3D differentiated in CINDA+ medium showed the most active wells (83 %) and electrodes (58 %) with the highest wMFR of 6.86 Hz after 7 weeks on MEAs (**Figure 4**, **Table 1**). BrainSpheres derived from the GNEIB protocol differentiated in CINDA+ had the second most active wells (83 %), electrodes (31 %), and wMFR (6.30 Hz), but also had the highest variance with a standard error of mean (SEM) of 1.25 Hz. Both NN showed a similar mean burst frequency (2D-NIM: 0.31 Hz, GNEIB: 0.36 Hz) and an increasing activity over the time course of 7 weeks on the MEAs. In contrast, NN derived from BrainSpheres either neurally induced with the 2D-NIM or GNEIB protocol and differentiated in Electro medium, and neurally induced with the Stemdiff protocol and differentiated in CINDA+ medium showed a decrease in the wMFR and burst frequency over the 7 weeks starting from 4 weeks on the MEAs. The NN derived from BrainSpheres neurally induced with the GNEIB protocol and differentiated in CINDA+ had the highest network burst frequency with 0.15 Hz. The percentages of spikes contributing to a network burst (network burst percentage) were similar for all conditions that generated network bursts after 7 weeks on MEA, but highest for the NN generated with the Stemdiff neural induction protocol and subsequent differentiation in Electro medium (48.02 %, **Table 1**). In general, neural induction of hiPSCs using the 2D-NIM and GNEIB protocols followed by BrainSphere differentiation in CINDA+ for 3 weeks generated the most functional NNs with the most active electrodes and highest wMFR, burst frequencies and network burst frequencies, indicating that these protocols are favorable for the generation of mature NN from BrainSpheres.

**Figure 4.**
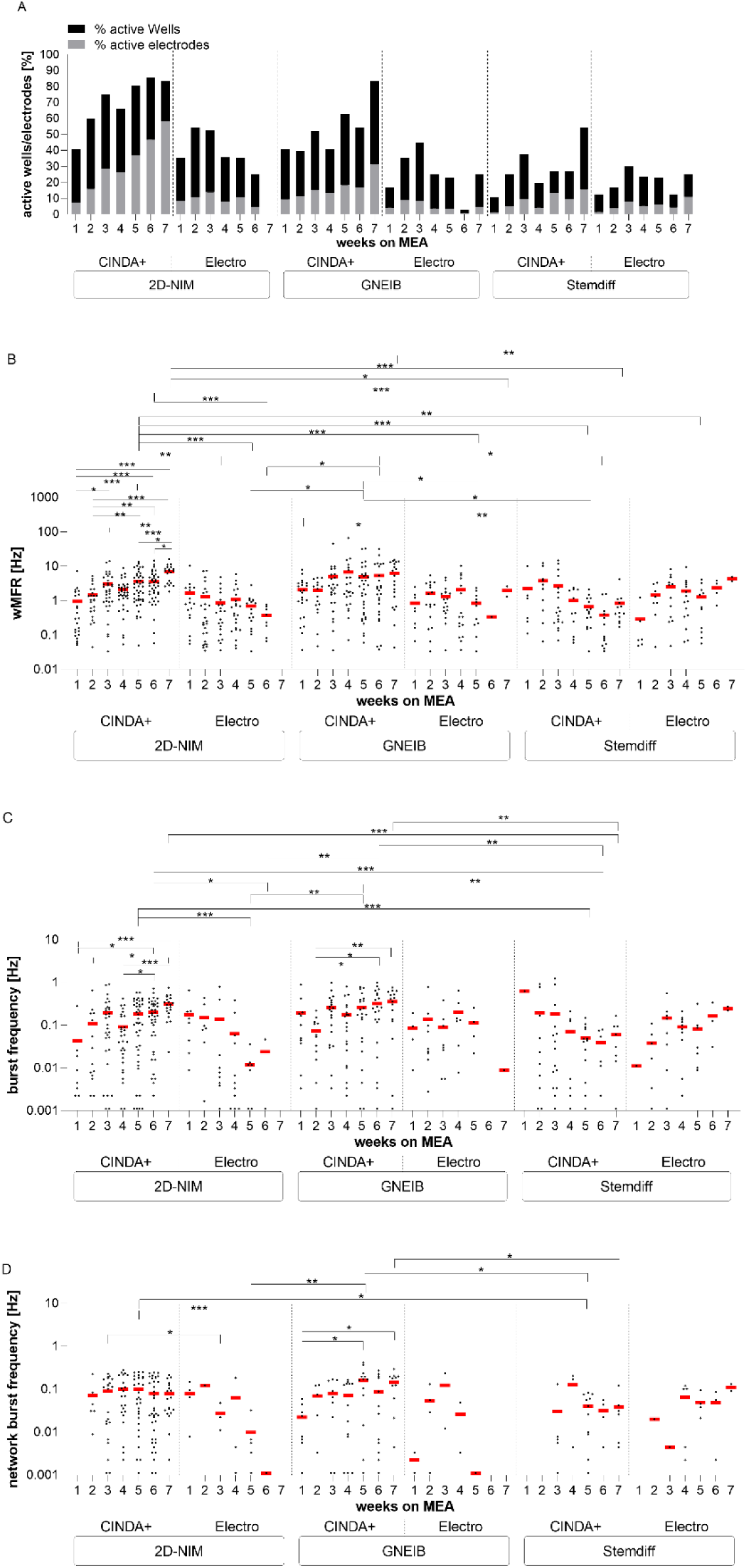
Comparison of electrical activity of 3 weeks 3D differentiated BrainSpheres for 7 weeks on microelectrode arrays (MEA). The neuronal functionality was measured twice per week and the parameters active wells and active electrodes (A), weighted mean firing rate (wMFR, B), burst frequency (C), and network burst frequency (D) were analyzed. Each dot represents the mean of one well containing 8 electrodes and the black, grey (A), and red (B, C, D) bars represents the mean of all wells resulting from three independent MEA experiments each (*p ≤ 0.05, **p ≤ 0.01, ***p ≤ 0.001).

**Table 1.**
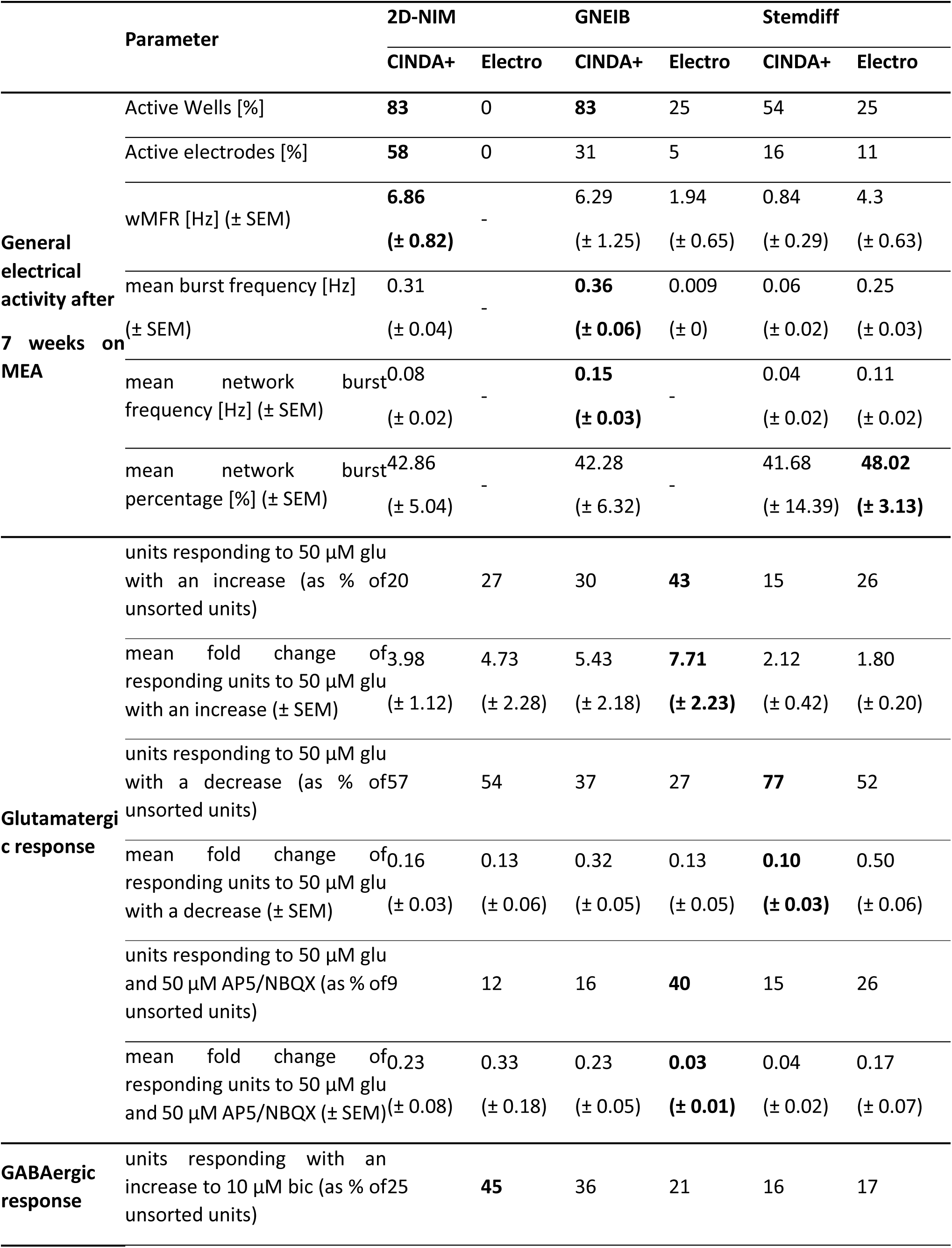
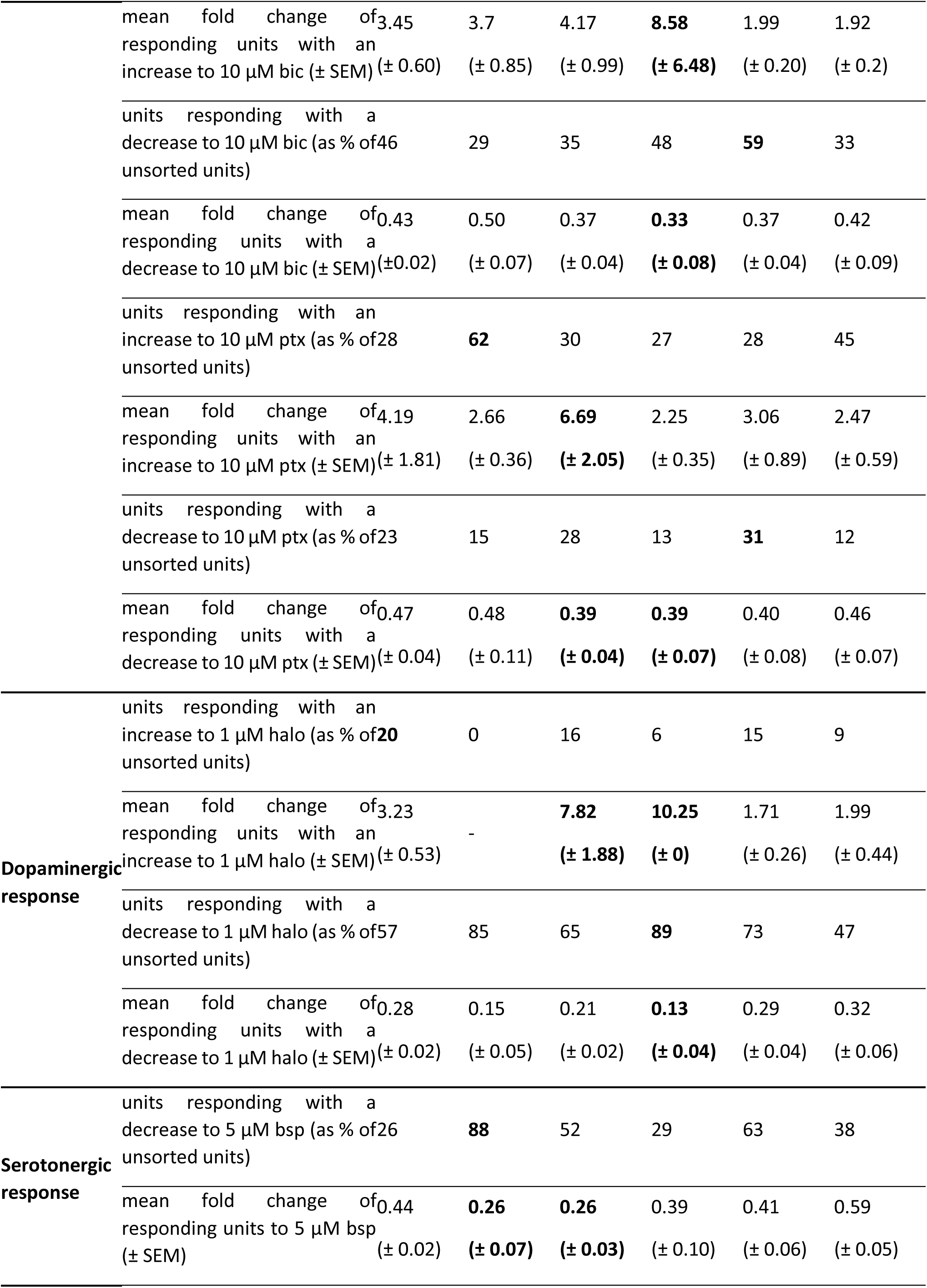
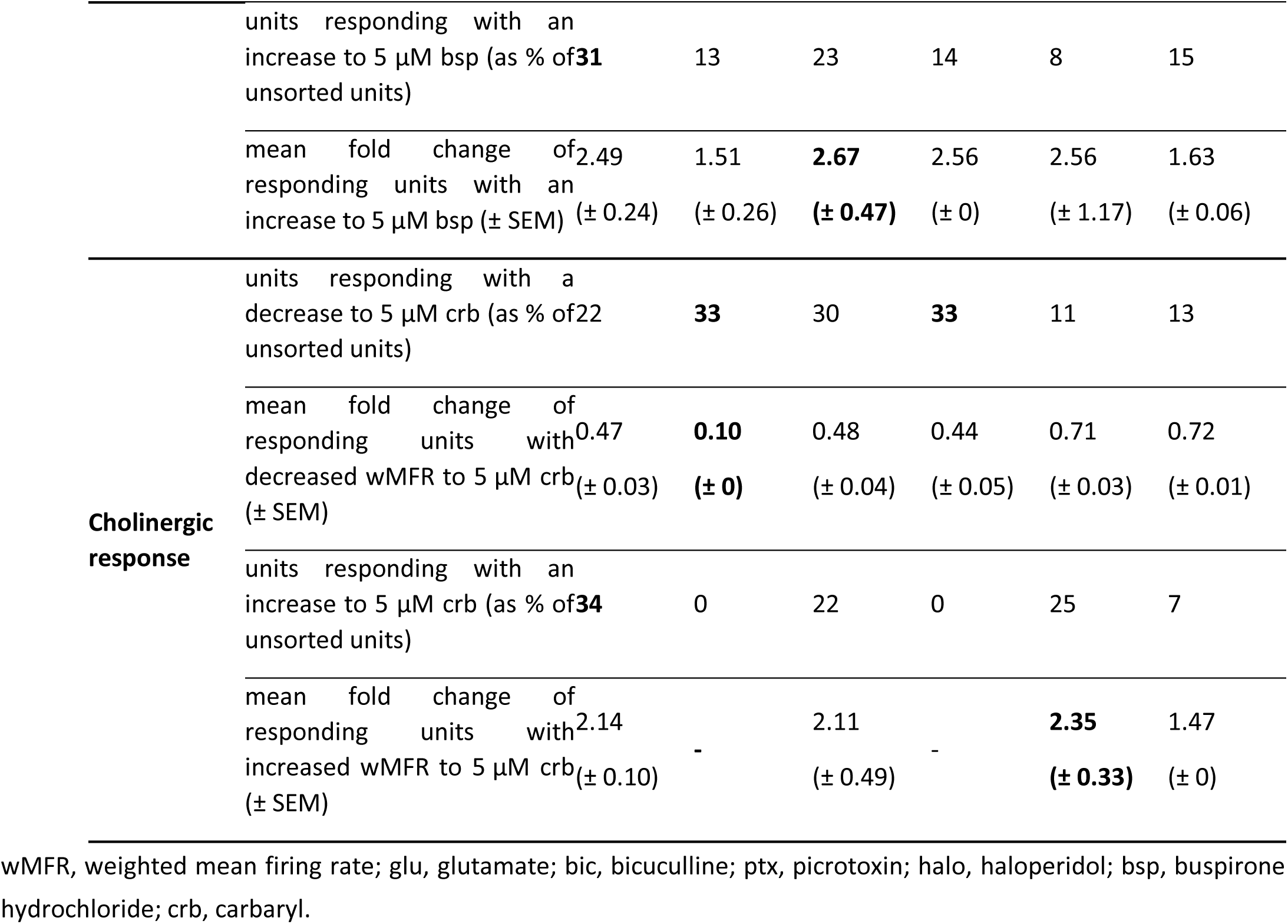
Summary of MEA and spike sorting data of 3 weeks 3D differentiated BrainSpheres.

### 3.5. Neural induction and differentiation media determine BrainSpheres’ neuronal subtype differentiation

For characterization of the NN not only the general electrical activity is important but also the responses to specific pharmacological modulators which highly depends on the occurrence of different neuronal subtypes. Therefore, we addressed the presence of glutamatergic, GABAergic, dopaminergic, serotonergic, and cholinergic neurons in 3 weeks 3D differentiated BrainSpheres after 2 to 6 weeks on MEAs. Quantification of neuronal units’ wMFR responding to pharmacological modulation was possible due to spike sorting with the software Offline Sorter (**Figure 5A**). This allowed the identification of specific responses of individual neurons within the integrated neuronal activities of single electrodes. Neural units reacting to the modulation with a change of at least ±25 % in comparison to the baseline measurement were defined as responding (glutamatergic, GABAergic, dopaminergic, serotonergic, or cholinergic) units. The percentages of responding units, their fold-change to the untreated baseline measurement (colored), and the comparison to the fold-changes of the unsorted (grey) units were analyzed.

**Figure 5.**
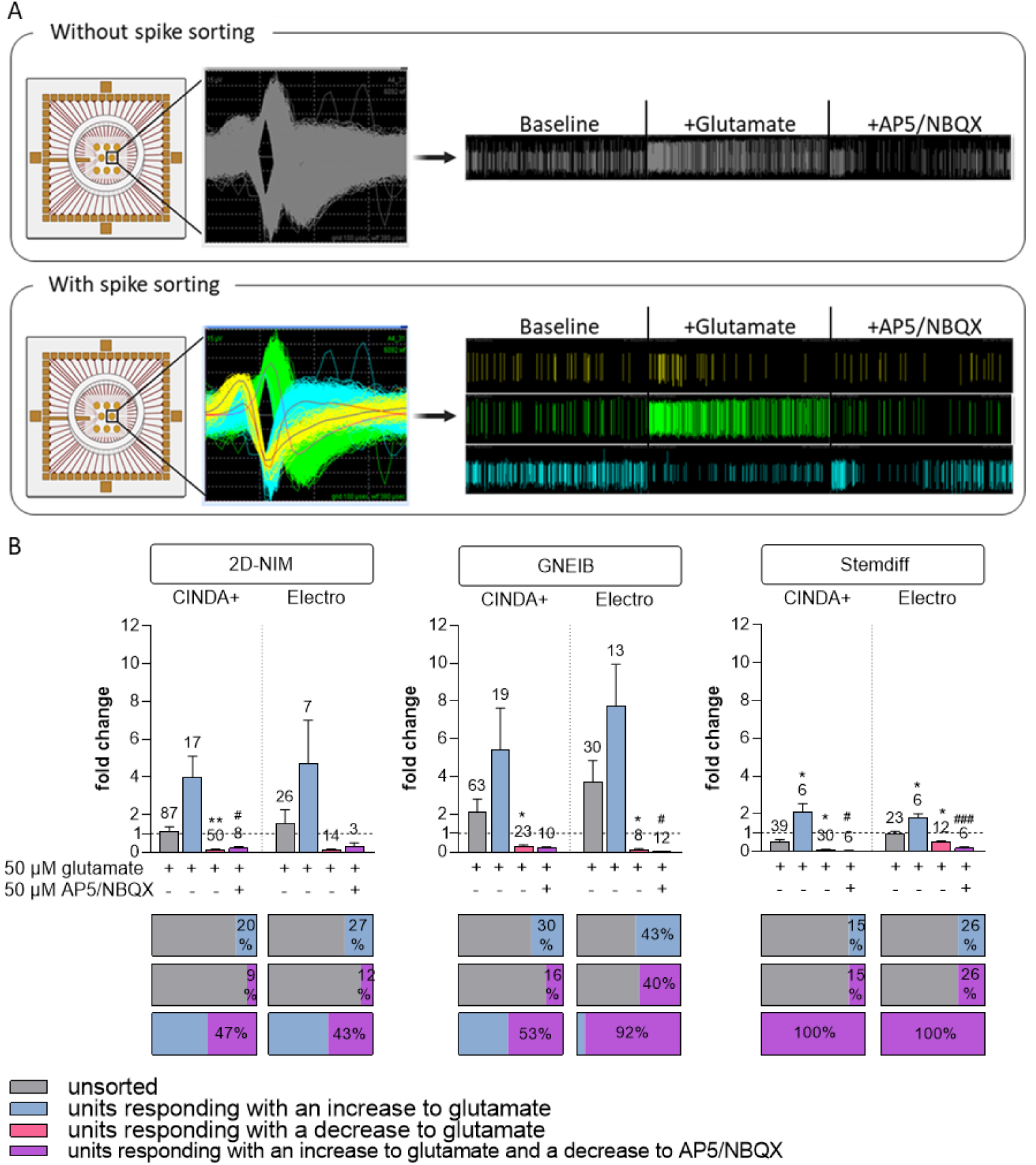
Neuronal network characterization by acute pharmacological modulation to assess glutamatergic response. **A)** Spike sorting of MEA recordings enables the distinction of individual units on the same electrode. Left: Waveforms detected on one electrode. Right: Spike raster plots of the baseline measurement and measurements after exposure to (i) 50 µM glutamate and (ii) 50 µM AP5 and 50 µM NBQX (AP5/NBQX). Each line represents one spike and the length of each measurement is 15 minutes. Shown are exemplary data of NN derived from BrainSpheres generated with the 2D-NIM/CINDA+ media. Top: Unsorted. Bottom: Spike-sorted signals broken down to the individual units. **B)** Brain*S*pheres were 3 weeks 3D differentiated before plated on MEAs and consecutively exposed to glutamate and AP5/NBQX. Shown are the fold changes to the untreated baseline measurements of all units (unsorted) and the responding units after spike sorting. Data are represented as mean ± SEM of three independent MEA experiments with 8 wells per condition (*: significant to unsorted (grey), #: significant to 50 µM glutamate (blue), */#p ≤ 0.05, **/#p ≤ 0.01, ***/###p ≤ 0.001). The numbers above the bars represent the number of units that responded accordingly. Created with biorender.com.

First, the glutamatergic response was measured by applying the neurotransmitter glutamate followed by application of the two glutamate receptor antagonists AP5 (antagonizes NMDA receptors) and NBQX (blocks AMPA receptors) [79]. It is expected that glutamate increases the electrical activity as an excitatory neurotransmitter. Without spike sorting, glutamate-dependent, increased electrical activity was only measured in NN derived from BrainSpheres differentiated with GNEIB/Electro media, whereas BrainSpheres derived from the other five protocols showed no clear positive response or even a decreased wMFR upon acute glutamate exposure (**Figure 5B**). After spike sorting, glutamate-responsive units from BrainSpheres produced with the 2D-NIM and GNEIB protocols produced the highest response to glutamate. Half of glutamate-responding neuronal units in 2D-NIM/CINDA+ and GNEIB/CINDA+ BrainSpheres also responded with a decline in the wMFR to subsequent glutamate receptor inhibition by AP5/NBQX. In contrast, wMFRs of all neuronal glutamate-responsive units from BrainSpheres produced with Stemdiff/CINDA+ and Stemdiff/Electro were inhibited by AP5/NBQX. However, the latter two protocols per se produced very few responding units with minor changes in the wMFR amplitude (**Figure 5B, Table 1**). In addition to the fold changes shown in **Figure 5B**, the raw values of the wMFR are given in the supplementary material (**Supplementary Figure S6**).

To characterize the GABAergic response, the GABA receptor antagonists bicuculline and picrotoxin were applied. Bicuculline binds to GABA_A_ receptors whereas picrotoxin targets GABA_A_ and GABA_C_ receptors [80,81]. It is expected that upon treatment with the GABA receptors antagonists mature GABAergic neurons post the GABA switch respond with an increased activity, e.g. wMFR, while immature GABAergic neurons before the GABA switch respond with a decreased firing. Without spike sorting, none of the BrainSphere cultures derived from any of the six protocols produced changes in NN activity upon GABA receptor antagonism (**Figure 6**). After spike sorting, neuronal units of all BrainSphere protocols were identified that contained increased or decreased wMFR upon bicuculine and picrotoxin treatment, yet most of the responses showed considerable variation in magnitude of responses and hence lacked statistical significance. Only NN from BrainSpheres generated with 2D-NIM/CINDA+, 2D-NIM/Electro, Stemdiff/CINDA+ and Stemdiff/Electro significantly increased the wMFR upon bicuculine treatment, with NN from 2D-NIM/Electro generated BrainSpheres being the least sensitive with the earliest response at 10 µM. In response to picrotoxin only the BrainSpheres generated with 2D-NIM/CINDA+ (5 µM) significantly induced the wMFR of NN. Neuronal units responding with an increased wMFR towards the GABA receptor antagonists ranged between 16 % (Stemdiff/CINDA+) and 62 % (2D-NIM/Electro) of all active neurons. However, the 2D-NIM/CINDA+ media generated the highest absolute number of positively responding neuronal units (Figure 6). Besides the units reacting to the GABA antagonists with increased wMFR, some units responded with a decreased activity. This is in line with the low gene expression of *SLC12A5*, which encodes for the ion receptor KCC2 (**Figure 2**). KCC2 increases its expression after the postnatal GABA switch from excitatory to inhibitory GABAergic neurons. Therefore, without spike sorting, the pre-mature and the more mature GABA receptor containing units compensated each other and resulted in no visible response to the antagonists. BrainSpheres generated with 2D-NIM/CINDA+ media produced the highest absolute numbers of neuronal units responding with an inhibitory and decreased wMFR towards the GABA receptor antagonists. Regarding the Stemdiff neural induction differentiated in CINDA+, in relation to all firing units, 31 % (picrotoxin) to 59 % (bicuculline) of BrainSphere-derived units responded with a decrease in activity towards GABA receptor antagonism (**Figure 6**). The raw values of the wMFR after exposure to bicuculline an picrotoxin are shown in **Supplementary Figures S7 and S8**.

**Figure 6.**
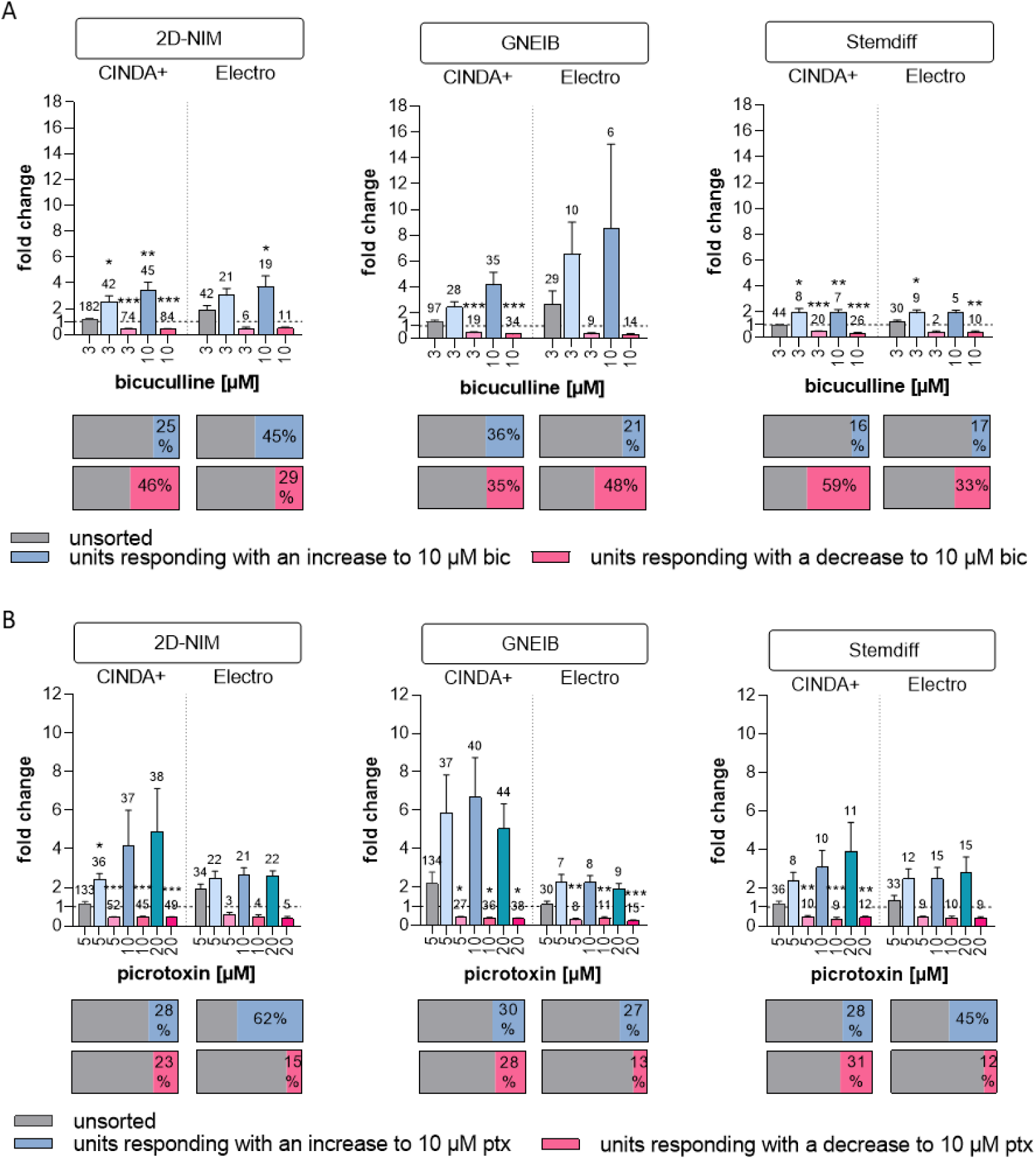
Neuronal network characterization by acute pharmacological modulation to assess GABAergic response. Brain*S*pheres were 3 weeks 3D differentiated before plated on MEA and consecutively exposed to **A)** bicuculline (bic) or **B)** picrotoxin (ptx). Shown are the fold changes to the untreated baseline measurements of all units (unsorted) and the sorted responding units (colored). Data are represented as mean ± SEM of three independent MEA experiments with 8 wells per condition (*: significant to unsorted, *p ≤ 0.05, **p ≤ 0.01, ***p ≤ 0.001). The numbers above the bars represent the number of units that responded accordingly.

Inhibitory dopaminergic D2 receptors were addressed by applying the antagonist haloperidol to the cultures which should increase the wMFR [82,83]. Without spike sorting, haloperidol enhanced the wMFR in GNEIB/CINDA+ (**Figure 7**). After spike sorting, all protocols derived units responding with an increased and decreased wMFR after exposure to haloperidol, except for BrainSpheres from 2D-NIM/Electro media only resulting in units with decreased activity. Haloperidol decreased the wMFR of neuronal units in NN generated with 2D-NIM/CINDA+ and GNEIB/CINDA+ BrainSpheres most effectively with significant wMFR reductions at 1 µM haloperidol in 57 % and 65 % of all neuronal units, respectively. In addition, NN generated with the GNEIB/CINDA+ protocol showed the strongest increased activity at 1 µM. However, most BrainSphere neurons differentiated with 2D-NIM/CINDA+ media seem to express inhibitory D2 receptors as 20 % of all recorded neuronal units responded to 1 µM haloperidol with an increased activity (**Figure 7**). Rising haloperidol concentrations decrease the number of units responding with an increase in wMFR, while the number of units reacting with decreased wMFR enhances under the treatment. The raw wMFR values of the unsorted and the responding units after exposure to haloperidol are shown in **Supplementary Figure S9** and **Table 1**.

**Figure 7.**
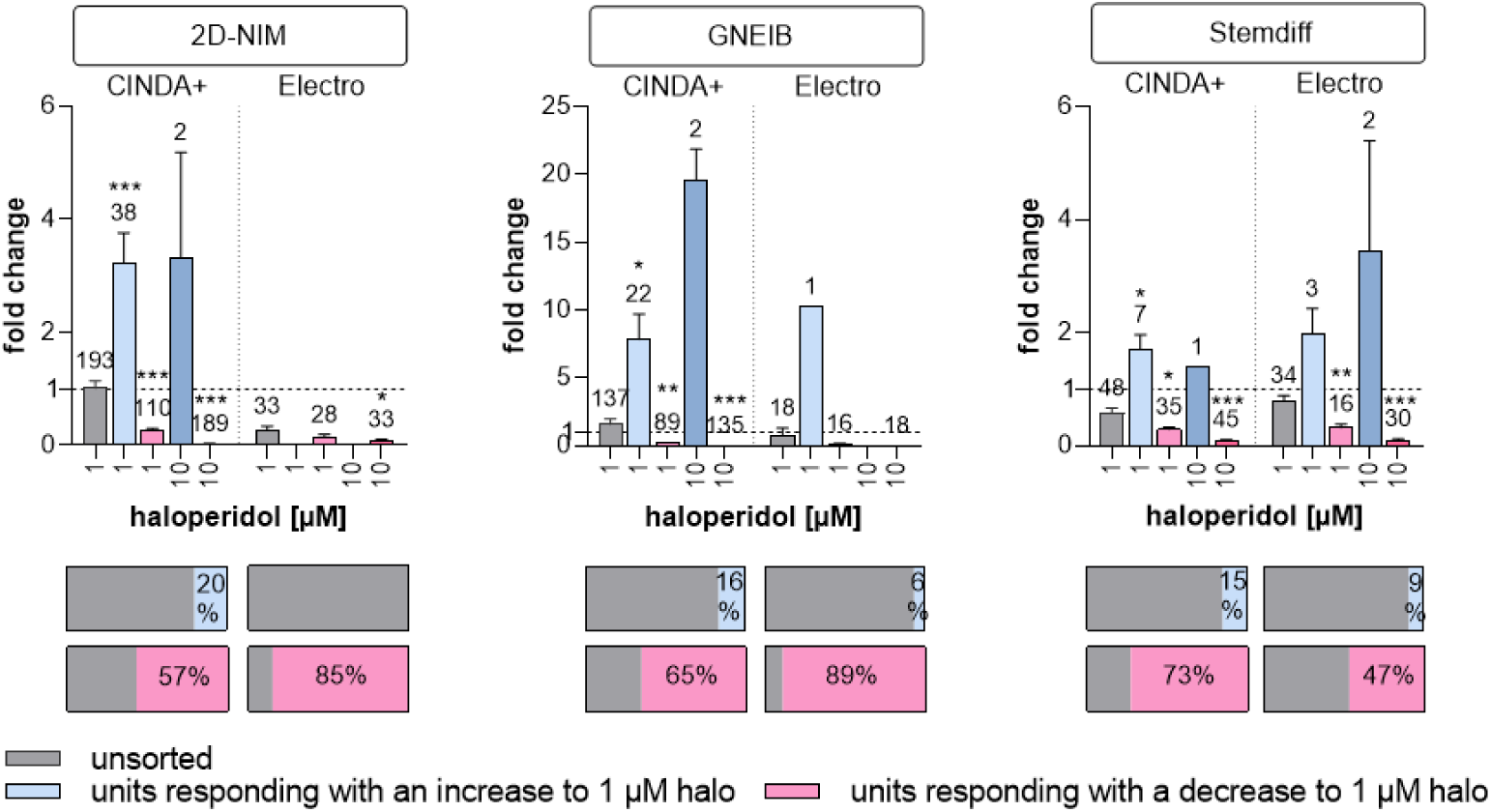
Neuronal network characterization by acute pharmacological modulation to assess dopaminergic responses. BrainSpheres were 3 weeks 3D differentiated before plated on MEAs and exposed to haloperidol (halo). Shown are the fold changes to the untreated baseline measurement of all units (unsorted, grey) and the responding units after sorting (colored). Data are represented as mean ± SEM of three independent MEA experiments with 8 wells per condition (*: significant to unsorted, *p ≤ 0.05, **p ≤ 0.01, ***p ≤ 0.001). The numbers above the bars represent the total number of units that respond accordingly.

Buspirone, which agonize the inhibitory serotonin 5-HT1A receptor, was applied to investigate the presence of serotonergic responses in the NN [84]. Without spike sorting, buspirone did not cause significant changes in neuronal activity in any of the BrainSpheres. However, after sorting, it decreased the wMFR of NN generated with 2D-NIM/CINDA+ and GNEIB/CINDA+ BrainSpheres most effectively with significant wMFR reductions at 5 µM buspirone. In BrainSpheres produced by these protocols, 26 % and 52 % of all neuronal units responded to the compound with a decreased activity, respectively. BrainSpheres differentiated in Electro medium overall produced less serotonergic neurons than CINDA+ differentiated spheres with all neural induction protocols. However, with 88 % of all units, neurons derived from BrainSphere generated with 2D-NIM/Electro media exhibited the highest percentage of neurons responding to buspirone with a decreased activity (**Figure 8, Table 1**). All protocols generated lower percentages of units responding with an increased wMFR to buspirone compared to a decreased activity except for BrainSpheres generated with 2D-NIM/CINDA+, where 31 % of units responded with an increase. In addition to the fold change, the raw values of the wMFR are shown in **Supplementary Figure S10**.

**Figure 8.**
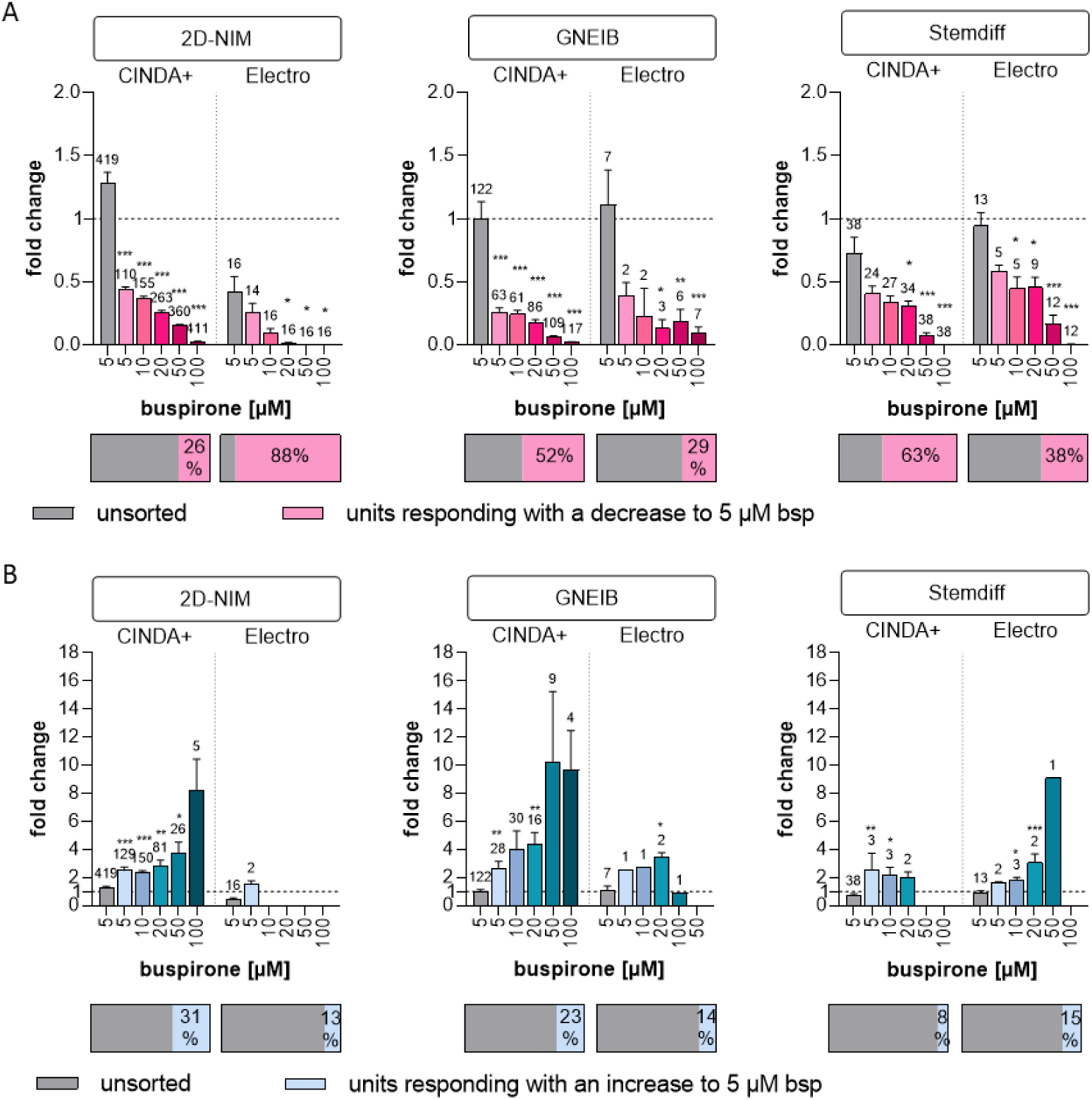
Neuronal network characterization by acute pharmacological modulation to assess serotonergic responses. BrainSpheres were 3 weeks 3D differentiated before plated on MEAs and exposed to buspirone (bsp). **A)** Decreased and **B)** increased responses after exposure to bsp were detected. Shown are the fold changes to the untreated baseline measurement of all units (unsorted, grey) and the responding units after sorting (colored). Data are represented as mean ± SEM of three independent MEA experiments with 8 wells per condition (*: significant to unsorted, *p ≤ 0.05, **p ≤ 0.01, ***p ≤ 0.001). The numbers above the bars represent the total number of units that respond accordingly.

Cholinergic signal transduction was modulated with the insecticide carbaryl, which inhibits the enzyme acetylcholinesterase (AChE) [85] and binds to nicotinic acetylcholine receptors (nAChR) [86]. While AChE blockage increases, binding to nAChR decreases cholinergic neuronal activity [87]. Without spike sorting, no significant change in wMFR was observed after exposure to 5 µM carbaryl (**Figure 9**). After spike sorting, carbaryl decreased the wMFR in NN generated with 2D-NIM/CINDA+ and GNEIB/CINDA+ BrainSpheres most effectively with significant wMFR reductions at 5 µM carbaryl and 22-30 % of all neuronal units responding to the compound. Both, Stemdiff/CINDA+ and Stemdiff/Electro protocols produced BrainSpheres with the least number (11 and 13 %, respectively) and least sensitive (50 and 10 µM carbaryl, respectively) responding neuronal units (**Figure 9A, Table 1**). NN derived from BrainSpheres generated with the 2D-NIM/CINDA+ and GNEIB/CINDA+ protocols also exhibited the most absolute number of neuronal units reacting with the highest increased activity in wMFR to carbaryl (**Figure 9B**). 2D-NIM BrainSpheres were over all the most sensitive to wMFR modulation in both directions significantly responding at 5 µM (**Figure 9**). Interestingly, rising carbaryl concentrations increased the number of neuronal units that responded with a decrease in wMFR, while the number of units reacting with enhanced wMFR decreased under the treatment. In addition to the fold change, the raw values of the wMFR are shown in **Supplementary Figure S11**.

**Figure 9.**
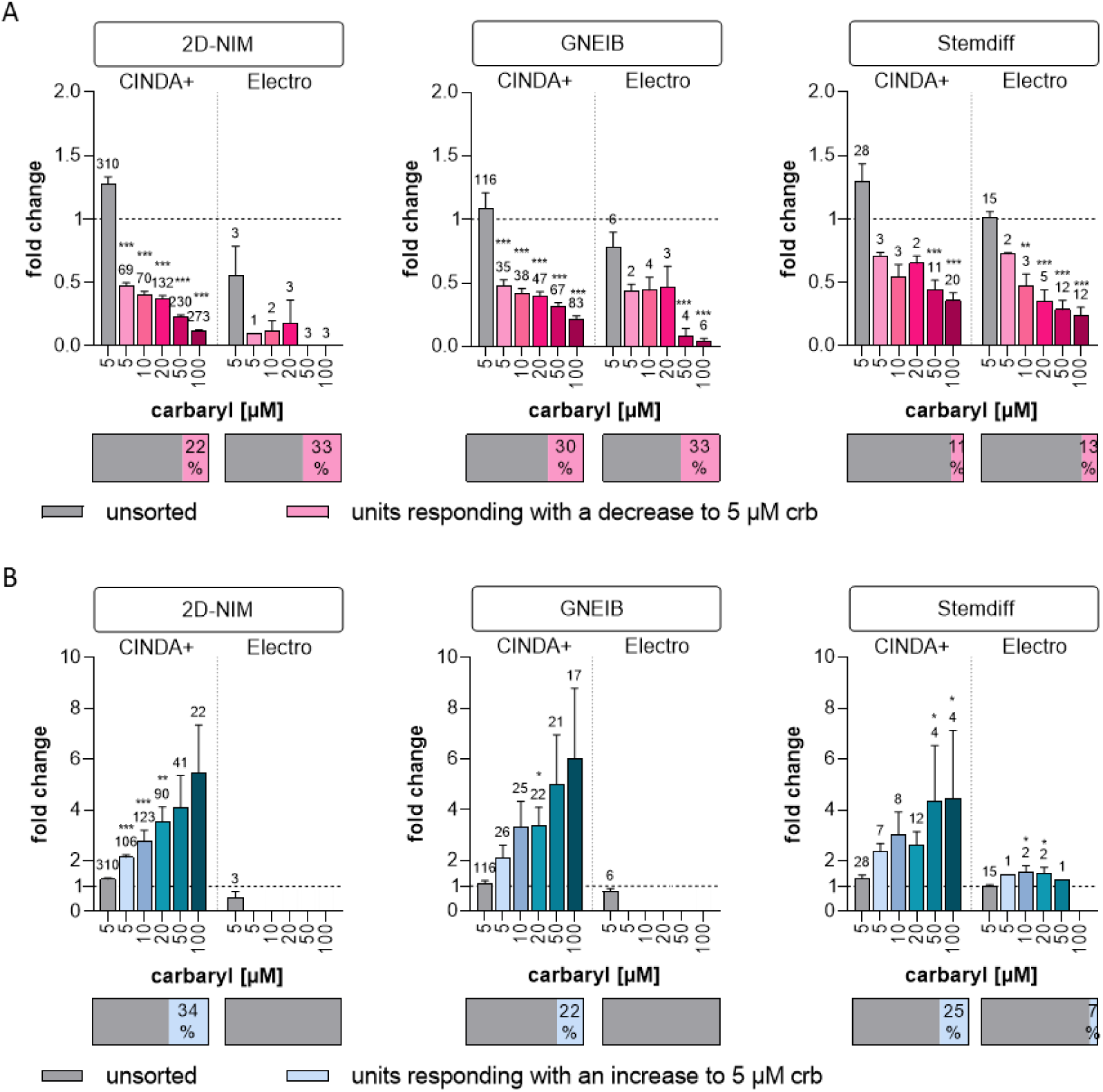
Neuronal network characterization by acute pharmacological modulation to assess cholinergic responses. BrainSpheres were 3 weeks 3D differentiated before plated on MEA and exposed to carbaryl (crb). **A)** Decreased and **B)** increased responses after exposure to crb were detected. Shown are the fold changes to the untreated baseline measurement of all units (unsorted, grey) and the responding units after sorting (colored). Data are represented as mean ± SEM of three independent MEA experiments with 8 wells per condition (*: significant to unsorted, *p ≤ 0.05, **p ≤ 0.01, ***p ≤ 0.001). The numbers above the bars represent the number of units that respond accordingly.

### 3.6. Set-up of a new NAM for acute neurotoxicity testing using MEAs and spike sorting, the human multi-neurotransmitter receptor (hMNR) assay

With the well-characterized BrainSpheres we propose the set-up of a test method as a NAM for acute neurotoxicity testing using MEAs and spike sorting. While general MEA activity can provide an overview of the general changes in NN activity, its resolution is not high enough to understand individual neuronal responses. Spike sorting in combination with neuronal subtype-specific model compounds seems to be a valuable solution for neuronal subtype identification in BrainSpheres on MEAs. To study, if this system is suitable for acute neurotoxicity assessment, we set up a standard operating procedure combining neuronal unit identification with consecutive compound testing (**Figure 10A**). With this set up, as a proof-of-concept, we measured the effects of two compounds, i.e. TMT, which enhances gluatamate release, and emamectin, a GABA-receptor agonist [88,89], on glutamatergic and GABAergic neuronal units in differentiated BrainSpheres (**Figure 10B**). These compounds are the first substances of a chemical training set for the test method and were selected from the mode-of-action analyses in Masjosthusmann et al. 2018 [20]. BrainSpheres generated with the 2D-NIM/CINDA+ protocol were used due to the resulting higher number of active electrodes and lower variance in comparison to the other protocols. Neural units reacting to neurotransmitter and antagonist with a change of at least ±25 % in comparison to the baseline measurement were defined as responding glutamatergic or GABAergic units (**Supplementary Figure S12A)**.

**Figure 10.**
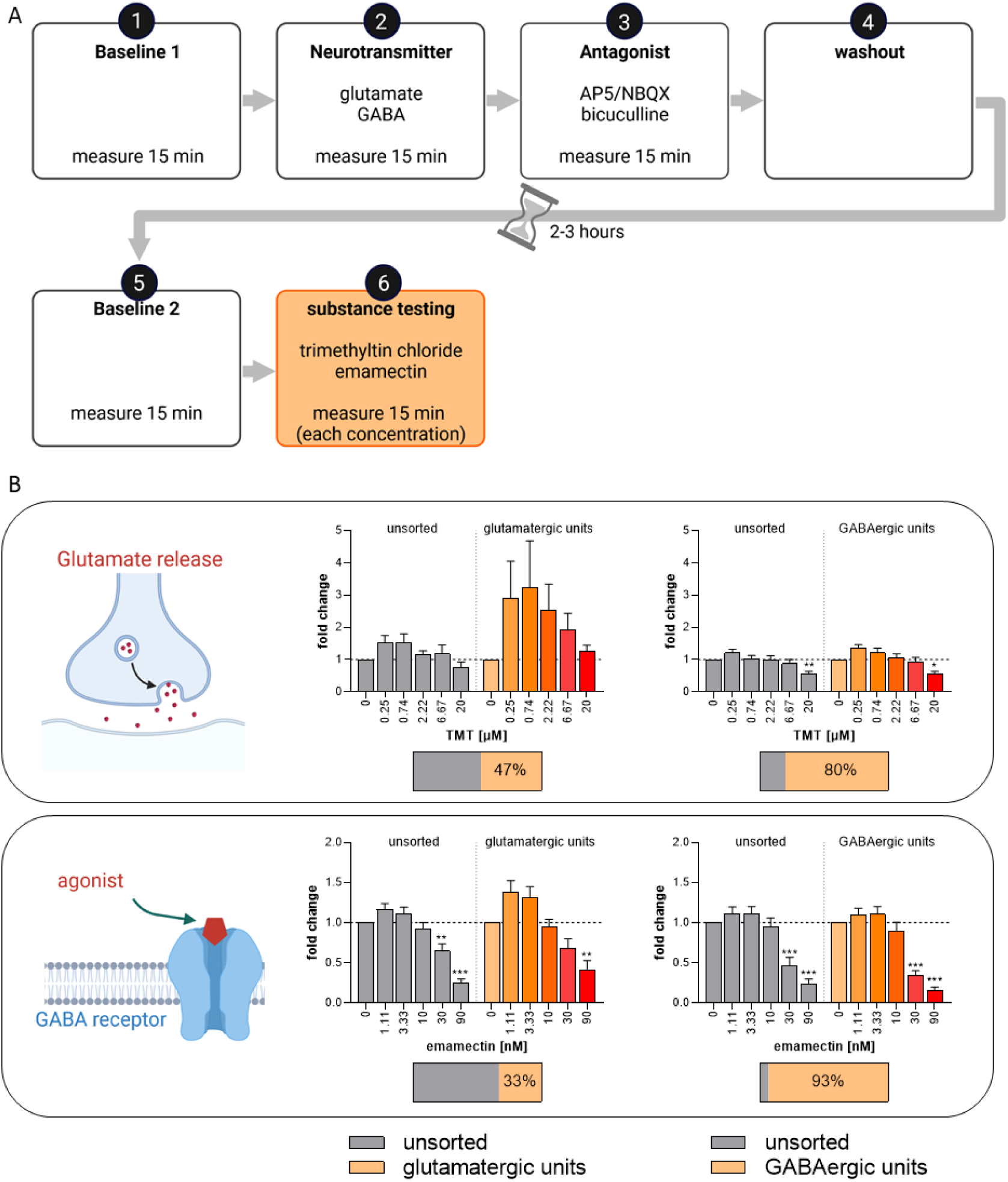
Test method set-up for the NAM assessing neuronal subtype-specific acute neurotoxicity using MEA and spike sorting. **A)** Schematic workflow for acute neurotoxicity testing. After the first baseline recording (baseline 1), first the indicated neurotransmitter (glutamate or GABA) followed by the corresponding antagonist (AP5/NBQX or bicuculline) were applied. The pharmacological modulators were removed by a complete washout and the neuronal networks were allowed to recover for 2 to 3 hours. After a second baseline measurement (baseline 2), the test substances TMT and emamectin were applied in increasing concentrations. **B)** 2D-NIM BrainSpheres were 3 weeks 3D-differentiated in CINDA+. After subsequent 4 weeks differentiation on the MEA, they were used for the described acute neurotoxicity testing. Shown are the fold changes of the wMFR to the respective untreated baseline measurements of all units (unsorted) or pre-sorted responding units (colored). The horizontal bar under each graph indicates the percentages of units responding to the modulation with neurotransmitter and antagonist and were thus defined as glutamatergic or GABAergic units. Data are represented as mean ± SEM (*: significant compared to the baseline, *p ≤ 0.05, **p ≤ 0.01, ***p ≤ 0.001). Created with biorender.com.

After exposure to TMT and respective spike sorting, TMT caused an increased wMFR in glutamatergic units in the sub-micromolar range with a decreasing effect starting at 2.22 µM TMT (**Figure 10B**). This expected increasing effect in wMFR due to enhanced glutamate release by TMT [89] was not observed in the unsorted and GABAergic units. Additionally, the highest TMT concentration (20 µM) also decreased the wMFR of the unsorted and the GABAergic units. Treatment with the GABA_A_ and GABA_C_ receptor agonist emamectin [88] decreased the wMFR of all unsorted and sorted units in a concentration-dependent manner. However, the effect was strongest in sorted GABAergic units with a reduction to a fold change of 0.34 at 30 nM (**Figure 10B**). That the emamectin effects can even be observed in the unsorted units can be explained by the high abundance of GABAergic units in these particular BrainSpheres with 80 to 95% of the units reacting to GABA and bicuculline.

Unspecific cytotoxic effects of the two test compounds were excluded by measuring LDH release (**Supplementary Figure S12B**).

## 4. Discussion

In recent years, industry, regulators, and academia agreed on the need for NAMs to test chemicals with higher throughput, lower costs, and better predictivity for humans [11,13]. For this task, human cell systems designed for a specific purpose should preferably be used and combined with other *in vitro* and *in silico* methods to cover multiple endpoints [90,91]. Human *in vitro* models for acute neurotoxicity testing that examine neurotransmission mainly refer to effects of total neural networks by measuring spike-, burst-, and network-related parameters. While these parameters provide valuable information, they do not necessarily account for a large variety of neurotoxic MoA [29,35,92–96]. The low granularity of the classical MEA evaluation by studying integrated signals over single electrodes is accompanied by high dependence on the NN composition. As especially hiPSC-derived NN are fairly variable [39], high uncertainty might be involved in using spontaneously formed NN from hiPSC for in vitro neurotoxicity studies. Therefore, we characterized mixed culture BrainSpheres [53] for setting up a multiplexed test method for acute neurotoxicity evaluation with the goal of adding multiple neurotoxicity MoA to the established parameters measured with MEAs. First, we characterized six different BrainSphere models resulting from three adherent neural induction protocols combined with two different media for subsequent differentiation. All neural induction protocols showed low variability and high efficiency by resulting in at least 97 % cells expressing the neural progenitor marker nestin [41,70]. However, hiNPC generated with the 2D-NIM, yet not with the GNEIB protocol, contained less Ki-67 positive cells on day 21 of neural induction. This might be the consequence of asymmetrical cell division into proliferative and non-proliferative daughter cells [97]. Pax6 controls various processes regarding the neuroectodermal fate in a concentration-dependent manner and, if absent, leads to asymmetric cell division and thus to neurogenesis [71,98,99]. Hence, the low Pax6 expression in 2D-NIM hiNPCs could explain the decrease in proliferating cells and indicate a more developed state in comparison to the other two protocols.

After the successful neural induction, hiNPCs were frozen in liquid nitrogen. This allows each subsequent experiment to be performed with the same hiNPC passage number, reduces variability and saves time and money. We confirmed, that this additional step does not alter the expression of nestin and Ki-67.

Gene expression data at various differentiation times showed that BrainSpheres produced in different media differ in genes referring to distinct brain regions, synapse formation, astrocyte differentiation, receptors, and neuronal subtypes. Interestingly, the neural induction media influence gene expression more strongly than the differentiation media. Moreover, gene expression comparison to the SynFire neural cells, which are cell ratio-controlled, pre-differentiated excitatory and inhibitory neurons and astrocytes forming functional and highly synchronous neural networks over 35 days in culture, revealed that the BrainSpheres are still fairly immature after 3 weeks of differentiation (e.g. comparing *MAP2*, *SYN1*, *DLG4*, *AQP4* expression). This is supported by the low expression of the K^+^-Cl^-^-co-transporter (KCC2, *SLC12A5*) we observed in BrainSpheres compared to the SynFire cells. The increase in KCC2 expression together with the decrease in expression of the Na^+^-K^+^-2Cl^-^-co-transporter (NKCC1, *SLC21A2*) initiates the postnatal switch from excitatory to inhibitory GABA signaling [74]. The presence of mature and immature GABAergic neurons was also supported by the MEA measurements after exposure to the GABA antagonists bicuculline and picrotoxin, which resulted in both increases and decreases in wMFR. We observed very low expression of genes encoding for serotonin receptor (HTR1A), choline receptor (CHRNA4), and choline synthesis enzyme (CHAT), however, the MEA analysis revealed functional receptors. Previous studies showed that some neuronal subtype-specific markers were only expressed in mature neurons, which can take up to 16 weeks to achieve with hiPSC-derived NN [100,101]. Therefore, such gene expression data in mixed cultures have to be regarded with caution as gene expression measured is an integration of cellular expression and cell abundance in the cultures. Hence, protein analyses using immunocytochemistry combined with functional studies, e.g. using MEAs, are needed for proper test system characterization.

Immunocytochemical stainings for different neuronal and astrocytic markers revealed different potentials of the distinct neural induction/differentiation protocols for differentiation into S100B-positive cells of the astrocyte lineage, with the 2D-NIM/CINDA+ and GNEIB/CINDA+ protocols being the most effective, while GNEIB/Electro and Stemdiff/Electro BrainSpheres only differentiated into very few or no S100B positive cells. These data were supported by the gene expression analyses. Astrocytes are essential for NN maturation and function since they play an important role in synaptogenesis, neuronal survival and outgrowth, phagocytosis, and NT uptake from the synaptic cleft [102,103]. However, spatiotemporal astrocyte marker expression has to be considered when analyzing astrocytes *in vitro*. Data from human *in vivo* investigations reveal that GFAP, a marker most commonly used in *in vitro* studies, should not be used as a general and sole astrocytic marker [76]. Therefore, immunocytochemical analyses supported by qPCR results using a panel of astrocytic lineage markers (e.g. GFAP, S100B, and AQP4) seem reasonable. Astrocyte presence is also important concerning the effects of chemicals, as neurons and astrocytes might react differently to toxic substances [104–106]. Moreover, astrocytes might be the mediators of neuronal toxicity [107,108] or even neuroprotective [109,110]. Hence, astrocytes’ presence in mixed co-cultures is thought to enhance the applicability domain compared to pure cultures.

Not only do astrocytes and neurons respond differently to certain chemicals, but toxic effects on individual neuronal subtypes are frequent causes of neurotoxicity [20,55,111]. Prominent examples are the pesticide rotenone, which acts specifically on dopaminergic neurons [112], or the acetylcholinesterase inhibitor parathion targeting cholinergic neurons [113]. In that regard, it might be of interest to generate fit-for-purpose BrainSpheres enabling the study of distinct neuronal subtypes depending on the scientific or regulatory question. Here, for example, ICC stainings revealed that BrainSpheres generated with the 2D-NIM (both differentiation media) or the GNEIB/CINDA+ protocols generated the highest numbers of TH-positive dopaminergic neurons, while in Stemdiff BrainSpheres they were much fewer. However, besides cell type-specific marker expression, for a physiologically-relevant neural test systems the formation of a functional neuronal network and adequate responses to model compounds have to be demonstrated.

In addition to the mRNA and protein expression data, BrainSpheres were characterized for their performance on MEAs. Similar to the expression analyses, the applied induction and differentiation protocols determined the BrainSphere’s activities on the MEAs. BrainSpheres induced in 2D-NIM and GNEIB media differentiated in CINDA+ showed the most active electrodes, the highest wMFRs, and the highest burst frequencies. The wMFR and bursting behavior depend on various factors, predominantly the presence of astrocytes and the ratio of excitatory to inhibitory neurons [29,30,114]. According to the expression data, these two protocols express the highest levels of the astrocytic lineage marker S100B. Hence, astrocyte presence might be responsible for the abundant firing activity of these BrainSpheres. In contrast to BrainSpheres neurally induced with the 2D-NIM and GNEIB media, BrainSphers generated with Stemdiff medium displayed higher electrical activity when subsequently differentiated with the Electro medium. This indicates that the combination of neural induction and differentiation media highly influences NN functionality. So far, only the influence of different neural induction protocols or various differentiation conditions were analyzed, yet a combination of both had not been examined so far [49,57,115,116].

While spike, burst, and network parameters give important information on general network function, their level of granularity is not particularly high. Therefore, we applied the method of spike sorting to the MEA data [117]. Spike sorting enables the identification of single active neuronal ‘units’ within the signal of one MEA electrode by curve progression analyses. These units can be evaluated individually and hence quantified across multiple electrodes. We challenged the NN with model compounds targeting glutamate, GABA, dopamine, serotonin and nACh receptors, as well as acetylcholine esterase to identify different neuronal subtype signalling.

Without spike sorting, glutamate did not significantly increase wMFR signals on MEAs. After spike sorting, neuronal units were identified that increased or decreased their activity. Neuronal response of increased activity was expected for an excitatory neurotransmitter. The opposite effect might be attributed to either presynaptic metabotropic glutamate receptors (mGluR) that can act as a negative feedback loop and inhibit glutamate release [118–122] possibly in early developing neurons as a counterpart to immature excitatory GABAergic neurons [123]. Such opposite glutamate effects were observed earlier in other neural in vitro models [30,52].

Spike sorting of MEA activity after exposure to the GABA receptor antagonists bicuculline and picrotoxin revealed that all established NN contain both, excitatory and inhibitory GABAergic neurons. This was not visible in the whole electrode recordings, since the two opposite reactions cancel each other out. Excitatory action of GABA is a physiological response before the GABA switch [74]. Hence, all NN contain also immature neurons that precede the GABA switch in addition to inhibitory GABAergic neurons. GABA_A_ and GABA_C_ receptors seem to mature differently as picrotoxin, which binds to GABA_A_ and GABA_C_ receptors [81], increases the increase/decrease ratios of neuronal units compared to bicuculline, which only interacts with GABA_A_ receptors [80], suggesting higher maturation states of GABA_A_ receptors. The gene expression for the two ion transporters NKCC1 and KCC2, which marks the switch from pre-mature excitatory to mature inhibitory GABAergic neurons, supports these functional data.

All BrainSphere models also responded to haloperidol, buspirone, and carbaryl, by directly or indirectly acting on baseline transmission of dopaminergic, serotonergic, and cholinergic receptors, respectively. Overall, differentiation in CINDA+ seems to produce higher numbers of these neuronal subtypes compared to differentiation in Electro medium, possibly due to the higher number of active electrodes these CINDA+ BrainSpheres produce in total. Haloperidol and buspirone bind to dopaminergic and serotonergic receptors, respectively. Haloperidol is an antagonist for the inhibitory dopamine D2 receptor [82,83], hence we observe an increasing effect of this drug on the wMFR of neuronal units. Interestingly, all NN contained more units responding with a decreased electrical activity, which is in line with previous *in vitro* studies that observed only inhibitory reactions after acute treatment [124–128]. Görtz and colleagues suggested, that this effect may occur due to a direct blockage of ion channels [124], which explains the rising number of units responding with decreased activity at the highest haloperidol concentration. Buspirone’s primary MoA is binding to presynaptic inhibitory 5-HT_1A_ receptors as an agonist [84], thus producing an inhibitory action as we observe for most units within the NN *in vitro*. Carbaryl inhibits the enzyme acetylcholine esterase, thereby leading to an accumulation of choline in the synaptic cleft [85]. However, the second MoA of this insecticide is binding to nACh receptors [86,87,129]. Interestingly, previous *in vitro* studies only showed decreased electrical activity after exposure to carbaryl [130–133]. In this study, all NN exhibit neuronal units that respond in both directions, thereby covering both MoA. This might be due to an abundance of nACh receptor expression independent of the cholinergic synapse on GABAergic neurons [134].

Finally, we used the BrainSpheres in combination with spike sorting for setting up a test method, the human multi-neurotransmitter receptor (hMNR) assay for acute neurotoxicity testing that aims at enlightening the neurotoxic MoA for unknown test compounds in the future. As a small proof-of-concept study we exemplified the use of this test method by studying the effects of the compounds TMT and emamectin with already described MoA for glutamatergic and GABAergic neurons. The hMNR confirmed the two MoA of the test compounds: 1) the enhanced glutamatergic activity by TMT-induced glutamate release [89]; 2) the reduced GABAergic neurotransmission caused by the GABA_A_ and GABA_C_ agonist emamectin [88]. Long-term exposure studies showed that TMT also affects synaptic vesicle fusion and recycling [135,136]. This could be a possible explanation for the wMFR reduction at higher TMT concentrations. Although the effect of emamectin was strongest in the GABAergic units, it was also observed in the unsorted and glutamatergic units. This is probably due to the high abundance of GABAergic units in these particular BrainSpheres with 80 to 95% of the units reacting to GABA and bicuculline.

As this is a rather restricted case study, respective proofs-of-concept for the applicabilities of the hMNR assay have to be brought also for the other neuronal subtypes. However, this small set-up already demonstrates for the two model compounds a high sensitivity by detecting compounds’ effects in the nM range. In the end, we envision a test method set-up that identifies all five different neurotransmitter receptors in the first identification phase followed by compound exposure with unknown substances. Spike sorting will identify compounds’ MoA by this effort and hence deliver a neurotoxic MoA profile for each tested compound. The advantage of this system is that one analyzes neuronal units in a mixed neuronal/glia network context, yet information on the individual neuronal level is assessed.

Additional applications for acute substance testing in hiPSC-based BrainSpheres combined with spike sorting analyses are disease modeling and drug development. The pathophysiology of several neurological disorders such as Rett syndrome, autism spectrum disorders, schizophrenia, Down Syndrome, and fragile X involves amongst others a disrupted GABA switch during brain development leading to an inhibitory/excitatory imbalance [137–140]. Moreover, they are suited as Parkinson’s disease model and for untargeted disease modeling revealing so far unstudied disease mechanisms as well as gene-environment interactions [59,141]. Another application can be envisioned in drug development for safety or efficacy evaluation, e.g. for seizure liability assessment [142–145]. In addition, interference of compounds with neurotransmitter systems might also be an indication for their developmental neurotoxicity (DNT) potential. Therefore, this test method might in the future also be a valuable addition to the current DNT in vitro testing battery [146] as test methods for substances’ effects on neuronal subtypes were identified as one gap in current NAM-based DNT evaluation [147].

Taken together, this work thoroughly characterized different BrainSphere protocols for acute neurotoxicity test method set up using fit-for-purpose models. One proof-of-principle indicating the future applicability of the multicellular BrainSphere models as the basis for the hMNR test method was given. Possible applications of this test method beyond acute neurotoxicity include disease modeling, safety and efficacy in drug development and DNT.

## Author Contributions

Conceptualization, J.H., K.K, and E.F.; methodology, J.H., E.F..; software, J.H., and A.D..; formal analysis, J.H, N.H., and A.D.; investigation, J.H., N.H., and K.B.; resources, E.F; writing—original draft preparation, J.H; writing—review and editing, J.H., K.K., and E.F.; visualization, J.H.; supervision, K.K., E.F.; project administration, E.F.; funding acquisition, E.F. All authors have read and agreed to the published version of the manuscript.

## Acknowledgments

The confocal imaging was performed at the Center for Advanced Imaging (CAI) at Heinrich Heine University (HHU), Düsseldorf. **Figures 1A, 1C, 5A**, and **10** were created with BioRender.com.

## References

1. OECD Guideline for the Testing of Chemicals 419 Delayed Neurotoxicity of Organophosphorus Substances: 28-Day Repeated Dose Study. 1995, 1–7.

2. OECD Guideline for the Testing of Chemicals 418 Delayed Neurotoxicity of Organophosphorus Substances Following Acute Exposure. 1995, 1–8.

3. OECD guideline for the testing of Chemicals Test No. 424: Neurotoxicity Study in Rodents. In OECD guideline for the testing of Chemicals; OECD Guidelines for the Testing of Chemicals, Section 4; OECD, 1997; pp. 1–15 ISBN 9789264071025.

4. Buschmann, J. The OECD Guidelines for the Testing of Chemicals and Pesticides. Methods Mol. Biol. 2013, 947, 37–56, doi:10.1007/978-1-62703-131-8_4.

5. Bal-Price, A.K.; Hogberg, H.T.; Buzanska, L.; Coecke, S. Relevance of in Vitro Neurotoxicity Testing for Regulatory Requirements: Challenges to Be Considered. Neurotoxicol. Teratol. 2010, 32, 36–41, doi:10.1016/j.ntt.2008.12.003.

6. Somel, M.; Liu, X.; Tang, L.; Yan, Z.; Hu, H.; Guo, S.; Jiang, X.; Zhang, X.; Xu, G.; Xie, G.; et al. MicroRNA-Driven Developmental Remodeling in the Brain Distinguishes Humans from Other Primates. PLoS Biol. 2011, 9, doi:10.1371/journal.pbio.1001214.

7. Pollen, A.A.; Nowakowski, T.J.; Chen, J.; Retallack, H.; Sandoval-Espinosa, C.; Nicholas, C.R.; Shuga, J.; Liu, S.J.; Oldham, M.C.; Diaz, A.; et al. Molecular Identity of Human Outer Radial Glia during Cortical Development. Cell 2015, 163, 55–67, doi:10.1016/j.cell.2015.09.004.

8. Waring, M.J.; Arrowsmith, J.; Leach, A.R.; Leeson, P.D.; Mandrell, S.; Owen, R.M.; Pairaudeau, G.; Pennie, W.D.; Pickett, S.D.; Wang, J.; et al. An Analysis of the Attrition of Drug Candidates from Four Major Pharmaceutical Companies. Nat. Rev. Drug Discov. 2015, 14, 475–486, doi:10.1038/nrd4609.

9. Mohs, R.C.; Greig, N.H. Drug Discovery and Development: Role of Basic Biological Research. Alzheimer’s Dement. Transl. Res. Clin. Interv. 2017, 3, 651–657, doi:10.1016/j.trci.2017.10.005.

10. Wang, Z.; Walker, G.W.; Muir, D.C.G.; Nagatani-Yoshida, K. Toward a Global Understanding of Chemical Pollution: A First Comprehensive Analysis of National and Regional Chemical Inventories. Environ. Sci. Technol. 2020, 54, 2575–2584, doi:10.1021/acs.est.9b06379.

11. Pallocca, G.; Moné, M.J.; Kamp, H.; Luijten, M.; Van de Water, B.; Leist, M. Next-Generation Risk Assessment of Chemicals - Rolling out a Human-Centric Testing Strategy to Drive 3R Implementation: The RISK-HUNT3R Project Perspective. ALTEX 2022, 39, 1–7, doi:10.14573/altex.2204051.

12. Dent, M.P.; Vaillancourt, E.; Thomas, R.S.; Carmichael, P.L.; Ouedraogo, G.; Kojima, H.; Barroso, J.; Ansell, J.; Barton-Maclaren, T.S.; Bennekou, S.H.; et al. Paving the Way for Application of next Generation Risk Assessment to Safety Decision-Making for Cosmetic Ingredients. Regul. Toxicol. Pharmacol. 2021, 125, 105026, doi:10.1016/j.yrtph.2021.105026.

13. Krewski, D.; Andersen, M.E.; Tyshenko, M.G.; Krishnan, K.; Hartung, T.; Boekelheide, K.; Wambaugh, J.F.; Jones, D.; Whelan, M.; Thomas, R.; et al. Toxicity Testing in the 21st Century: Progress in the Past Decade and Future Perspectives; Springer Berlin Heidelberg, 2020; Vol. 94; ISBN 0123456789.

14. Kavlock, R.J.; Bahadori, T.; Barton-Maclaren, T.S.; Gwinn, M.R.; Rasenberg, M.; Thomas, R.S. Accelerating the Pace of Chemical Risk Assessment. Chem. Res. Toxicol. 2018, 31, 287–290, doi:10.1021/acs.chemrestox.7b00339.

15. Vinken, M.; Benfenati, E.; Busquet, F.; Castell, J.; Clevert, D.A.; de Kok, T.M.; Dirven, H.; Fritsche, E.; Geris, L.; Gozalbes, R.; et al. Safer Chemicals Using Less Animals: Kick-off of the European ONTOX Project. Toxicology 2021, 458, 1–7, doi:10.1016/j.tox.2021.152846.

16. Carmichael, P.L.; Baltazar, M.T.; Cable, S.; Cochrane, S.; Dent, M.; Li, H.; Middleton, A.; Muller, I.; Reynolds, G.; Westmoreland, C.; et al. Ready for Regulatory Use: NAMs and NGRA for Chemical Safety Assurance. ALTEX 2022, 39, 359–366, doi:10.14573/altex.2204281.

17. Maertens, A.; Golden, E.; Luechtefeld, T.H.; Hoffmann, S.; Tsaioun, K.; Hartung, T. Probabilistic Risk Assessment - The Keystone for the Future of Toxicology. ALTEX 2022, 39, 3–29, doi:10.14573/altex.2201081.

18. Mahony, C.; Ashton, R.S.; Birk, B.; Boobis, A.R.; Cull, T.; Daston, G.P.; Ewart, L.; Knudsen, T.B.; Manou, I.; Maurer-Stroh, S.; et al. New Ideas for Non-Animal Approaches to Predict Repeated-Dose Systemic Toxicity: Report from an EPAA Blue Sky Workshop. Regul. Toxicol. Pharmacol. 2020, 114, 104668, doi:10.1016/j.yrtph.2020.104668.

19. van der Stel, W.; Carta, G.; Eakins, J.; Delp, J.; Suciu, I.; Forsby, A.; Cediel-Ulloa, A.; Attoff, K.; Troger, F.; Kamp, H.; et al. New Approach Methods (NAMs) Supporting Read-Across: Two Neurotoxicity AOP-Based IATA Case Studies. ALTEX 2021, 38, 615–635, doi:10.14573/altex.2103051.

20. Masjosthusmann, S.; Barenys, M.; El-Gamal, M.; Geerts, L.; Gerosa, L.; Gorreja, A.; Kühne, B.; Marchetti, N.; Tigges, J.; Viviani, B.; et al. Literature Review and Appraisal on Alternative Neurotoxicity Testing Methods. EFSA Support. Publ. 2018, 15, 1–108, doi:10.2903/sp.efsa.2018.en-1410.

21. Harry, G.J.; Tiffany-Castiglioni, E. Evaluation of Neurotoxic Potential by Use of in Vitro Systems. Expert Opin. Drug Metab. Toxicol. 2005, 1, 701–713, doi:10.1517/17425255.1.4.701.

22. Lisek, M.; Boczek, T.; Stragierowicz, J.; Wawrzyniak, J.; Guo, F.; Klimczak, M.; Kilanowicz, A.; Zylinska, L. Hexachloronaphthalene (HxCN) Impairs the Dopamine Pathway in an in Vitro Model of PC12 Cells. Chemosphere 2022, 287, doi:10.1016/j.chemosphere.2021.132284.

23. Schultz, L.; Zurich, M.G.; Culot, M.; da Costa, A.; Landry, C.; Bellwon, P.; Kristl, T.; Hörmann, K.; Ruzek, S.; Aiche, S.; et al. Evaluation of Drug-Induced Neurotoxicity Based on Metabolomics, Proteomics and Electrical Activity Measurements in Complementary CNS in Vitro Models. Toxicol. Vitr. 2015, 30, 138–165, doi:10.1016/j.tiv.2015.05.016.

24. Hausherr, V.; Thriel, C. van; Krug, A.; Leist, M.; Schöbel, N. Impairment of Glutamate Signaling in Mouse Central Nervous System Neurons in Vitro by Tri-Ortho-Cresyl Phosphate at Noncytotoxic Concentrations. Toxicol. Sci. 2014, 142, 274–284, doi:10.1093/toxsci/kfu174.

25. Loser, D.; Schaefer, J.; Danker, T.; Möller, C.; Brüll, M.; Suciu, I.; Ückert, A.K.; Klima, S.; Leist, M.; Kraushaar, U. Human Neuronal Signaling and Communication Assays to Assess Functional Neurotoxicity. Arch. Toxicol. 2021, 95, 229–252, doi:10.1007/s00204-020-02956-3.

26. Kosnik, M.B.; Strickland, J.D.; Marvel, S.W.; Wallis, D.J.; Wallace, K.; Richard, A.M.; Reif, D.M.; Shafer, T.J. Concentration–Response Evaluation of ToxCast Compounds for Multivariate Activity Patterns of Neural Network Function. Arch. Toxicol. 2020, 94, 469–484, doi:10.1007/s00204-019-02636-x.

27. Johnstone, A.F.M.; Gross, G.W.; Weiss, D.G.; Schroeder, O.H.-U.; Gramowski, A.; Shafer, T.J. Microelectrode Arrays: A Physiologically Based Neurotoxicity Testing Platform for the 21st Century. Neurotoxicology 2010, 31, 331– 350, doi:10.1016/J.NEURO.2010.04.001.

28. Vassallo, A.; Chiappalone, M.; De Camargos Lopes, R.; Scelfo, B.; Novellino, A.; Defranchi, E.; Palosaari, T.; Weisschu, T.; Ramirez, T.; Martinoia, S.; et al. A Multi-Laboratory Evaluation of Microelectrode Array-Based Measurements of Neural Network Activity for Acute Neurotoxicity Testing. Neurotoxicology 2017, 60, 280–292, doi:10.1016/j.neuro.2016.03.019.

29. Saavedra, L.; Wallace, K.; Freudenrich, T.F.; Mall, M.; Mundy, W.R.; Davila, J.; Shafer, T.J.; Wernig, M.; Haag, D. Comparison of Acute Effects of Neurotoxic Compounds on Network Activity in Human and Rodent Neural Cultures. Toxicol. Sci. 2021, 180, 295–312, doi:10.1093/toxsci/kfab008.

30. Tukker, A.M.; De Groot, M.W.G.D.M.; Wijnolts, F.M.J.; Kasteel, E.E.J.J.; Hondebrink, L.; Westerink, R.H.S.S.; Tukker, A.M. Is the Time Right for in Vitro Neurotoxicity Testing Using Human IPSC-Derived Neurons? ALTEX 2016, 33, 261–271, doi:10.14573/altex.1510091.

31. Tukker, A.M.; van Kleef, R.G.D.M.; Wijnolts, F.M.J.; de Groot, A.; Westerink, R.H.S. Towards Animal-Free Neurotoxicity Screening: Applicability of HiPSC-Derived Neuronal Models for in Vitro Seizure Liability Assessment. ALTEX 2019, 37, 121–135, doi:10.14573/altex.1907121.

32. Tukker, A.M.; Wijnolts, F.M.J.J.; de Groot, A.; Westerink, R.H.S.S.; Groot, A. De; Westerink, R.H.S.S.; de Groot, A.; Westerink, R.H.S.S. Human IPSC-Derived Neuronal Models for in Vitro Neurotoxicity Assessment. Neurotoxicology 2018, 67, 215–225, doi:10.1016/j.neuro.2018.06.007.

33. Mack, C.M.; Lin, B.J.; Turner, J.D.; Johnstone, A.F.M.; Burgoon, L.D.; Shafer, T.J. Burst and Principal Components Analyses of MEA Data for 16 Chemicals Describe at Least Three Effects Classes. Neurotoxicology 2014, 40, 75–85, doi:10.1016/j.neuro.2013.11.008.

34. Brown, J.P.; Hall, D.; Frank, C.L.; Wallace, K.; Mundy, W.R.; Shafer, T.J.; Epa, U.S. Evaluation of a Microelectrode Array-Based Assay for Neural Network Ontogeny Using Training Set Chemicals. Toxicol. Sci. 2016, 154, 126–139, doi:10.1093/toxsci/kfw147.

35. Tukker, A.M.; Wijnolts, F.M.J.; de Groot, A.; Westerink, R.H.S. Applicability of HiPSC-Derived Neuronal Cocultures and Rodent Primary Cortical Cultures for In Vitro Seizure Liability Assessment. Toxicol. Sci. 2020, 178, 71–87, doi:10.1093/toxsci/kfaa136.

36. Takahashi, K.; Yamanaka, S. Induction of Pluripotent Stem Cells from Mouse Embryonic and Adult Fibroblast Cultures by Defined Factors. Cell 2006, 126, 663–676, doi:10.1016/j.cell.2006.07.024.

37. Takahashi, K.; Tanabe, K.; Ohnuki, M.; Narita, M.; Ichisaka, T.; Tomoda, K.; Yamanaka, S. Induction of Pluripotent Stem Cells from Adult Human Fibroblasts by Defined Factors. Cell 2007, 131, 861–872, doi:10.1016/j.cell.2007.11.019.

38. Shi, Y.; Inoue, H.; Wu, J.C.; Yamanaka, S. Induced Pluripotent Stem Cell Technology: A Decade of Progress. Nat. Rev. Drug Discov. 2017, 16, 115–130, doi:10.1038/nrd.2016.245.

39. Galiakberova, A.A.; Dashinimaev, E.B. Neural Stem Cells and Methods for Their Generation From Induced Pluripotent Stem Cells in Vitro. 2020, 8, doi:10.3389/fcell.2020.00815.

40. Li, W.; Sun, W.; Zhang, Y.; Wei, W.; Ambasudhan, R.; Xia, P.; Talantova, M.; Lin, T.; Kim, J.; Wang, X.; et al. Rapid Induction and Long-Term Self-Renewal of Primitive Neural Precursors from Human Embryonic Stem Cells by Small Molecule Inhibitors. Proc. Natl. Acad. Sci. U. S. A. 2011, 108, 8299–8304, doi:10.1073/pnas.1014041108.

41. Chambers, S.M.; Fasano, C.A.; Papapetrou, E.P.; Tomishima, M.; Sadelain, M.; Studer, L. Highly Efficient Neural Conversion of Human ES and IPS Cells by Dual Inhibition of SMAD Signaling. Nat. Biotechnol. 2009, 27, 275–280, doi:10.1038/nbt.1529.

42. Shi, Y.; Kirwan, P.; Smith, J.; Robinson, H.P.C.; Livesey, F.J. Human Cerebral Cortex Development from Pluripotent Stem Cells to Functional Excitatory Synapses. Nat. Neurosci. 2012, 15, 477–486, doi:10.1038/nn.3041.

43. Izsak, J.; Seth, H.; Andersson, M.; Vizlin-hodzic, D.; Theiss, S.; Hanse, E.; Ågren, H.; Funa, K.; Illes, S. Robust Generation of Person-Specific, Synchronously Active Neuronal Networks Using Purely Isogenic Human IPSC-3D Neural Aggregate Cultures. Front. Neurosci. 2019, 13, doi:10.3389/fnins.2019.00351.

44. Hyvärinen, T.; Hyysalo, A.; Kapucu, F.E.; Aarnos, L.; Vinogradov, A.; Eglen, S.J.; Ylä-Outinen, L.; Narkilahti, S. Functional Characterization of Human Pluripotent Stem Cell-Derived Cortical Networks Differentiated on Laminin-521 Substrate: Comparison to Rat Cortical Cultures. Sci. Rep. 2019, 9, 1–15, doi:10.1038/s41598-019-53647-8.

45. Paavilainen, T.; Pelkonen, A.; Mäkinen, M.E.; Peltola, M.; Huhtala, H.; Fayuk, D.; Narkilahti, S. Effect of Prolonged Differentiation on Functional Maturation of Human Pluripotent Stem Cell-Derived Neuronal Cultures. Stem Cell Res. 2018, 27, 151–161, doi:10.1016/j.scr.2018.01.018.

46. Gunhanlar, N.; Shpak, G.; van der Kroeg, M.; Gouty-Colomer, L.A.; Munshi, S.T.; Lendemeijer, B.; Ghazvini, M.; Dupont, C.; Hoogendijk, W.J.G.; Gribnau, J.; et al. A Simplified Protocol for Differentiation of Electrophysiologically Mature Neuronal Networks from Human Induced Pluripotent Stem Cells. Mol. Psychiatry 2017, 23, 1336–1344, doi:10.1038/mp.2017.56.

47. de Leeuw, V.C.; van Oostrom, C.T.M.; Wackers, P.F.K.; Pennings, J.L.A.; Hodemaekers, H.M.; Piersma, A.H.; Hessel, E.V.S. Neuronal Differentiation Pathways and Compound-Induced Developmental Neurotoxicity in the Human Neural Progenitor Cell Test (HNPT) Revealed by RNA-Seq. Chemosphere 2022, 304, 135298, doi:10.1016/j.chemosphere.2022.135298.

48. Pistollato, F.; Canovas-Jorda, D.; Zagoura, D.; Price, A. Protocol for the Differentiation of Human Induced Pluripotent Stem Cells into Mixed Cultures of Neurons and Glia for Neurotoxicity Testing. J. Vis. Exp. 2017, 2017, 1–14, doi:10.3791/55702.

49. Schenke, M.; Schjeide, B.M.; Püschel, G.P.; Seeger, B. Analysis of Motor Neurons Differentiated from Human Induced Pluripotent Stem Cells for the Use in Cell-Based Botulinum Neurotoxin Activity Assays. Toxins (Basel). 2020, 12, 1–20, doi:10.3390/toxins12050276.

50. Leeuw, V.C. De; Oostrom, C.T.M. Van; Zwart, E.P.; Heusinkveld, H.J.; Hessel, E.V.S. Prolonged Differentiation of Neuron-Astrocyte Co-Cultures Results in Emergence of Dopaminergic Neurons. 2023.

51. Suzuki, I.K.; Vanderhaeghen, P. Is This a Brain Which i See before Me? Modeling Human Neural Development with Pluripotent Stem Cells. Dev. 2015, 142, 3138–3150, doi:10.1242/dev.120568.

52. Nimtz, L.; Hartmann, J.; Tigges, J.; Masjosthusmann, S.; Schmuck, M.; Keßel, E.; Theiss, S.; Köhrer, K.; Petzsch, P.; Adjaye, J.; et al. Characterization and Application of Electrically Active Neuronal Networks Established from Human Induced Pluripotent Stem Cell-Derived Neural Progenitor Cells for Neurotoxicity Evaluation. Stem Cell Res. 2020, 45, 101761, doi:10.1016/j.scr.2020.101761.

53. Pamies, D.; Barreras, P.; Block, K.; Makri, G.; Kumar, A.; Wiersma, D.; Smirnova, L.; Zang, C.; Bressler, J.; Christian, K.M.; et al. A Human Brain Microphysiological System Derived from Induced Pluripotent Stem Cells to Study Neurological Diseases and Toxicity. ALTEX 2017, 34, 362–376, doi:10.14573/altex.1609122.

54. Pistollato, F.; Louisse, J.; Scelfo, B.; Mennecozzi, M.; Accordi, B.; Basso, G.; Gaspar, J.A.; Zagoura, D.; Barilari, M.; Palosaari, T.; et al. Development of a Pluripotent Stem Cell Derived Neuronal Model to Identify Chemically Induced Pathway Perturbations in Relation to Neurotoxicity: Effects of CREB Pathway Inhibition. Toxicol. Appl. Pharmacol. 2014, 280, 378–388, doi:10.1016/j.taap.2014.08.007.

55. Fritsche, E.; Tigges, J.; Hartmann, J.; Kapr, J.; Serafini, M.M.; Viviani, B. Neural In Vitro Models for Studying Substances Acting on the Central Nervous System. 2020, doi:10.1007/164_2020_367.

56. Logan, S.; Arzua, T.; Canfield, S.G.; Seminary, E.R.; Sison, S.L.; Ebert, A.D.; Bai, X. Studying Human Neurological Disorders Using Induced Pluripotent Stem Cells: From 2D Monolayer to 3D Organoid and Blood Brain Barrier Models. In Comprehensive Physiology; Wiley, 2019; pp. 565–611.

57. Nimtz, L.; Hartmann, J.; Tigges, J.; Masjosthusmann, S.; Schmuck, M.; Keßel, E.; Theiss, S.; Köhrer, K.; Petzsch, P.; Adjaye, J.; et al. Characterization and Application of Electrically Active Neuronal Networks Established from Human Induced Pluripotent Stem Cell-Derived Neural Progenitor Cells for Neurotoxicity Evaluation. Stem Cell Res. 2020, doi:10.1016/j.scr.2020.101761.

58. Abreu, C.M.; Gama, L.; Krasemann, S.; Chesnut, M.; Odwin-Dacosta, S.; Hogberg, H.T.; Hartung, T.; Pamies, D. Microglia Increase Inflammatory Responses in IPSC-Derived Human BrainSpheres. Front. Microbiol. 2018, 9, 1–12, doi:10.3389/fmicb.2018.02766.

59. Pamies, D.; Wiersma, D.; Katt, M.E.; Zhao, L.; Burtscher, J.; Harris, G.; Smirnova, L.; Searson, P.C.; Hartung, T.; Hogberg, H.T. Human Organotypic Brain Model as a Tool to Study Chemical-Induced Dopaminergic Neuronal Toxicity. Neurobiol. Dis. 2022, 169, 105719, doi:10.1016/j.nbd.2022.105719.

60. Pamies, D.; Block, K.; Lau, P.; Gribaldo, L.; Pardo, C.A.; Barreras, P.; Smirnova, L.; Wiersma, D.; Zhao, L.; Harris, G.; et al. Rotenone Exerts Developmental Neurotoxicity in a Human Brain Spheroid Model. Toxicol. Appl. Pharmacol. 2018, doi:10.1016/j.taap.2018.02.003.

61. Nunes, C.; Gorczyca, G.; Mendoza-deGyves, E.; Ponti, J.; Bogni, A.; Carpi, D.; Bal-Price, A.; Pistollato, F. Upscaling Biological Complexity to Boost Neuronal and Oligodendroglia Maturation and Improve in Vitro Developmental Neurotoxicity (DNT) Evaluation. Reprod. Toxicol. 2022, 110, 124–140, doi:10.1016/j.reprotox.2022.03.017.

62. Leite, P.E.C.; Pereira, M.R.; Harris, G.; Pamies, D.; Maria, L.; Granjeiro, J.M.; Hogberg, H.T.; Hartung, T.; Smirnova, L.; Leite, P.E.C.; et al. Suitability of 3D Human Brain Spheroid Models to Distinguish Toxic Effects of Gold and Poly-Lactic Acid Nanoparticles to Assess Biocompatibility for Brain Drug Delivery. Part. Fibre Toxicol. 2019, 16, 1–20, doi:10.1186/s12989-019-0307-3.

63. Kobolak, J.; Teglasi, A.; Bellak, T.; Janstova, Z.; Molnar, K.; Zana, M.; Bock, I.; Laszlo, L.; Dinnyes, A. Human Induced Pluripotent Stem Cell-Derived 3D-Neurospheres Are Suitable for Neurotoxicity Screening. Cells 2020, 9, doi:10.3390/cells9051122.

64. Harris, G.; Hogberg, H.; Hartung, T.; Smirnova, L. 3D Differentiation of LUHMES Cell Line to Study Recovery and Delayed Neurotoxic Effects. Curr. Protoc. Toxicol. 2017, 2017, 1–42, doi:10.1002/cptx.29.

65. Smirnova, L.; Harris, G.; Delp, J.; Valadares, M.; Pamies, D.; Hogberg, H.T.; Waldmann, T.; Leist, M.; Hartung, T. A LUHMES 3D Dopaminergic Neuronal Model for Neurotoxicity Testing Allowing Long-Term Exposure and Cellular Resilience Analysis. Arch. Toxicol. 2016, 90, 2725–2743, doi:10.1007/s00204-015-1637-z.

66. Tigges, J.; Bielec, K.; Brockerhoff, G.; Hildebrandt, B.; Hübenthal, U.; Kapr, J.; Koch, K.; Teichweyde, N.; Wieczorek, D.; Rossi, A.; et al. Academic Application of Good Cell Culture Practice for Induced Pluripotent Stem Cells. ALTEX 2021, 1–19, doi:10.14573/altex.2101221.

67. Bartmann, K.; Hartmann, J.; Kapr, J.; Fritsche, E. Measurement of of Differentiated Human IPSC-Derived Neurospheres Recorded by Microelectrode Arrays (MEA). In; Humana, New York, NY, 2021; pp. 473–488 ISBN 978-1-0716-1637-6.

68. Bartmann, K.; Bendt, F.; Dönmez, A.; Haag, D.; Keßel, E.; Masjosthusmann, S.; Noel, C.; Wu, J.; Zhou, P.; Fritsche, E. A Human IPSC-Based in Vitro Neural Network Formation Assay to Investigate Neurodevelopmental Toxicity of Pesticides. bioRxiv 2023, 1–41, doi:10.1101/2023.01.12.523741.

69. Walter, K.M.; Dach, K.; Hayakawa, K.; Giersiefer, S.; Heuer, H.; Lein, P.J.; Fritsche, E. Ontogenetic Expression of Thyroid Hormone Signaling Genes: An in Vitro and in Vivo Species Comparison. PLoS One 2019, 14, e0221230, doi:10.1371/journal.pone.0221230.

70. Park, D.; Xiang, A.P.; Mao, F.F.; Zhang, L.; Di, C.G.; Liu, X.M.; Shao, Y.; Ma, B.F.; Lee, J.H.; Ha, K.S.; et al. Nestin Is Required for the Proper Self-Renewal of Neural Stem Cells. Stem Cells 2010, 28, 2162–2171, doi:10.1002/stem.541.

71. Sansom, S.N.; Griffiths, D.S.; Faedo, A.; Kleinjan, D.J.; Ruan, Y.; Smith, J.; Van Heyningen, V.; Rubenstein, J.L.; Livesey, F.J. The Level of the Transcription Factor Pax6 Is Essential for Controlling the Balance between Neural Stem Cell Self-Renewal and Neurogenesis. PLoS Genet. 2009, 5, 20–23, doi:10.1371/journal.pgen.1000511.

72. Honegger, P.; Lenoir, D.; Favrod, P. Growth and Differentiation of Aggregating Fetal Brain Cells in a Serum-Free Defined Medium [21]. Nature 1979, 282, 305–308, doi:10.1038/282305a0.

73. Tao, Y.; Zhang, S.C. Neural Subtype Specification from Human Pluripotent Stem Cells. Cell Stem Cell 2016, 19, 573–586, doi:10.1016/j.stem.2016.10.015.

74. Leonzino, M.; Busnelli, M.; Antonucci, F.; Verderio, C.; Mazzanti, M.; Chini, B. The Timing of the Excitatory-to-Inhibitory GABA Switch Is Regulated by the Oxytocin Receptor via KCC2. Cell Rep. 2016, 15, 96–103, doi:10.1016/j.celrep.2016.03.013.

75. Maccioni, R.B.; Cambiazo, V. Role of Microtubule-Associated Proteins in the Control of Microtubule Assembly. Physiol. Rev. 1995, 75, 835–864, doi:10.1152/physrev.1995.75.4.835.

76. Holst, C.B.; Brøchner, C.B.; Vitting-Seerup, K.; Møllgård, K. Astrogliogenesis in Human Fetal Brain: Complex Spatiotemporal Immunoreactivity Patterns of GFAP, S100, AQP4 and YKL-40. J. Anat. 2019, 235, 590–615, doi:10.1111/joa.12948.

77. Rowley, N.M.; Madsen, K.K.; Schousboe, A.; Steve White, H. Glutamate and GABA Synthesis, Release, Transport and Metabolism as Targets for Seizure Control. Neurochem. Int. 2012, 61, 546–558, doi:10.1016/j.neuint.2012.02.013.

78. Walls, A.B.; Nilsen, L.H.; Eyjolfsson, E.M.; Vestergaard, H.T.; Hansen, S.L.; Schousboe, A.; Sonnewald, U.; Waagepetersen, H.S. GAD65 Is Essential for Synthesis of GABA Destined for Tonic Inhibition Regulating Epileptiform Activity. J. Neurochem. 2010, 115, 1398–1408, doi:10.1111/j.1471-4159.2010.07043.x.

79. Niciu, M.J.; Kelmendi, B.; Sanacora, G. Overview of Glutamatergic Neurotransmission in the Nervous System. Pharmacol. Biochem. Behav. 2012, 100, 656–664, doi:10.1016/j.pbb.2011.08.008.

80. Bormann, J. The “ABC” of GABA Receptors. Trends Pharmacol. Sci. 2000, 21, 16–19, doi:10.1016/S0165-6147(99)01413-3.

81. Popova, E. Ionotropic GABA Receptors and Distal Retinal ON and OFF Responses. Scientifica (Cairo). 2014, 2014, 1–23, doi:10.1155/2014/149187.

82. Fan, L.; Tan, L.; Chen, Z.; Qi, J.; Nie, F.; Luo, Z.; Cheng, J.; Wang, S. Haloperidol Bound D2 Dopamine Receptor Structure Inspired the Discovery of Subtype Selective Ligands. Nat. Commun. 2020, 11, 1–11, doi:10.1038/s41467-020-14884-y.

83. Martel, J.C.; Gatti McArthur, S. Dopamine Receptor Subtypes, Physiology and Pharmacology: New Ligands and Concepts in Schizophrenia. Front. Pharmacol. 2020, 11, 1–17, doi:10.3389/fphar.2020.01003.

84. Sagarduy, A.; Llorente, J.; Miguelez, C.; Morera-Herreras, T.; Ruiz-Ortega, J.A.; Ugedo, L. Buspirone Requires the Intact Nigrostriatal Pathway to Reduce the Activity of the Subthalamic Nucleus via 5-HT1A Receptors. Exp. Neurol. 2016, 277, 35–45, doi:10.1016/j.expneurol.2015.12.005.

85. Qujeq, D.; Roushan, T.; Norouzy, A.; Habibi-Rezaei, M.; Mehdinejad-Shani, M. Effects of Dichlorvos and Carbaryl on the Activity of Free and Immobilized Acetylcholinesterase. Toxicol. Ind. Health 2012, 28, 291–295, doi:10.1177/0748233711410907.

86. Smulders, C.J.G.M.; Bueters, T.J.H.; Van Kleef, R.G.D.M.; Vijverberg, H.P.M. Selective Effects of Carbamate Pesticides on Rat Neuronal Nicotinic Acetylcholine Receptors and Rat Brain Acetylcholinesterase. Toxicol. Appl. Pharmacol. 2003, 193, 139–146, doi:10.1016/j.taap.2003.07.011.

87. Nagata, K.; Huang, C.S.; Song, J.H.; Narahashi, T. Direct Actions of Anticholinesterases on the Neuronal Nicotinic Acetylcholine Receptor Channels. Brain Res. 1997, 769, 211–218, doi:10.1016/S0006-8993(97)00707-5.

88. Xu, X.; Sepich, C.; Lukas, R.J.; Zhu, G.; Chang, Y. Emamectin Is a Non-Selective Allosteric Activator of Nicotinic Acetylcholine Receptors and GABAA/C Receptors. Biochem. Biophys. Res. Commun. 2016, 473, 795–800, doi:10.1016/j.bbrc.2016.03.097.

89. Patterson, T.A.; Eppler, B.; Dawson, R. Attenuation of Trimethyltin-Evoked Glutamate (GLU) Efflux from Rat Cortical and Hippocampal Slices. Neurotoxicol. Teratol. 1996, 18, 697–702, doi:10.1016/S0892-0362(96)00132-8.

90. Bal-Price, A.K.; Suñol, C.; Weiss, D.G.; van Vliet, E.; Westerink, R.H.S.; Costa, L.G. Application of in Vitro Neurotoxicity Testing for Regulatory Purposes: Symposium III Summary and Research Needs. Neurotoxicology 2008, 29, 520–531, doi:10.1016/j.neuro.2008.02.008.

91. Coecke, S.; Eskes, C.; Gartlon, J.; Kinsner, A.; Price, A.; Van Vliet, E.; Prieto, P.; Boveri, M.; Bremer, S.; Adler, S.; et al. The Value of Alternative Testing for Neurotoxicity in the Context of Regulatory Needs. Environ. Toxicol. Pharmacol. 2006, 21, 153–167, doi:10.1016/j.etap.2005.07.006.

92. Tate, K.; Kirk, B.; Tseng, A.; Ulffers, A.; Litwa, K. Effects of the Selective Serotonin Reuptake Inhibitor Fluoxetine on Developing Neural Circuits in a Model of the Human Fetal Cortex. Int. J. Mol. Sci. 2021, 22, doi:10.3390/ijms221910457.

93. Tukker, A.M.; Bouwman, L.M.S.; van Kleef, R.G.D.M.; Hendriks, H.S.; Legler, J.; Westerink, R.H.S. Perfluorooctane Sulfonate (PFOS) and Perfluorooctanoate (PFOA) Acutely Affect Human Α1β2γ2L GABAA Receptor and Spontaneous Neuronal Network Function in Vitro. Sci. Rep. 2020, 10, 1–14, doi:10.1038/s41598-020-62152-2.

94. Sirenko, O.; Parham, F.; Dea, S.; Sodhi, N.; Biesmans, S.; Mora-Castilla, S.; Ryan, K.; Behl, M.; Chandy, G.; Crittenden, C.; et al. Functional and Mechanistic Neurotoxicity Profiling Using Human IPSC-Derived Neural 3D Cultures. Toxicol. Sci. 2019, 167, 249–257, doi:10.1093/toxsci/kfy218.

95. van Melis, L.V.J.; Heusinkveld, H.J.; Langendoen, C.; Peters, A.; Westerink, R.H.S. Organophosphate Insecticides Disturb Neuronal Network Development and Function via Non-AChE Mediated Mechanisms. Neurotoxicology 2022, 94, 35–45, doi:10.1016/j.neuro.2022.11.002.

96. Loser, D.; Grillberger, K.; Hinojosa, M.G.; Blum, J.; Haufe, Y.; Danker, T.; Johansson, Y.; Möller, C.; Nicke, A.; Bennekou, S.H.; et al. Acute Effects of the Imidacloprid Metabolite Desnitro-Imidacloprid on Human NACh Receptors Relevant for Neuronal Signaling. Arch. Toxicol. 2021, 95, 3695–3716, doi:10.1007/s00204-021-03168-z.

97. Kosodo, Y.; Röper, K.; Haubensak, W.; Marzesco, A.M.; Corbeil, D.; Huttner, W.B. Asymmetric Distribution of the Apical Plasma Membrane during Neurogenic Divisions of Mamalian Neuroepithelial Cells. EMBO J. 2004, 23, 2314– 2324, doi:10.1038/sj.emboj.7600223.

98. Bayatti, N.; Sarma, S.; Shaw, C.; Eyre, J.A.; Vouyiouklis, D.A.; Lindsay, S.; Clowry, G.J. Progressive Loss of PAX6, TBR2, NEUROD and TBR1 MRNA Gradients Correlates with Translocation of EMX2 to the Cortical Plate during Human Cortical Development. Eur. J. Neurosci. 2008, 28, 1449–1456, doi:10.1111/j.1460-9568.2008.06475.x.

99. Estivill-Torrus, G.; Pearson, H.; van Heyningen, V.; Price, D.J.; Rashbass, P. Pax6 Is Required to Regulate the Cell Cycle and the Rate of Progression from Symmetrical to Asymmetrical Division in Mammalian Cortical Progenitors. Development 2002, 129, 455–466, doi:10.1242/dev.129.2.455.

100. Paavilainen, T.; Pelkonen, A.; Mäkinen, M.E.-L.E.L.L.; Peltola, M.; Huhtala, H.; Fayuk, D.; Narkilahti, S.; Narkilathi, S. Effect of Prolonged Differentiation on Functional Maturation of Human Pluripotent Stem Cell-Derived Neuronal Cultures. Stem Cell Res. 2018, 27, 151–161, doi:10.1016/j.scr.2018.01.018.

101. Togo, K.; Fukusumi, H.; Shofuda, T.; Ohnishi, H.; Yamazaki, H.; Hayashi, M.K.; Kawasaki, N.; Takei, N.; Nakazawa, T.; Saito, Y.; et al. Postsynaptic Structure Formation of Human IPS Cell-Derived Neurons Takes Longer than Presynaptic Formation during Neural Differentiation in Vitro. Mol. Brain 2021, 14, 1–17, doi:10.1186/s13041-021-00851-1.

102. Reemst, K.; Noctor, S.C.; Lucassen, P.J.; Hol, E.M. The Indispensable Roles of Microglia and Astrocytes during Brain Development. Front. Hum. Neurosci. 2016, 10, 1–28, doi:10.3389/fnhum.2016.00566.

103. Mahmoud, S.; Gharagozloo, M.; Simard, C.; Gris, D. Astrocytes Maintain Glutamate Homeostasis in the CNS by Controlling the Balance between Glutamate Uptake and Release. Cells 2019, 8, 184, doi:10.3390/cells8020184.

104. Oyanagi, K.; Tashiro, T.; Negishi, T. Cell-Type-Specific and Differentiation-Status-Dependent Variations in Cytotoxicity of Tributyltin in Cultured Rat Cerebral Neurons and Astrocytes. J. Toxicol. Sci. 2015, 40, 459–468, doi:10.2131/jts.40.459.

105. Laurenza, I.; Pallocca, G.; Mennecozzi, M.; Scelfo, B.; Pamies, D.; Bal-Price, A. A Human Pluripotent Carcinoma Stem Cell-Based Model for in Vitro Developmental Neurotoxicity Testing: Effects of Methylmercury, Lead and Aluminum Evaluated by Gene Expression Studies. Int. J. Dev. Neurosci. 2013, 31, 679–691, doi:10.1016/j.ijdevneu.2013.03.002.

106. Pei, Y.; Peng, J.; Behl, M.; Sipes, N.S.; Shockley, K.R.; Rao, M.S.; Tice, R.R.; Zeng, X. Comparative Neurotoxicity Screening in Human IPSC-Derived Neural Stem Cells, Neurons and Astrocytes. Brain Res. 2016, 1638, 57–73, doi:10.1016/j.brainres.2015.07.048.

107. Wang, Y.; Zhao, F.; Liao, Y.; Jin, Y.; Sun, G. Effects of Arsenite in Astrocytes on Neuronal Signaling Transduction. Toxicology 2013, 303, 43–53, doi:10.1016/j.tox.2012.10.024.

108. Guttenplan, K.A.; Weigel, M.K.; Prakash, P.; Wijewardhane, P.R.; Hasel, P.; Rufen-Blanchette, U.; Münch, A.E.; Blum, J.A.; Fine, J.; Neal, M.C.; et al. Neurotoxic Reactive Astrocytes Induce Cell Death via Saturated Lipids. Nature 2021, 599, 102–107, doi:10.1038/s41586-021-03960-y.

109. Stary, C.M.; Sun, X.; Giffard, R.G. Astrocytes Protect against Isoflurane Neurotoxicity by Buffering Pro-Brain– Derived Neurotrophic Factor. Anesthesiology 2015, 123, 810–819, doi:10.1097/ALN.0000000000000824.

110. Brüll, M.; Spreng, A.S.; Gutbier, S.; Loser, D.; Krebs, A.; Reich, M.; Kraushaar, U.; Britschgi, M.; Patsch, C.; Leist, M. Incorporation of Stem Cell-Derived Astrocytes into Neuronal Organoids to Allow Neuro-Glial Interactions in Toxicological Studies. ALTEX 2020, 37, 409–428, doi:10.14573/altex.1911111.

111. Crofton, K.M.; Bassan, A.; Behl, M.; Chushak, Y.G.; Fritsche, E.; Gearhart, J.M.; Marty, M.S.; Mumtaz, M.; Pavan, M.; Ruiz, P.; et al. Current Status and Future Directions for a Neurotoxicity Hazard Assessment Framework That Integrates in Silico Approaches. Comput. Toxicol. 2022, 22, 100223, doi:10.1016/j.comtox.2022.100223.

112. Betarbet, R.; Sherer, T.B.; Mackenzie, G.; Garcia-osuna, M.; Panov, A. V; Greenamyre, J.T. Chronic Systemic Pesticide Exposure Produces Pd Symptoms Betarbet. Nat. Neurosci. 2000, 26, 1301–1306.

113. Jokanović, M. Neurotoxic Effects of Organophosphorus Pesticides and Possible Association with Neurodegenerative Diseases in Man: A Review. Toxicology 2018, 410, 125–131, doi:10.1016/j.tox.2018.09.009.

114. Tukker, A.M.; Wijnolts, F.M.J.; de Groot, A.; Westerink, R.H.S. Human IPSC-Derived Neuronal Models for in Vitro Neurotoxicity Assessment. Neurotoxicology 2018, 67, 215–225, doi:10.1016/j.neuro.2018.06.007.

115. Pauly, M.G.; Krajka, V.; Stengel, F.; Seibler, P.; Klein, C.; Capetian, P. Adherent vs. Free-Floating Neural Induction by Dual SMAD Inhibition for Neurosphere Cultures Derived from Human Induced Pluripotent Stem Cells. Front. Cell Dev. Biol. 2018, 6, doi:10.3389/fcell.2018.00003.

116. Nadadhur, A.G.; Leferink, P.S.; Holmes, D.; Hinz, L.; Cornelissen-Steijger, P.; Gasparotto, L.; Heine, V.M. Patterning Factors during Neural Progenitor Induction Determine Regional Identity and Differentiation Potential in Vitro. Stem Cell Res. 2018, 32, 25–34, doi:10.1016/j.scr.2018.08.017.

117. McCready, F.P.; Gordillo-Sampedro, S.; Pradeepan, K.; Martinez-Trujillo, J.; Ellis, J. Multielectrode Arrays for Functional Phenotyping of Neurons from Induced Pluripotent Stem Cell Models of Neurodevelopmental Disorders. Biology (Basel). 2022, 11, doi:10.3390/biology11020316.

118. Tanabe, Y.; Masu, M.; Ishii, T.; Shigemoto, R.; Nakanishi, S. A Family of Metabotropic Glutamate Receptors. Neuron 1992, 8, 169–179, doi:10.1016/0896-6273(92)90118-W.

119. Bodzęta, A.; Scheefhals, N.; MacGillavry, H.D. Membrane Trafficking and Positioning of MGluRs at Presynaptic and Postsynaptic Sites of Excitatory Synapses. Neuropharmacology 2021, 200, doi:10.1016/j.neuropharm.2021.108799.

120. Panatier, A.; Poulain, D.A.; Oliet, S.H.R. Regulation of Transmitter Release by High-Affinity Group III MGluRs in the Supraoptic Nucleus of the Rat Hypothalamus. Neuropharmacology 2004, 47, 333–341, doi:10.1016/j.neuropharm.2004.05.003.

121. Mateo, Z.; Porter, J.T. Group II Metabotropic Glutamate Receptors Inhibit Glutamate Release at Thalamocortical Synapses in the Developing Somatosensory Cortex. Neuroscience 2007, 146, 1062–1072, doi:10.1016/j.neuroscience.2007.02.053.

122. Bocchio, M.; Lukacs, I.P.; Stacey, R.; Plaha, P.; Apostolopoulos, V.; Livermore, L.; Sen, A.; Ansorge, O.; Gillies, M.J.; Somogyi, P.; et al. Group II Metabotropic Glutamate Receptors Mediate Presynaptic Inhibition of Excitatory Transmission in Pyramidal Neurons of the Human Cerebral Cortex. Front. Cell. Neurosci. 2019, 12, doi:10.3389/fncel.2018.00508.

123. Van Den Pol, A.N.; Gao, X.B.; Patrylo, P.R.; Ghosh, P.K.; Obrietan, K. Glutamate Inhibits GABA Excitatory Activity in Developing Neurons. J. Neurosci. 1998, 18, 10749–10761, doi:10.1523/jneurosci.18-24-10749.1998.

124. Görtz, P.; Henning, U.; Theiss, S.; Lange-Asschenfeldt, C. Effect Fingerprints of Antipsychotic Drugs on Neural Networks in Vitro. J. Neural Transm. 2019, 126, 1363–1371, doi:10.1007/s00702-019-02050-8.

125. Gemperle, A.Y.; Enz, A.; Pozza, M.F.; Lüthi, A.; Olpe, H.R. Effects of Clozapine, Haloperidol and Iloperidone on Neurotransmission and Synaptic Plasticity in Prefrontal Cortex and Their Accumulation in Brain Tissue: An in Vitro Study. Neuroscience 2003, 117, 681–695, doi:10.1016/S0306-4522(02)00769-8.

126. Chen, W.; Zhu, F.; Guo, J.; Sheng, J.; Li, W.; Zhao, X.; Wang, G.; Li, K. Chronic Haloperidol Increases Voltage-Gated Na+ Currents in Mouse Cortical Neurons. Biochem. Biophys. Res. Commun. 2014, 450, 55–60, doi:10.1016/j.bbrc.2014.05.081.

127. Dzyubenko, E.; Juckel, G.; Faissner, A. The Antipsychotic Drugs Olanzapine and Haloperidol Modify Network Connectivity and Spontaneous Activity of Neural Networks in Vitro. Sci. Rep. 2017, 7, 1–13, doi:10.1038/s41598-017-11944-0.

128. Valdivia, P.; Martin, M.; LeFew, W.R.; Ross, J.; Houck, K.A.; Shafer, T.J. Multi-Well Microelectrode Array Recordings Detect Neuroactivity of ToxCast Compounds. Neurotoxicology 2014, 44, 204–217, doi:10.1016/j.neuro.2014.06.012.

129. Smulders, C.J.G.M.; Van Kleef, R.G.D.M.; de Groot, A.; Gotti, C.; Vijverberg, H.P.M. A Noncompetitive, Sequential Mechanism for Inhibition of Rat α 4β2 Neuronal Nicotinic Acetylcholine Receptors by Carbamate Pesticides. Toxicol. Sci. 2004, 82, 219–227, doi:10.1093/toxsci/kfh261.

130. McConnell, E.R.; McClain, M.A.; Ross, J.; LeFew, W.R.; Shafer, T.J. Evaluation of Multi-Well Microelectrode Arrays for Neurotoxicity Screening Using a Chemical Training Set. Neurotoxicology 2012, 33, 1048–1057, doi:10.1016/j.neuro.2012.05.001.

131. Defranchi, E.; Novellino, A.; Whelan, M.; Vogel, S.; Ramirez, T.; van Ravenzwaay, B.; Landsiedel, R. Feasibility Assessment of Micro-Electrode Chip Assay as a Method of Detecting Neurotoxicity in Vitro. Front. Neuroeng. 2011, 4, 1–12, doi:10.3389/fneng.2011.00006.

132. Alloisio, S.; Nobile, M.; Novellino, A. Multiparametric Characterisation of Neuronal Network Activity for in Vitro Agrochemical Neurotoxicity Assessment. Neurotoxicology 2015, 48, 152–165, doi:10.1016/j.neuro.2015.03.013.

133. Dingemans, M.M.L.; Schütte, M.G.; Wiersma, D.M.M.; de Groot, A.; van Kleef, R.G.D.M.; Wijnolts, F.M.J.; Westerink, R.H.S. Chronic 14-Day Exposure to Insecticides or Methylmercury Modulates Neuronal Activity in Primary Rat Cortical Cultures. Neurotoxicology 2016, 57, 194–202, doi:10.1016/j.neuro.2016.10.002.

134. Kocaturk, S.; Guven, E.B.; Shah, F.; Tepper, J.M.; Assous, M. Cholinergic Control of Striatal GABAergic Microcircuits. Cell Rep. 2022, 41, doi:10.1016/j.celrep.2022.111531.

135. Schvartz, D.; González-Ruiz, V.; Walter, N.; Antinori, P.; Jeanneret, F.; Tonoli, D.; Boccard, J.; Zurich, M.G.; Rudaz, S.; Monnet-Tschudi, F.; et al. Protein Pathway Analysis to Study Development-Dependent Effects of Acute and Repeated Trimethyltin (TMT) Treatments in 3D Rat Brain Cell Cultures. Toxicol. Vitr. 2019, 60, 281–292, doi:10.1016/j.tiv.2019.05.020.

136. Brock, T.O.; O’Callaghan, J.P. Quantitative Changes in the Synaptic Vesicle Proteins Synapsin I and P38 and the Astrocyte-Specific Protein Glial Fibrillary Acidic Protein Are Associated with Chemical-Induced Injury to the Rat Central Nervous System. J. Neurosci. 1987, 7, 931–942, doi:10.1523/jneurosci.07-04-00931.1987.

137. Amin, H.; Marinaro, F.; Tonelli, D.D.P.; Berdondini, L. Developmental Excitatory-to-Inhibitory GABA-Polarity Switch Is Disrupted in 22q11.2 Deletion Syndrome: A Potential Target for Clinical Therapeutics. Sci. Rep. 2017, 7, 1–18, doi:10.1038/s41598-017-15793-9.

138. Deidda, G.; Parrini, M.; Naskar, S.; Bozarth, I.F.; Contestabile, A.; Cancedda, L. Reversing Excitatory GABA A R Signaling Restores Synaptic Plasticity and Memory in a Mouse Model of Down Syndrome. Nat. Med. 2015, 21, 318–326, doi:10.1038/nm.3827.

139. He, Q.; Nomura, T.; Xu, J.; Contractor, A. The Developmental Switch in GABA Polarity Is Delayed in Fragile X Mice. J. Neurosci. 2014, 34, 446–450, doi:10.1523/JNEUROSCI.4447-13.2014.

140. Braat, S.; Kooy, R.F. The GABAA Receptor as a Therapeutic Target for Neurodevelopmental Disorders. Neuron 2015, 86, 1119–1130, doi:10.1016/j.neuron.2015.03.042.

141. Modafferi, S.; Zhong, X.; Kleensang, A.; Murata, Y.; Fagiani, F.; Pamies, D.; Hogberg, H.T.; Calabrese, V.; Lachman, H.; Hartung, T.; et al. Gene–Environment Interactions in Developmental Neurotoxicity: A Case Study of Synergy between Chlorpyrifos and Chd8 Knockout in Human Brainspheres. Environ. Health Perspect. 2021, 129, doi:10.1289/EHP8580.

142. Zhong, X.; Harris, G.; Smirnova, L.; Zufferey, V.; Sá, R. de C. da S. e; Baldino Russo, F.; Baleeiro Beltrao Braga, P.C.; Chesnut, M.; Zurich, M.-G.; Hogberg, H.T.; et al. Antidepressant Paroxetine Exerts Developmental Neurotoxicity in an IPSC-Derived 3D Human Brain Model. Front. Cell. Neurosci. 2020, 14, doi:10.3389/fncel.2020.00025.

143. Emílio, P.; Leite, C.; Pereira, M.R.; Harris, G.; Pamies, D.; Maria, L.; Granjeiro, J.M.; Hogberg, H.T.; Hartung, T.; Smirnova, L. Suitability of 3D Human Brain Spheroid Models to Distinguish Toxic Effects of Gold and Poly-Lactic Acid Nanoparticles to Assess Biocompatibility for Brain Drug Delivery. 2019, 1–20.

144. Rockley, K.L.; Roberts, R.A.; Morton, M.J. Innovative Models for: In Vitro Detection of Seizure. Toxicol. Res. (Camb). 2019, 8, 784–788, doi:10.1039/c9tx00210c.

145. Tukker, A.M.; Westerink, R.H.S. Novel Test Strategies for in Vitro Seizure Liability Assessment. Expert Opin. Drug Metab. Toxicol. 2021, 17, 923–936, doi:10.1080/17425255.2021.1876026.

146. Blum, J.; Masjosthusmann, S.; Bartmann, K.; Bendt, F.; Dolde, X.; Dönmez, A.; Förster, N.; Holzer, A.-K.; Hübenthal, U.; Keßel, H.E.; et al. Establishment of a Human Cell-Based in Vitro Battery to Assess Developmental Neurotoxicity Hazard of Chemicals. Chemosphere 2023, 311, 137035, doi:10.1016/j.chemosphere.2022.137035.

147. Crofton, K.M.; Mundy, W.R. External Scientific Report on the Interpretation of Data from the Developmental Neurotoxicity In Vitro Testing Assays for Use in Integrated Approaches for Testing and Assessment. EFSA Support. Publ. 2021, 18, doi:10.2903/sp.efsa.2021.en-6924.

