## Supplementary Material for "Molecular and functional characterization of different BrainSphere models for use in neurotoxicity testing on microelectrode arrays"

**Supplementary Table S1.** Coatings used for neural differentiation of BrainSpheres generated with different neural induction protocols.

| Neural induction protocol | Differentiation medium | coating | Order number and supplier |
|---|---|---|---|
| 2D-NIM | CINDA+ | 0.1 mg/mL PLO[1] / 50 µg/mL laminin | PLO: #P4957, Sigma-Aldrich, USA<br>laminin: #LN521, Biolamina, Sweden |
| | Electro | 0.1 % PEI[2] / 50 µg/mL laminin | PEI: #181978, Sigma-Aldrich, USA<br>laminin: #LN521, Biolamina, Sweden |
| GNEIB | CINDA+ | 25 µg/mL laminin | laminin: #LN521, Biolamina, Sweden |
| | Electro | 0.015 mg/mL PLO[1] / 50 µg/mL laminin | PLO: #P4957, Sigma-Aldrich, USA<br>laminin: #LN521, Biolamina, Sweden |
| Stemdiff | CINDA+ | 50 µg/mL laminin | laminin: #LN521, Biolamina, Sweden |
| | Electro | 0.1 mg/mL PLO[1] / 50 µg/mL laminin | PLO: #P4957, Sigma-Aldrich, USA<br>laminin: #LN521, Biolamina, Sweden |

[1]PLO: poly-L-ornithine; [2]PEI: polyethyleneimine

**Supplementary Table S2.** Information about used compounds and their solvents.

| Compound | CASRN | Order number and supplier | solvent | effect |
|---|---|---|---|---|
| Bicuculline | 485-49-4 | #0130, Tocris, Great Britain | DMSO | $GABA_A$ antagonist |
| Buspirone hydrochloride | 33386-08-2 | #HY-B1115, MedChemExpress, USA | DMSO | 5-HT1A agonist |
| Carbaryl | 63-25-2 | #32055, Sigma-Aldrich, USA | DMSO | inhibits acetylcholinesterase |
| DL-2-amino-5-phosphonovaleric acid (AP5) | 76326-31-3 | #SC-201503, Santa Cruz Biotechnology Inc., USA | $H_2O$ | NMDA antagonist |
| Emamectin benzoate | 155569-91-8 | #31733, Merck, Germany | DMSO | $GABA_A$ and $GABA_C$ agonist |
| Haloperidol | 52-86-8 | #HY-14538, MedChemExpress, USA | DMSO | D2 antagonist |
| L-glutamate | 6106-04-3 | #49621, Sigma-Aldrich, USA | $H_2O$ | Neurotransmitter |
| NBQX disodium salt (NBQX) | 479347-86-9 | #SC-222048, Santa Cruz Biotechnology Inc., USA | DMSO | AMPA antagonist |

| | | | | |
|---|---|---|---|---|
| Picrotoxin | 124-87-8 | #SC-202765, Santa Cruz Biotechnology Inc., USA | DMSO | $GABA_A$ and $GABA_C$ antagonist |
| Trimethyltin chloride (TMT) | 1066-45-1 | #146498, Merck, Germany | $H_2O$ | Increases glutamate release |
| γ-Aminobutyric acid (GABA) | 56-12-2 | #A5835, Sigma-Aldrich, USA | $H_2O$ | Neurotransmitter |

**Supplementary Table S3.** Primer sequences.

| **Gene** | **Forward primer sequence (5' → 3')** | **Reverse primer sequence (5' → 3')** |
|---|---|---|
| ACHE | GTCCAGACTAACGTACTGCT | ATGCGATACTGGGCCAAC |
| ACTB | CAGGAAGTCCCTTGCCATCC | ACCAAAAGCCTTCATACATCTCA |
| AQP4 | TGGACAGAAGACATACTCATAAAGG | GGTGCCAGCATGAATCCC |
| CHAT | GCCCATAGTATTGCTTCATGC | ACCAGCTGAGGTTTGCAG |
| CHRNA4 | CTCGCGGGCTCTAGATG | CCACAGGAGAAGACGAACC |
| DLG4 | CCAGAAGAGTACAGCCGATTC | CTTGGTCTTGTCGTAATCAAACAG |
| DRD2 | ACGGCGAGCATCCTGAACTT | GCCGGGTTGGCAATGATGCA |
| EN1 | CTGGGTGTACTGCACACGTTAT | TACTCGCTCTCGTCTTTGTCCT |
| FOXG1 | TCCATCCGCCACAATCTGTC | CCCAGCGAGTTCTGAGTCAA |
| GABRA1 | ACAACACTTACGCTCCAACA | TTTCGGGCTTGACCTCTTTAG |
| GAD1 | GACCCCAATACCACTAACCTG | GCTGTTGGTCCTTTGCAAG |
| GFAP | CACTGTGAGGCAGAAGCTC | CCTCCAGCGACTCAATCTTC |
| GLS | CAGTTCCAAGATCATTAACAGCA | CGATTTGTGGGGTGTGTCT |
| GRIA1 | CTTAATCGAGTTCTGCTACAAATCC | GTATGGCTTCGTTGATGGATTG |
| GRIN1 | CTCCTGGAAGATTCAGCTCAA | GTGGATGGCTAACTAGGATGG |
| HOXA2 | CACCACGTCTACGGGCAAGAAC | AGCTGCTGATGCTGACTTCTGA |
| HTR1A | TGTCGCTTCTCACAAACTCTC | CTTCTCCTCTTTCTCTCTGCTC |
| LMX1A | CGGGTCATCTTGGACAGGTTT | CAGAAGCAGCTCAGGTGGTA |
| MAP2 | TGCCTCAGAACAGACTGTCAC | AAGGCTCAGCTGTAGAGGGA |
| S100B | CACATTCGCCGTCTCCATC | CACAAGCTGAAGAAATCCGAAC |
| SLC12A2 | ACAAAGTTGAGGAAGAGGATGGC | CCTGATCTGCCGGTATGTCTTGG |
| SLC12A5 | CCGATGACAATTACCCATGGA | CTGTCTACATCAGCTCCGTTG |

| | | |
|---|---|---|
| SLC17A6 | CCTCTACCCTAAATATGCTAATTCCA | TTGCTCCATATCCCATGACAT |
| SLC17A7 | CAGAAAGCCCAGTTCAGC | ACAATAGCAAAGCCGAAAACTC |
| SYN1 | GTGCTGCTGGTCATCGAC | CCACAAGGTTGAGATCAGAGAA |
| TH | GGTCTCTAGATGGTGGATTTTGG | TGCTAAACCTGCTCTTCTCC |
| TPH1 | TGTCTCACATATTGAGTGCAGT | CCACAATTGGAAGATGTCTCC |

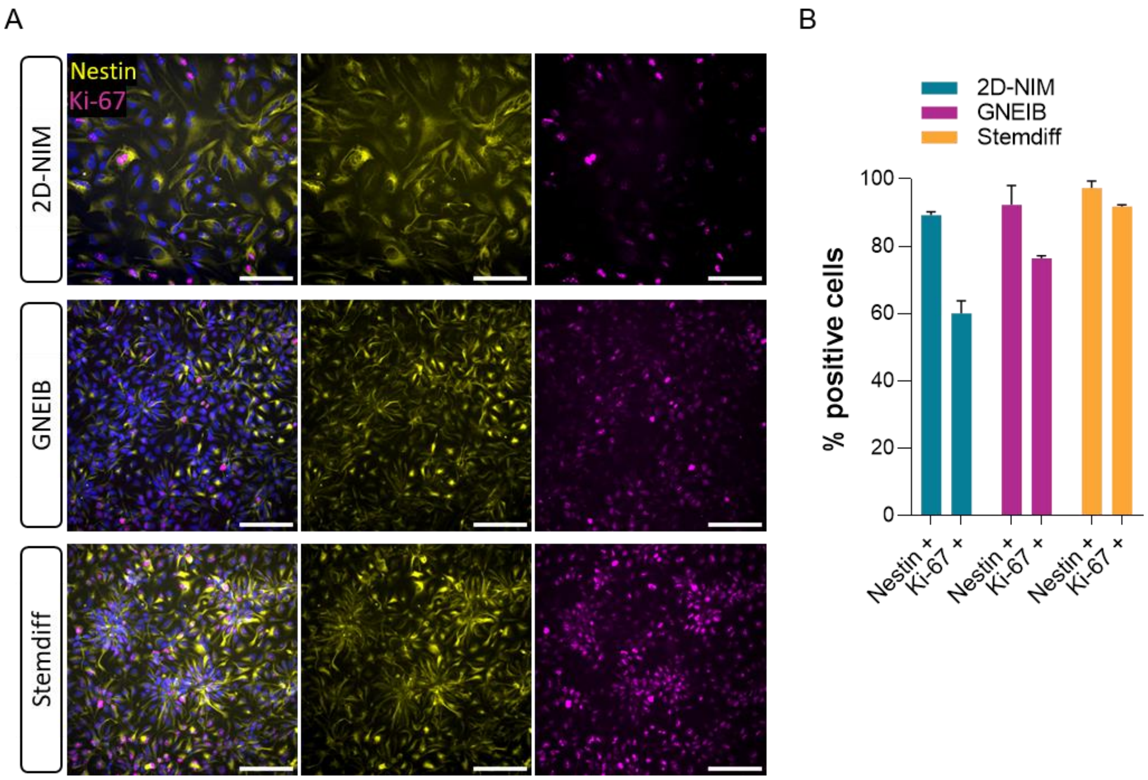


**Supplementary Figure S1.** Immunocytochemistry staining of human induced neural progenitor cells (hiNPCs). A) After thawing, the hiNPCs were replated onto microscopy slides and stained for the neural progenitor marker nestin (yellow) and the proliferation marker Ki-67 (magenta). The nuclei were counterstained with Hoechst 34580 (blue). Representative images are shown. Scale bar =100 µm. B) Quantification of hiNPCs stained for the NPC marker nestin or the proliferation marker Ki-67. Data are represented as mean ± SEM of two independent experiments.

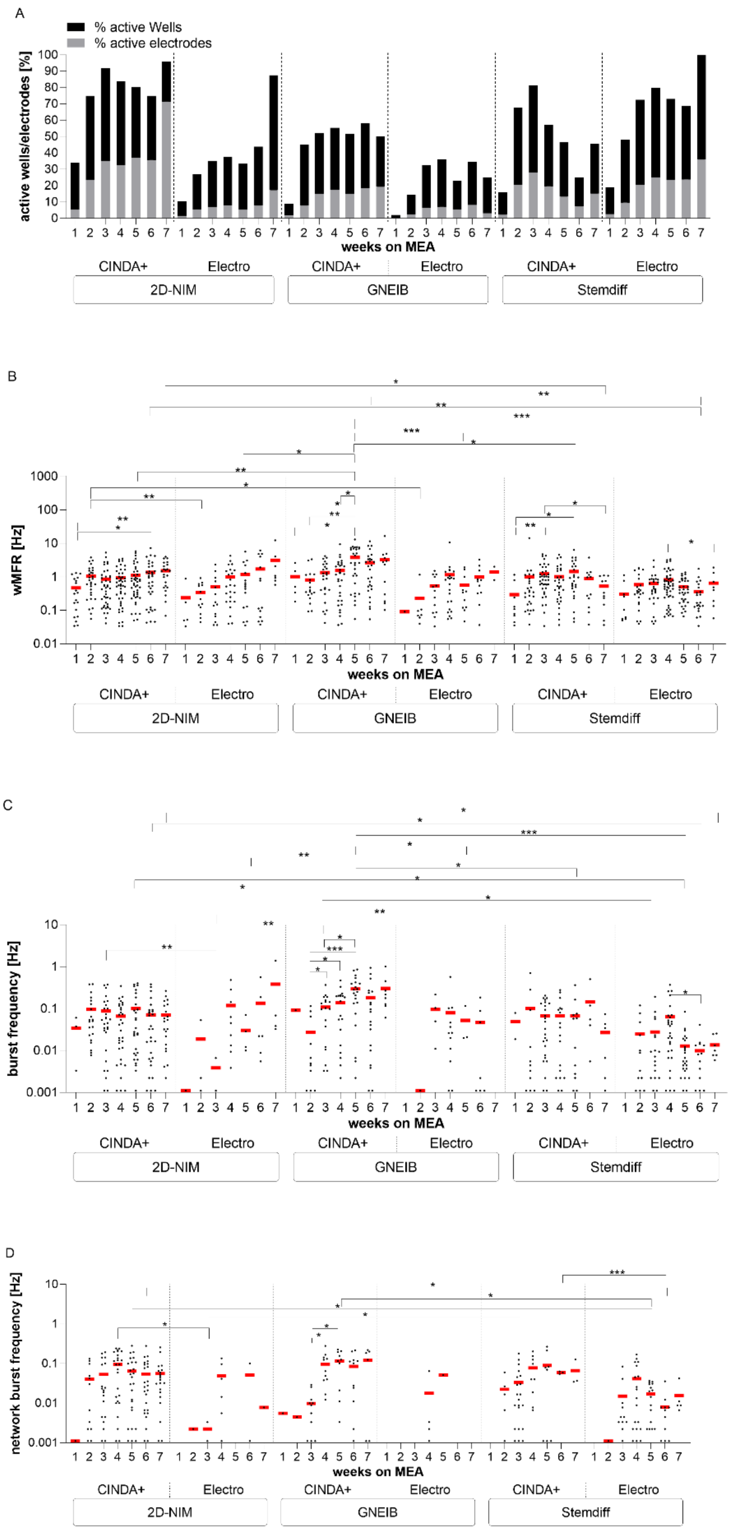


**Supplementary Figure S2.** Comparison of electrical activity of BrainSpheres (without 3D differentiation) for 7 weeks on microelectrode arrays (MEA). The neuronal functionality was measured twice per week and the parameters active wells and active electrodes (A), weighted mean firing rate (wMFR, B), burst frequency (C), and network burst frequency (D) were analyzed. Each dot represents the mean of one well containing 8 electrodes and the red bar represents the mean of all wells resulting from three independent MEA experiments each ($p \leq 0.05$).

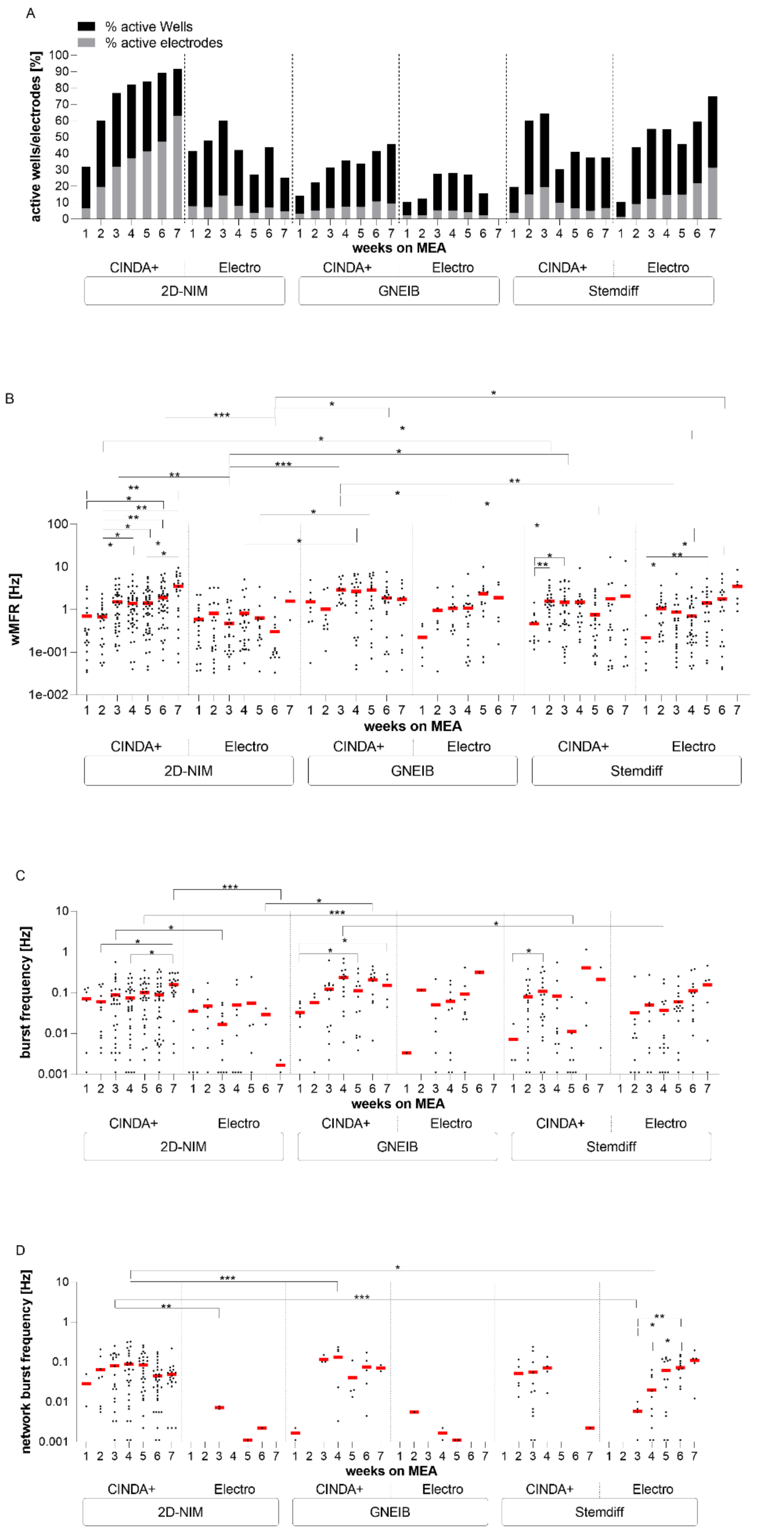


**Supplementary Figure S3.** Comparison of electrical activity of 1 week 3D differentiated BrainSpheres for 7 weeks on microelectrode arrays (MEA). The neuronal functionality was measured twice per week and the parameters active wells and active electrodes (A), weighted mean firing rate (wMFR, B), burst frequency (C), and network burst frequency (D) were analyzed. Each dot represents the mean of one well containing 8 electrodes and the red bar represents the mean of all wells resulting from three independent MEA experiments each ($p \leq 0.05$).

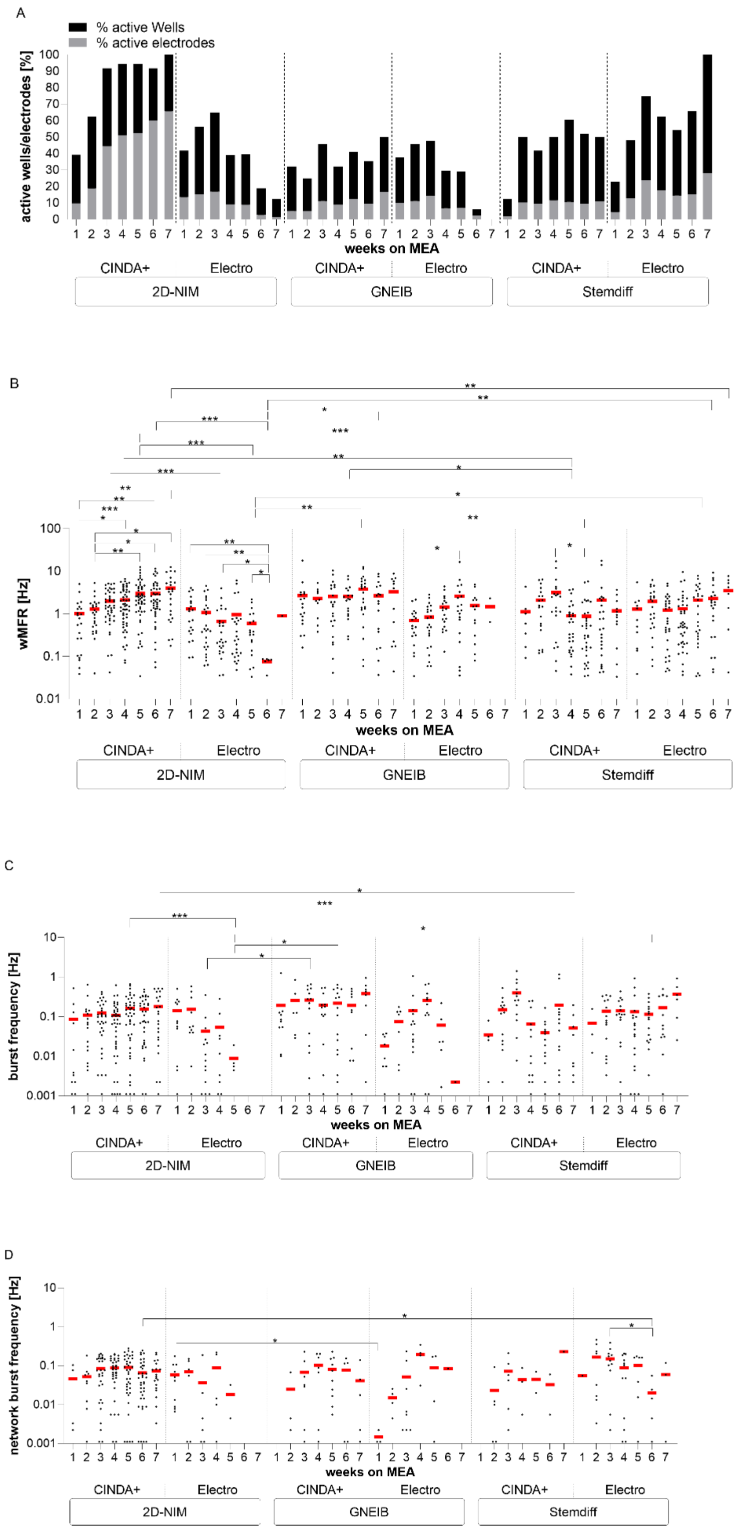


**Supplementary Figure S4.** Comparison of electrical activity of 2 weeks 3D differentiated BrainSpheres for 7 weeks on microelectrode arrays (MEA). The neuronal functionality was measured twice per week and the parameters active wells and active electrodes (A), weighted mean firing rate (wMFR, B), burst frequency (C), and network burst frequency

(D) were analyzed. Each dot represents the mean of one well containing 8 electrodes and the red bar represents the mean of all wells resulting from three independent MEA experiments each ($p \leq 0.05$)

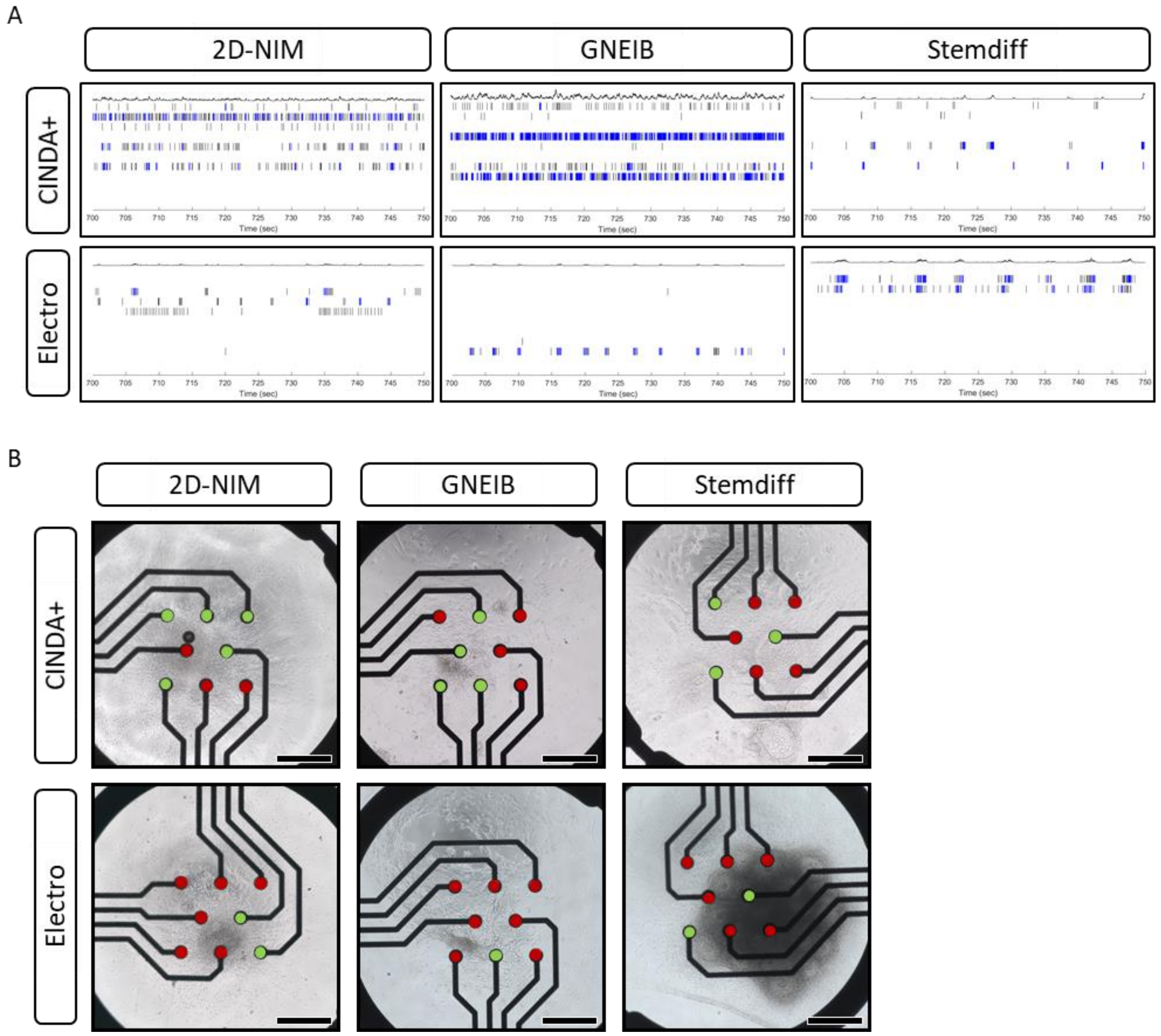


**Supplementary Figure S5.** Spike raster plots (SRP) and microscopy images of 3 weeks 3D differentiated BrainSpheres differentiated for further 3 weeks on microelectrode arrays (MEA). A) Representative 50 seconds spike raster plots of the respective networks. Spikes are shown as black bars and bursts are shown as blue bars. B) Exemplary microscopy pictures show the position of the BrainSpheres in the wells and the cells growing over the electrodes. Active Electrodes are marked with a green dot and inactive electrodes are marked with a red dot. Scale bar = 200 µm.

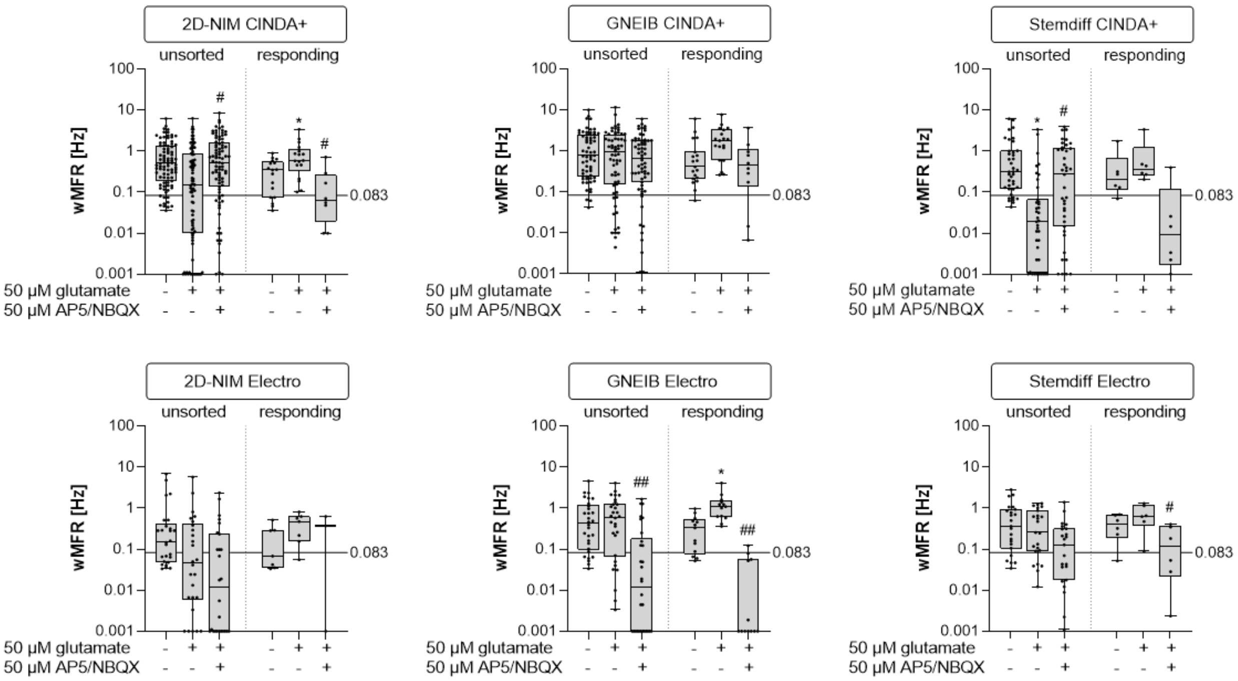


**Supplementary Figure S6.** Modification of electrical activity by acute pharmacological modulation with glutamate and ap5/nbqx. BrainSpheres were 3 weeks 3D differentiated before plated on MEA and exposed to glutamate (glu) and ap5/nbqx. Shown are the weighted mean firing rate (wMFR) of the untreated baseline measurement and after exposure to glu and ap5/nbqx of all units (unsorted) in comparison to the responding units (responding). Data are shown as box-whisker plots of three independent MEA experiments with 8 wells per condition (*p ≤ 0.05, **p ≤ 0.01, ***p ≤ 0.001). Each dot represents one unit.

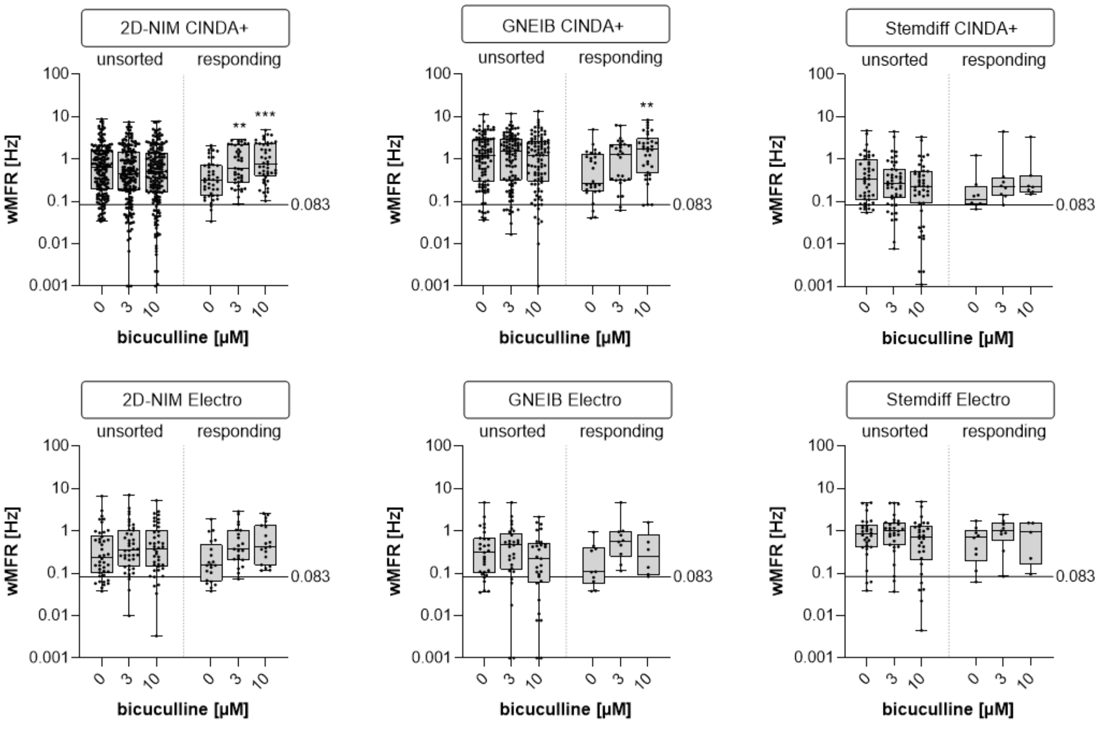

**Supplementary Figure S7.** Modification of electrical activity by acute pharmacological modulation with bicuculline. BrainSpheres were 3 weeks 3D differentiated before plated on MEA and exposed to bicuculline. Shown are the weighted mean firing rate (wMFR) of the untreated baseline measurement and after exposure to bicuculline of all units (unsorted) in comparison to the responding units (responding). Data are shown as box-whisker plots of three independent MEA experiments with 8 wells per condition (*: significant to the lowest concentration, *p ≤ 0.05, **p ≤ 0.01, ***p ≤ 0.001). Each dot represents one unit.

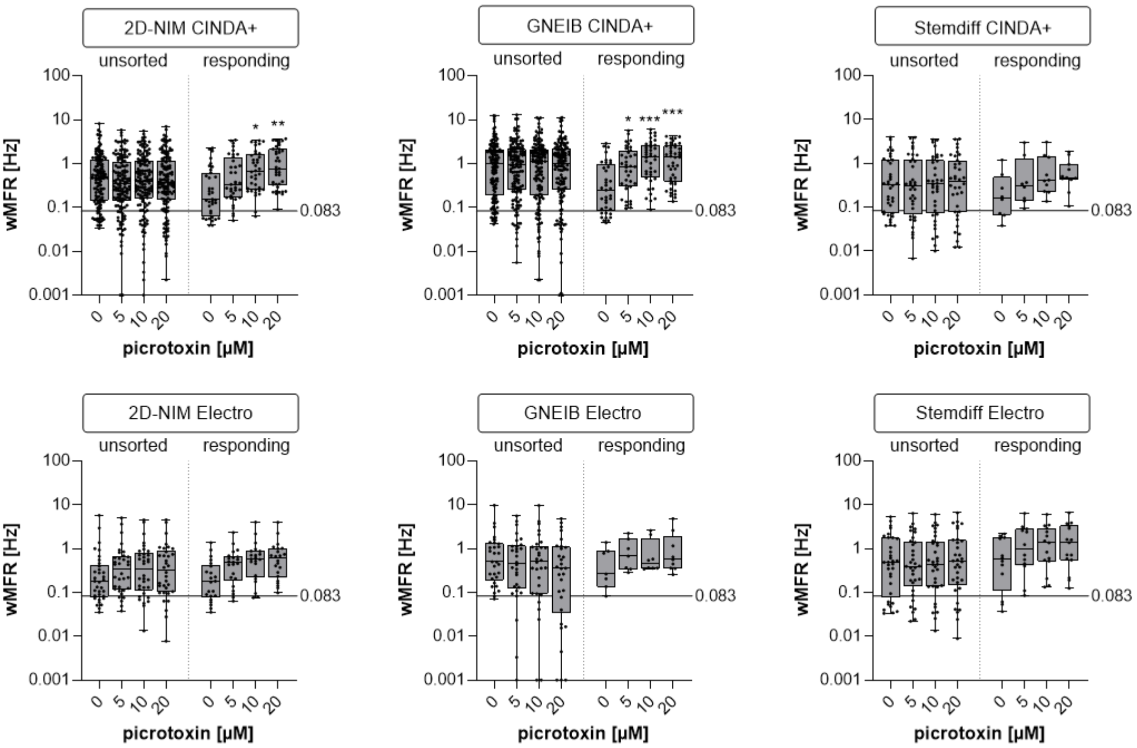


**Supplementary Figure S8.** Modification of electrical activity by acute pharmacological modulation with picrotoxin. BrainSpheres were 3 weeks 3D differentiated before plated on MEA and exposed to picrotoxin. Shown are the weighted mean firing rate (wMFR) of the untreated baseline measurement and after exposure to picrotoxin of all units (unsorted) in comparison to the responding units (responding). Data are shown as box-whisker plots of three independent MEA experiments with 8 wells per condition (*: significant to the lowest concentration, *p ≤ 0.05, **p ≤ 0.01, ***p ≤ 0.001). Each dot represents one unit.

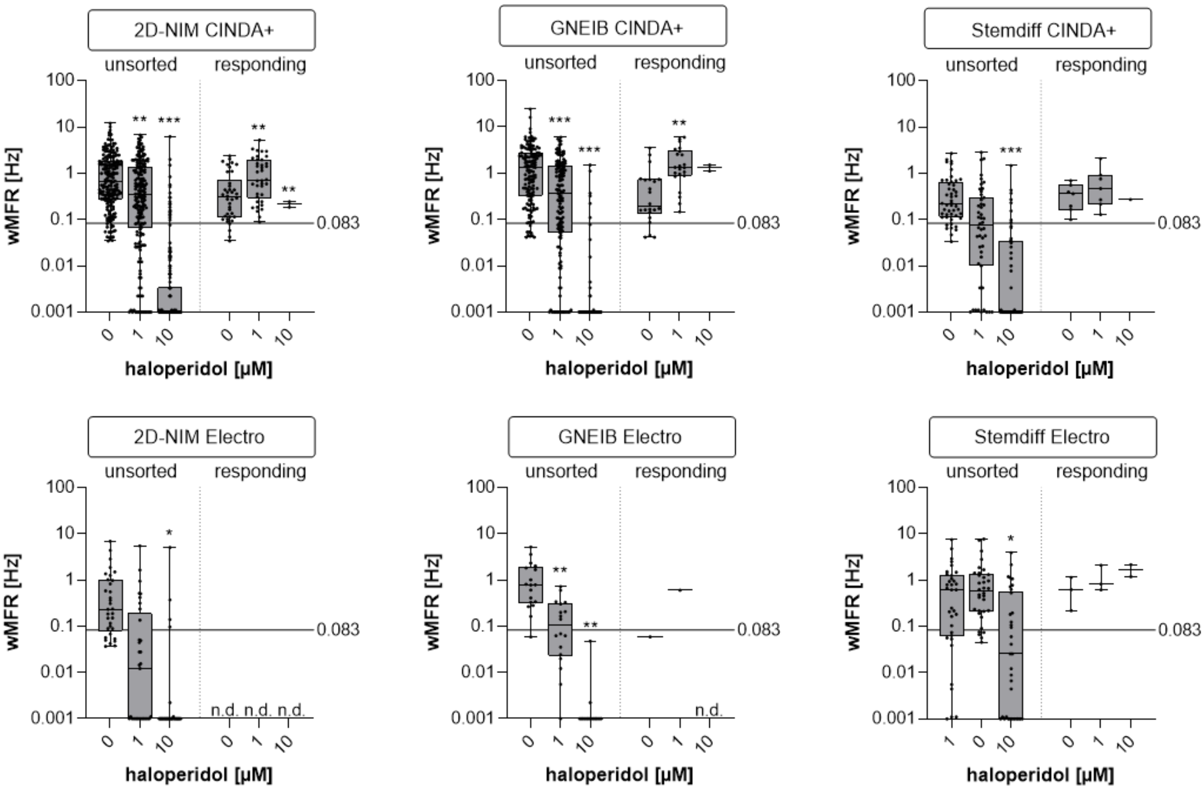


**Supplementary Figure S9.** Modification of electrical activity by acute pharmacological modulation with haloperidol. BrainSpheres were 3 weeks 3D differentiated before plated on MEAs and exposed to haloperidol. Shown are the weighted mean firing rate (wMFR) of the untreated baseline measurements and after exposure to haloperidol of all units (unsorted) in comparison to the responding units (responding). Data are shown as box-whisker plots of three independent MEA experiments with 8 wells per condition (*: significant to the lowest concentration, *p ≤ 0.05, **p ≤ 0.01, ***p ≤ 0.001). Each dot represents one unit.

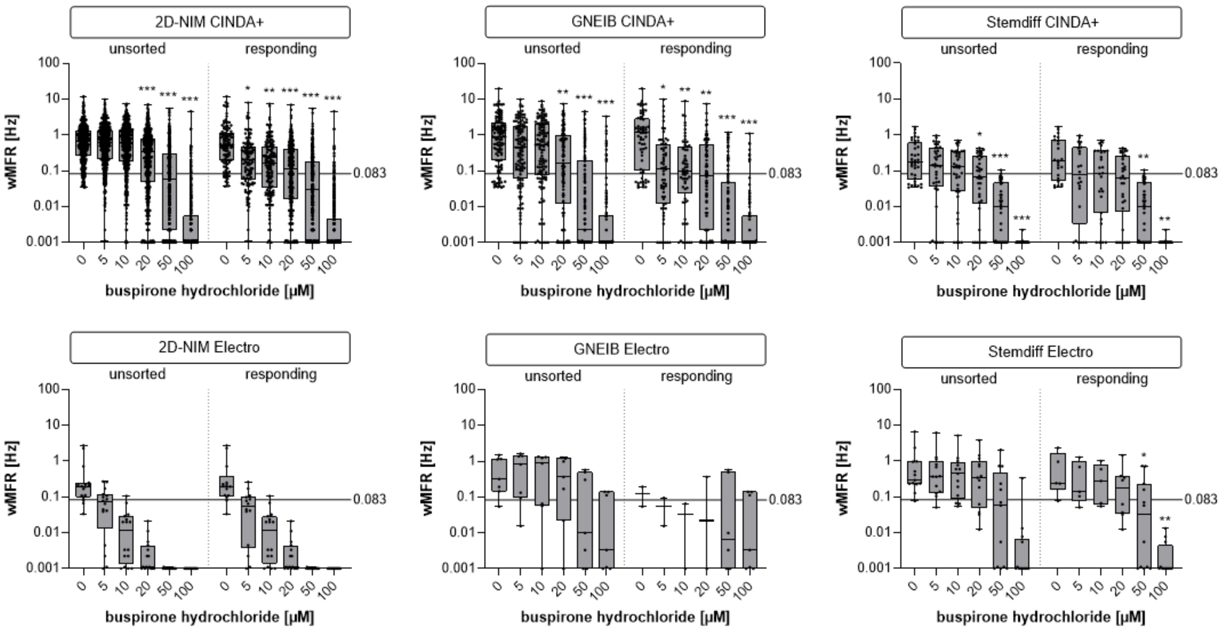

**Supplementary Figure S10.** Modification of electrical activity by acute pharmacological modulation with buspirone. BrainSpheres were 3 weeks 3D differentiated before plated on MEA and exposed to buspirone. Shown are the weighted mean firing rate (wMFR) of the untreated baseline measurement and after exposure to buspirone of all units (unsorted) in comparison to the responding units (responding). Data are shown as box-whisker plots of three independent MEA experiments with 8 wells per condition (*: significant to the lowest concentration, *p ≤ 0.05, **p ≤ 0.01, ***p ≤ 0.001). Each dot represents one unit.

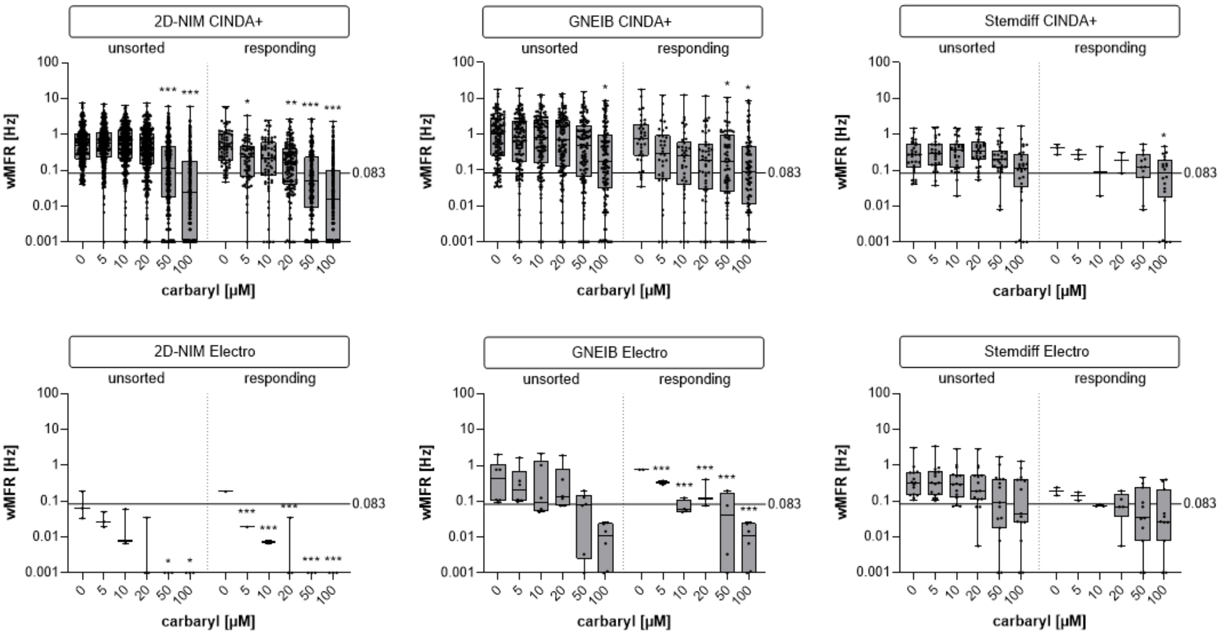


**Supplementary Figure S11.** Modification of electrical activity by acute pharmacological modulation with carbaryl. BrainSpheres were 3 weeks 3D differentiated before plated on MEAs and exposed to carbaryl. Shown are the weighted mean firing rate (wMFR) of the untreated baseline measurement and after exposure to carbaryl of all units (unsorted) in comparison to the sorted responding units (sorted). Data are shown as box-whisker plots of three independent MEA experiments with 8 wells per condition (*: significant to the lowest concentration, *p ≤ 0.05, **p ≤ 0.01, ***p ≤ 0.001). Each dot represents one unit.

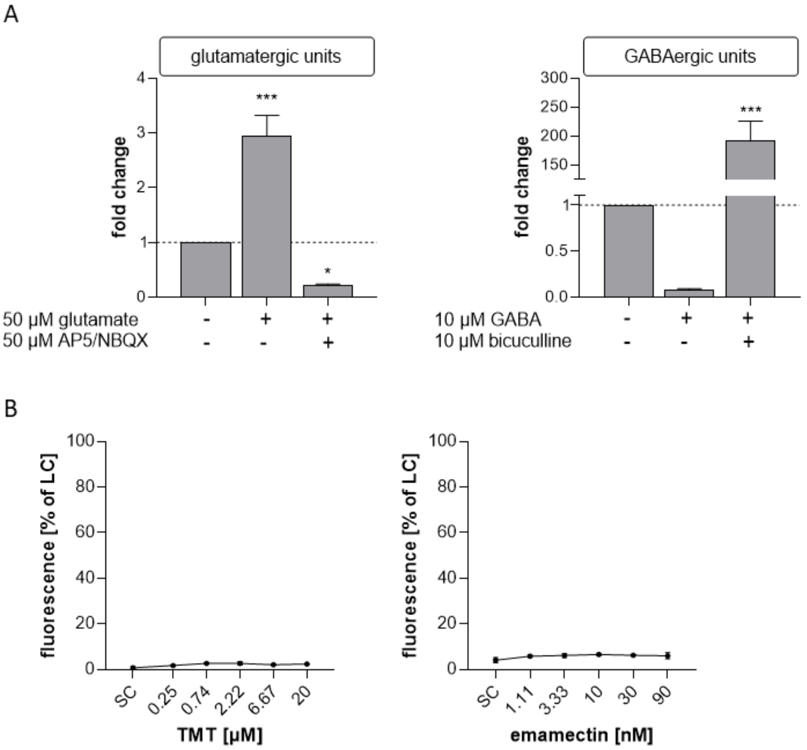


**Supplementary Figure S12.** Specification of glutamatergic and GABAergic units for acute neurotoxicity testing. A) Response of neural units to glutamate and AP5/NBQX (left) or GABA and bicuculline (right). Shown are the fold changes of the wMFR to the untreated baseline measurement. Data are represented as mean ± SEM of 24 wells with 8 electrodes each. (*: significant to the lowest concentration, *p ≤ 0.05, **p ≤ 0.01, ***p ≤ 0.001). B) Cytotoxicity assessment after acute exposure to the training compounds TMT and emamectin. Shown are the % of the LC as mean ± SEM of 3 technical replicates. TMT, trimethyltin chloride; SC, solvent control; LC, lysis control.
